# Mathematical modelling of in vitro replication dynamics for multiple highly pathogenic avian influenza clade 2.3.4.4 viruses in chicken and duck cells

**DOI:** 10.1101/2025.04.22.649950

**Authors:** Luca Bordes, Peter Hobbelen, Elena Blokker, Wim H. M. van der Poel, Catherine A. A. Beauchemin, Nancy Beerens, José L. Gonzales

## Abstract

The introduction, spread, and subsequent detection of highly pathogenic avian influenza viruses (HPAIV) in poultry are influenced by the virus replication fitness, transmission fitness, and virulence. Quantifying these fitness parameters is important for implementing surveillance and control measures for poultry. This study employs an avian *in vitro* model using primary chicken (CEF) and duck (DEF) embryo fibroblasts, to identify cellular correlates of fitness for a reference panel of eight dominant HPAI clade 2.3.4.4 virus genotypes (H5N1, H5N8, and H5N6) isolated between 2014 and 2024. Because HPAI viruses are pantropic in poultry, the use of primary fibroblasts could provide a robust systemic model to assess virus fitness in different poultry species. By integrating longitudinal infectious virus titres and cytopathogenicity data with a mathematical model, which delineates cell populations into susceptible, latent, infectious, and dead compartments, we disentangled “traditional” virological parameters such as peak virus titre and time to 50% cell death, from mechanistic parameters, including the infecting time (*t*_*inf*_), generation time (*t*_*gen*_) and basic reproduction number (R_0_). Our results confirm host-specific fitness characteristics consistent with *in vivo* observations. This study underscores the potential of integrating avian *in vitro* models with mathematical modeling to provide high resolution characterization of genetically similar viruses building towards rapid risk assessments of novel HPAI viruses.

**Importance:** HPAI outbreaks in poultry have resulted in culling of millions of animals over the past decade. The introduction and spread of HPAI in poultry are driven by the ability of the virus to replicate, transmit, and cause disease, which varies among different poultry species. These viral fitness traits can change quickly through mutations or reassortment, leading to new variants with an altered risk of introduction and transmission. In this study, we integrated an *in vitro* and mathematical model to assess viral fitness of genetically similar viruses and the risk they represent for two important poultry species. This approach offers a rapid and scalable alternative to traditional animal models, aligning with the 3R’s principle and supporting rapid assessment of the viral fitness of emerging virus variants for poultry. This *approach* can be easily expanded to other species including humans and is suitable to assess the viral fitness of all viral pathogens.

## Introduction

Highly pathogenic avian influenza (HPAI) H5-viruses are circulating in wild birds and poultry, causing mass mortality in poultry and occasionally in wild birds [1, 2]. Repeated introduction of HPAI H5-viruses from wild birds to poultry have been observed in regions where the infection is not enzootic in poultry, resulting in significant losses to the poultry sector [1]. HPAI virus evolution is characterized by reassortment and mutations resulting in the emergence of multiple clades and genotypes [3]. In Europe HPAI H5-viruses of clade 2.3.4.4 have caused multiple epizootics since 2014 but virulence and severity for different host species as well as the number of species affected during these epizootics differed between epizootic years [4]. Further genetic diversification occurred during the emergence of the H5N1 virus in Europe in 2020 and an additional nomenclature system was implemented. Letters are used to further distinguish between H5N1 genotypes based on all eight viral genome segments, unlike the clade system which is based solely on the Hemagglutinin (HA) segment. Some HPAI viruses are able to efficiently transmit (reproduction number R > 1) without causing apparent clinical signs in poultry such as ducks and geese. Infections in these species can thus remain undetected, increasing the risk of virus spread to other farms, wild birds and mammals [5-8]. The risk of introduction and subsequent detection of introduction in poultry depends among other things on the replication fitness (ability to replicate within a host), transmission fitness (ability to infect other animals) and virulence (ability to cause disease) in wild birds and poultry. Determining these fitness parameters could offer vital insights for evaluating the risk of a particular genotype in causing an outbreak across various poultry species.

Replication fitness of a virus in a specific host species and transmission fitness are altered by changes in replication efficiency in certain tissues and virus replication-induced tissue damage [9], which subsequently affect peak shedding and shedding duration (replication dynamics). Animal experiments are primarily used to evaluate the replication fitness and/or dynamics, transmission fitness and virulence of a virus for specific host species [6, 7, 10-14]. This limits the number of HPAI strains (genotypes) that can be evaluated, are slow to implement during emerging outbreaks, are costly and time consuming. Furthermore, the inoculation route, dosage, age and model species often differs between animal experiments which may lead to contradicting conclusions [15].

Replication dynamics could be evaluated without animal experiments using *in vitro* models, which allow for better control over experimental variables, the ability to compare a broader range of viruses across different species (low costs), and the potential for real-time risk assessment of emerging AIV variants (fast)[16-18]. Replication dynamics of influenza viruses on cell cultures have three distinct phases; a latent period, an exponential growth phase during which cells are rapidly destroyed by the virus and an exponential decay phase [19]. The dynamics of the different phases could differ between influenza viruses depending on their genetic composition. Translation of *in vitro* replication dynamics to *in vivo* replication dynamics can be improved by the use of mathematical models that quantitatively characterize these dynamics by estimating replication parameters at the individual cell level, similar to epidemiological parameters at the individual host level, such as the infecting time (time from the start of virus production by a cell until the first secondary infection), generation time (the average time from infection until a secondary infection, including the latent period) and R_0_ (average number of cells that become infected by one infectious cell in a susceptible population) [19-21]. Depending on the sampling method, the number of samples that can be collected and analyzed over time is usually limited and requires strain specific optimization to capture the replication dynamics. Mathematical models can, at least in part, circumvent these shortcomings by interpolating dynamics between the snap-shot experimental measurements, providing a continuous view of the dynamics. Mathematical models have been applied to characterize human influenza viruses in immortalized cell lines like Madin-Darby Canine Kidney (MDCK) cells and human lung cells (A549) [19, 20, 22]. However, no prior studies have explored mathematical models for characterizing HPAI in poultry cells.

We have selected a reference panel of eight dominant HPAI H5 clade 2.3.4.4 genotypes, isolated during the 2014–2024 epizootics, against which novel genotypes can be benchmarked. An avian *in vitro* model was established to characterize the viral replication dynamics of the reference panel, which was subsequently integrated with a mathematical model for in depth analysis. The mathematical model was developed using previously published data from our laboratory [23] to identify optimal sampling points for contemporary HPAI viruses without compromising the accuracy of the estimates. This was done to increase the number of viruses that could be characterized whilst reducing laboratory effort (“cost-effective”). The virus replication dynamics are described using a compartmental mathematical model that divides the cell population into susceptible, latent, infectious and dead compartments. The mathematical model was then used to estimate infection time, generation time, R_0_, and other parameters for the eight HPAI H5 viruses included in the panel, using primary chicken and duck fibroblasts.

## Methods

### Virus isolation, propagation and sequencing

All experiments involving HPAI viruses were strictly performed under biosafety level 3 (BSL-3) regime, in accordance with national and institutional biosafety requirements.

Virus selection was based on the dominant HPAI genotypes during the 2014-2024 epizootics in the Netherlands. Index cases from seven Dutch HPAI viruses were isolated from poultry (H5N8-2014 chicken, H5N8-2016 duck, H5N6-2017 duck, H5N8-2020 chicken, H5N1-2020-C chicken, H5N1-2021-AC chicken, H5N1-2021-AB chicken) and were passaged twice on 9-to 11-day-old specific pathogen free (SPF) embryonated chicken eggs (E2) (Royal GD, Deventer, the Netherlands). One wild bird isolate (H5N1-2022-BB gull) was passaged once on 9-to 11-day-old SPF embryonated chicken eggs (E1). The E2 passage for poultry isolates and E1 passage for the wild bird isolate was sequenced using full genome sequencing as described previously [24]. In addition, RNA isolated directly from the swab material was sequenced. In short, virus RNA was isolated using the High Pure Viral RNA kit (Roche, Basel, Switzerland), amplified using universal eight segment primers and directly sequenced at high coverage (average > 1000 per nucleotide position) using Illumina DNA Prep and Illumina MiSeq 150PE sequencing. The CLC Genomics Workbench extension ViralProfiler Workflow (Qiagen, Germany) was used to map the reads to a reference set of genomes. Consensus sequences were generated and uploaded to GISAID. Only minor differences between the sequences of the swabs and egg passages were detected, and no mutations on positions associated with virulence changes were identified. Differences detected between the sequences of the swab materials, and the egg passage are listed in Table 1. Host shift adaptations were identified using the FluMutGUI version 3.1.1 and FluMutDB 6.2 (released on 15-07-2024) [25]. Similarity between the selected HPAI viruses based on each genome segment was determined with a nucleotide sequence identity threshold value of 1.5%.

**Table 1:** Genetic background and general information of HPAI H5-viruses in this study.

| Abbreviation | Virus isolate | EPI number | Clade 2.3.4.4 | Poultry species virus was isolated from | Differences between swab isolate and second egg passage |
| --- | --- | --- | --- | --- | --- |
| H5N8-2014 | A/chicken/Netherlands/14015531/2014 | EPI_ISL_18600237 | Group c | Chicken | PB2-R574K, PA-F451S, NA-A332V |
| H5N8-2016 | A/duck/Netherlands/16014829-001005/2016 | EPI_ISL_529179 | Group b | Duck | PB2-V344A |
| H5N6-2017 | A/Duck/Netherlands/17017236-001-005/2017 | EPI_ISL_18600770 | Group b | Duck | PB2-I30L |
| H5N8-2020 | A/chicken/Netherlands/20016597-026030/2020 | EPI_ISL_8650948 | Group b | Chicken | - |
| H5N1-2020-C | A/Chicken/Netherlands/20019879-001005/2020 | EPI_ISL_17791407 | Group b – genogroup C | Chicken | - |
| H5N1-2021-AB | A/chicken/Netherlands/21037287-006010/2021 | EPI_ISL_9856775 | Group b – genogroup AB | Chicken | - |
| H5N1-2021-AC | A/chicken/Netherlands/21038165-006010/2021 | EPI_ISL_13360259 | Group b – genogroup AC | Chicken | - |
| H5N1-2022-BB | A/lesser black-backed gull/Netherlands/22012469-002/2022 | EPI_ISL_19201535 | Group b – genogroup BB | Gull | - |

### Virus titration

MDCK cells obtained from Philips-Duphar (Weesp, the Netherlands) were cultured at 37°C, 5% CO_2_ using cell culture medium consisting of Dulbecco’s Modified Eagle Medium GlutaMAX (DMEM)(Thermo Fisher Scientific) supplemented with 5% fetal calf serum (Capricorn scientific, Germany) and 0.1% penicillin-streptomycin (Thermo Fisher Scientific). Cells were passaged when confluent using 0.05% Trypsin-EDTA (Thermo Fischer Scientific). The median tissue culture infective dose (TCID_50_) of the isolated viruses was determined using 10-fold serial dilutions, 8 wells/dilution, as described earlier [26]. For each isolated virus, TCID_50_ was measured in triplicate and repeated on a different day (n=6).

### Primary embryonic fibroblast isolation and culture

Chicken embryo fibroblasts (CEF) were harvested from 11-day-old SPF embryonated chicken eggs (Royal GD) and duck embryo fibroblasts (DEF) were harvested from 11-day-old Pekin duck embryos obtained from a commercial breeding farm. Three Pekin duck egg yolks from each breeding pair were tested to exclude the presence of maternal antibodies using the IDEXX Influenza A virus Ab ELISA according to the manufacturer’s instructions. Primary embryo fibroblasts were harvested by first removing the embryo head, limbs and internal organs. The remaining tissue was minced, washed three times in PBS and incubated at 37°C for 5 minutes in 0.25% Trypsin (Thermo Fisher Scientific). Supernatant was collected and trypsinization of the remaining tissue was repeated twice with fresh 0.25% Trypsin. Collected supernatant was diluted in cell culture medium and strained over a 100 µm pore size filter. The cells were spun down for 5 minutes at 300 g and resuspended in primary fibroblast medium consisting of medium 199 with Earle’s salts supplemented with 3.6% new born calf serum (NBCS, 0.12% sodium bicarbonate, 2 mM l-glutamine, 0.1× MEM Vitamin solution, 1% penicillin-streptomycin and 0.5% gentamicin (all Thermo Fisher Scientific). After one day, cells were washed twice with PBS, harvested using 0.05% Trypsin-EDTA (Thermo Fischer Scientific), resuspended in freezing medium consisting of medium 199 supplemented with 30% FCS and 10% DMSO (Sigma-Aldrich) and frozen in liquid nitrogen for later use.

### Cytopathogenic effect and replication curves

CEF and DEF cells were thawed, resuspended in fibroblast medium and a total of 3.5 x 10^5^ cells per well were seeded in a 24-well plate for virus growth curves or in an electronic tissue culture plate (E-plate)(ACEA Biosciences) for measuring the cytopathogenic effect. After 24 hours, cells were gently washed using PBS with calcium and magnesium (Thermo Fischer Scientific). After 2 hours of recovery time the cells were infected with a multiplicity of infection (MOI) of 0.001 in serum-free fibroblast medium. Electrical impedance, displayed as cell index, was measured on the E-plates every 30 minutes up to 48 hours post infection (hpi) and the cell index values were normalized at 2 hpi. For the virus growth curves, the supernatant from the entire well was collected after 10 hpi, 18 hpi, 26 hpi and 34 hpi and stored at -80°C until used for virus titration on MDCK cells as described above (one TCID_50_ in eight-fold for each time point). After supernatant harvesting, the cells were washed twice with PBS, dried and frozen at -20°C. Cells were subsequently fixed with 10% neutral buffered formalin and stained using 1 µg/ml 4C,6-Diamidine-2C-phenylindole dihydrochloride (DAPI)(Roche). Both the cytopathogenic effect and replication curves were measured two times in duplicate for each virus and cell type combination (n = 4). Peak titer and the time until 50% reduction of the cell index was extracted for each replicate and the difference between CEF and DEF cells were compared using the Wilcoxon test.

### Mathematical model

#### Model structure

A mechanistic ordinary differential equation model was used to describe the population dynamics of HPAI viruses on CEF and DEF cells. The mathematical model divides the cell population in susceptible, latent, infectious and dead compartments. Assumptions of the mathematical model include, no cell divisions during the experiment, infection rate was proportional to both the number of susceptible cells and the number of infectious virus particles, and infectious cells produce new infectious virus particles at a constant rate until they die. A gamma distribution was used for the sojourn time of cells in the latent and infectious stages. Similar model structures were used in previous modelling studies on the in vitro dynamics of human influenza A viruses on immortalized cells [19, 22, 27]. The model was extended by including the loss of infectious virus particles due to entry into target cells during the infection process as previously described [20]. These assumptions lead to the following system of ordinary differential equations with state variables and parameters defined in Table S1 and Table S2:

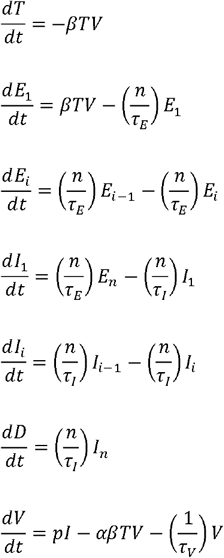

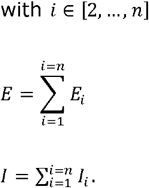

#### Parameter estimation

The initial values of state variables were derived from the experimental design. All other model parameters were estimated as described below based on the infectious virus titre and cell index data. For this purpose, cell index values were converted to fractions of the initial number of cells that were still alive at time *t*, from the cell index (CI) data by the following equation:

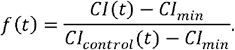

In this equation, *CI*(*t*) denotes the cell index at time *t* for the inoculated well and *CI*_*min*_ represents the cell index at the end of the experiment when all cells were assumed to have died. Variable *Cl*_*control*_(*t*) denotes the value for the corresponding control well at time *t*. Since the number of cell index data points was much higher than the number of infectious virus titre data points, the data was weighted when inferring parameters. We used an Approximate Bayesian Computation algorithm based on sequential Monte Carlo (ABC-SMC) for parameter inference [28, 29]. This algorithm quantifies the goodness-of-fit using a distance measure. In case there are different types of data available, separate distance measures can be used for each data type (intersection approach) or a single distance measure can be specified that combines the goodness of fit for different data types (union approach)[30]. We used this last approach and formulated a single distance measure as the sum of squared residuals with each data type weighed by the variation in the data as well as the number of data points:

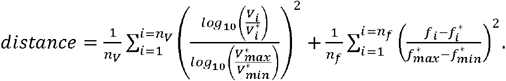

In this equation, symbols *n*_v_ and *n*_*v*_ represent the number of datapoints for the virus titre and for the fraction of living cells. Symbols *V*_*1*_ and 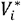 represent the predicted and observed virus titre for datapoint *i*. Similarly, *f*_*i*_ and 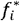 represent the predicted and observed fraction of living cells for datapoint *i*. Symbols 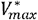 and 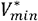 represent the highest and lowest observed virus titre in the data. Similarly, 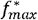 and 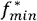 represent the highest and lowest observed fraction of living cells in the data.

Infectious virus titre data in the distance measure were expressed on a log_10_ scale in order to prevent bias towards datapoints with higher virus concentrations. In short, the ABC-SMC algorithm described by [28] samples particles (a set of model parameters) from the prior distributions and keeps the ones whose distance measure satisfies the tolerance criterion. It subsequently starts a loop producing intermediate particle populations. In this loop, each particle population is generated from the previous population using weighted sampling and perturbation according to a perturbation kernel. Perturbed particles were accepted if their probability under the prior distributions was >0 and the distance measure met the tolerance criterion. The last particle population is used to generate the posterior distributions of model parameters.

The size of the particle populations should be sufficient to generate stable posterior distributions of model parameters. To set the size of the particle populations in the ABC-SMC algorithm, we evaluated the posterior distributions of model parameters for a range of population sizes when using data for strain H5N1-AB on CEF cells (Figure S3). Based on this analysis, we required each population to contain 1000 accepted particles. We used two types of perturbation kernels (*K*). For a given continuous parameter *i*, we used:

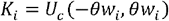

with *U*_*c*_ representing the continuous uniform distribution. The boundaries of this distribution were set by multiplying the width of the previous marginal distribution for parameter *i*(*w*_*1*_) with constant *θ* [31]. The value of *θ* should be just high enough to prevent the algorithm from becoming stuck in local modes without unnecessarily increasing the rejection rate of proposed particles and reducing the efficiency. To find this optimum value, the value of *θ* was increased in a number of steps until the posterior distributions of model parameters seemed stable when using data for strain H5N1-AB on CEF cells (Figure S4). Based on this analysis, *θ* was set to 0.1. A similar value for *θ* was previously used [31, 32]. Discrete variables were derived as:

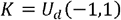

with *U*_*d*_ representing the discrete uniform distribution.

As tolerance criterion, the distance measure of a proposed particle should be less than the median distance measure from the previous particle population. The loop creating new particle populations ended when the median distance measure of the last particle population differed by less than 1% from the median distance measure of the previous particle population. Convergence of the posterior distributions was visually monitored by plotting the 95% interquantile range of parameters for subsequent particle populations (Section 6 of the Supplementary Information). To set a tolerance criterion for generating the first intermediate particle population from the prior distributions, a random sample from the prior distributions was taken to perform model simulations and obtain the distance measure for 1000 prior particles. The median distance measure was then used as tolerance criterion for generating the first intermediate population. Table S2 shows the prior distributions for each calibrated model parameter. The identifiability [33] of the calibrated model parameters was assessed from the credible interval of model parameters and the degree of correlation between the model parameters (Section 6 of the Supplementary Information).

#### Calculation of mechanistic traits

The infecting time is defined as the time from the start of the infectious period of a cell until the first secondary infection by this cell in an otherwise susceptible population and was derived as [34]:

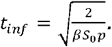

The basic reproductive number (*R*_0_) is defined as the average number of cells that become infected by one infectious cell in an otherwise susceptible population [21]. The *R*_0_ was calculated as described previously [21] adjusting the equation to account for the loss of virions due to the infection of cells [20], resulting in the following equation:

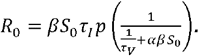

The generation time is defined as the average time interval from the infection of one cell in an otherwise susceptible population until a new infection caused by the initially infected cell and was calculated as [20]:

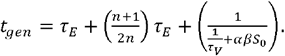

## Data availability

Whole genome sequencing data has been deposited in the GISAID database and can be accessed via the GISAID Accession IDs (Table 1). The mathematical models, and corresponding code developed in this study are publicly available on GitHub at **[To be added]**.

## Results

### Genetic analysis and potential host shift adaptations

The HPAI viruses H5N8-2016 and H5N6-2017 were isolated from Pekin ducks, HPAI H5N8-2014, H5N1-2020-C, H5N1-2021-AB, H5N1-2021-AC were isolated from chickens and HPAI H5N1-2022-BB was isolated from a Gull (Table 1). The HPAI H5N8-2014 virus belongs to clade 2.3.4.4c while the remaining seven HPAI H5 viruses belong to clade 2.3.4.4b. Virus sequences were screened for previously identified virulence factors known to influence replication, virulence or binding of host proteins using FluMut (Table S3)[25]. Only one marker, M105V on the nucleoprotein (NP), was shown to increase virulence in chickens and was present on all investigated viruses except in the H5N8-2014 and H5N1-2021-AB sequences. Other mutations included increased or decreased replication on mammalian cells or virulence in mammals. Disruption of the second sialic acid binding site was identified for the H5N8-2014 and H5N1-2022-BB virus. Genome segment clustering revealed similarities between the H5N8-2016 and H5N6-2017 viruses, as well as among the four H5N1 viruses (Table S4). Additionally, the hemagglutinin (HA) and matrix protein (MP) of the H5N8-2020 virus showed similarity to all the selected H5N1 viruses, while the H5N8-2014 virus did not share any similar genome segments with the other viruses.

### Assessing replication dynamics without mathematical modelling

All eight HPAI viruses replicated efficiently on CEF and DEF cells and caused extensive cytopathogenicity resulting in complete loss of the cell layer within 48 hpi (Figure 1 & Figure S1). Peak virus replication was 10-100-fold lower on CEF cells than on DEF cells and the peak virus replication was slightly higher for the H5N1 viruses compared to the H5N8 viruses on CEF cells. No significant difference in peak virus replication could be measured for the H5N8-2014 virus on CEF compared to DEF cells. Interestingly, the difference in median peak titres observed between DEF and CEF cells for the two viruses isolated from ducks (H5N8-2016 and H5N6-2017) was wider (2 and 1.56 log_10_ TCID50/ml respectively) than the difference between the median peak titres (≤ log_10_ 1.1 TCID50/ml) observed for the viruses isolated from chickens. The time to 50% cell death measured by the cell index was longer on DEF cells than on CEF cells for the H5N8-2014 and four H5N1 viruses while similar or shorter time to 50% cell death was measured for the H5N8-2020, H5N8-2016, H5N6-2017 viruses. With the latter two being the viruses isolated from ducks..

**Figure 1:**
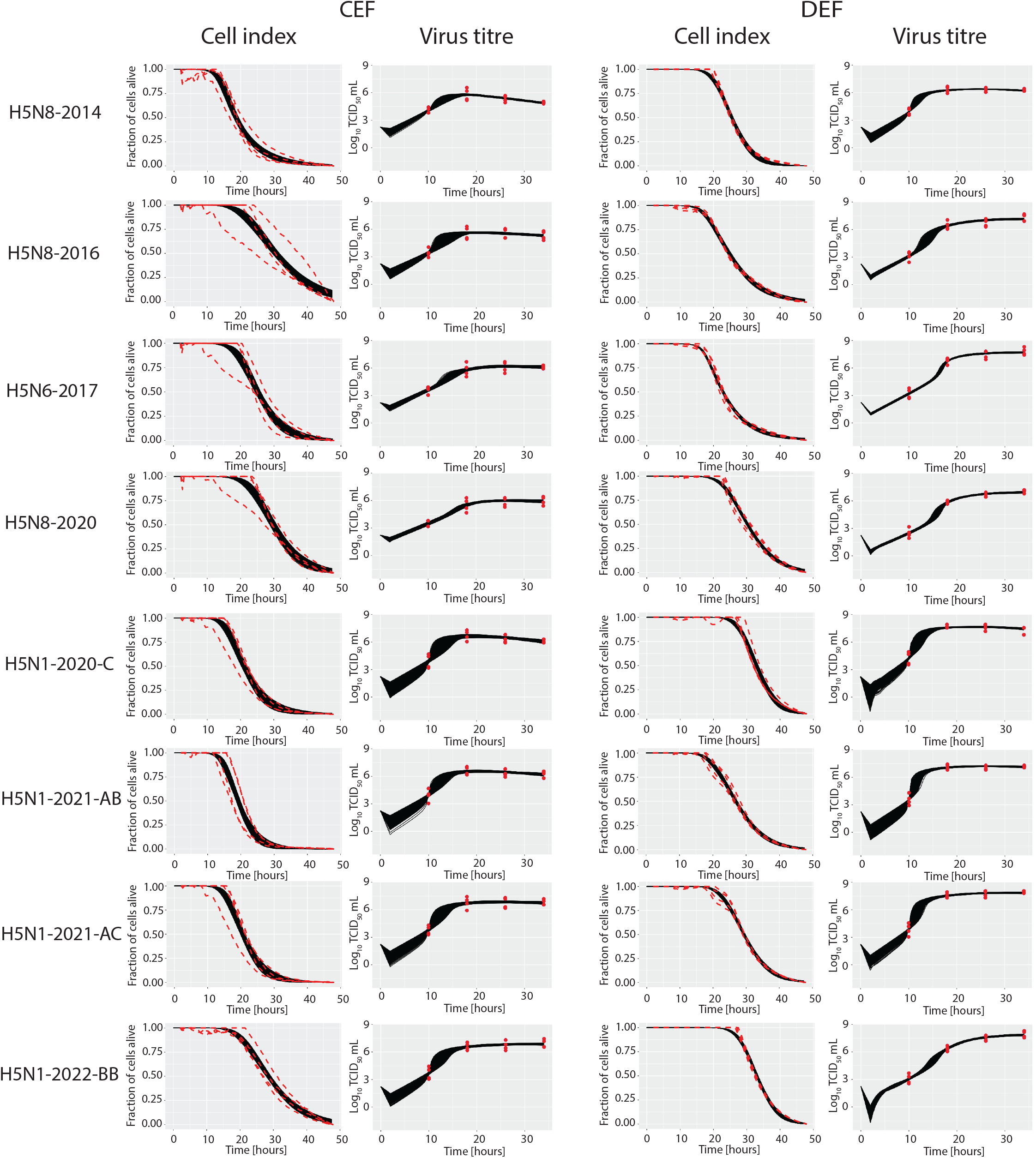
A) Peak virus replication and B) time to 50% cell death measured by the cell index calculated without mathematical modelling. Complete virus growth curves and cell index values over time are displayed in Figure 2. The extremes are displayed by the whiskers and all data points are depicted by blue or red dots. p < 0.05 is indicated by *.

### Model fit and identifiability of parameters

The variability in the cell index data was slightly less on DEF cells than on CEF cells but no apparent differences were observed between viruses (Figure 2). Cell index data in DEF cells reached zero well before the end of the experiment while on CEF cells the H5N8-2016, H5N8-2020 and H5N1-2022-BB viruses reached zero shortly before termination of the experiment. Images of DAPI-stained CEF cells from the virus titre experiment showed that all cells had died before the end of the experiment (Figure S1). This observation is in agreement with the assumption that was made for calculating the fraction of cells alive from the cell index data (see parameter estimation section). The credible interval of parameters was smaller when the peak virus titre was located past 18 hpi suggesting earlier measurements may further improve the model fit (for example H5N6-2017 on DEF cells). Overall, the model fitted well to the cell index and virus titre data.

**Figure 2:**
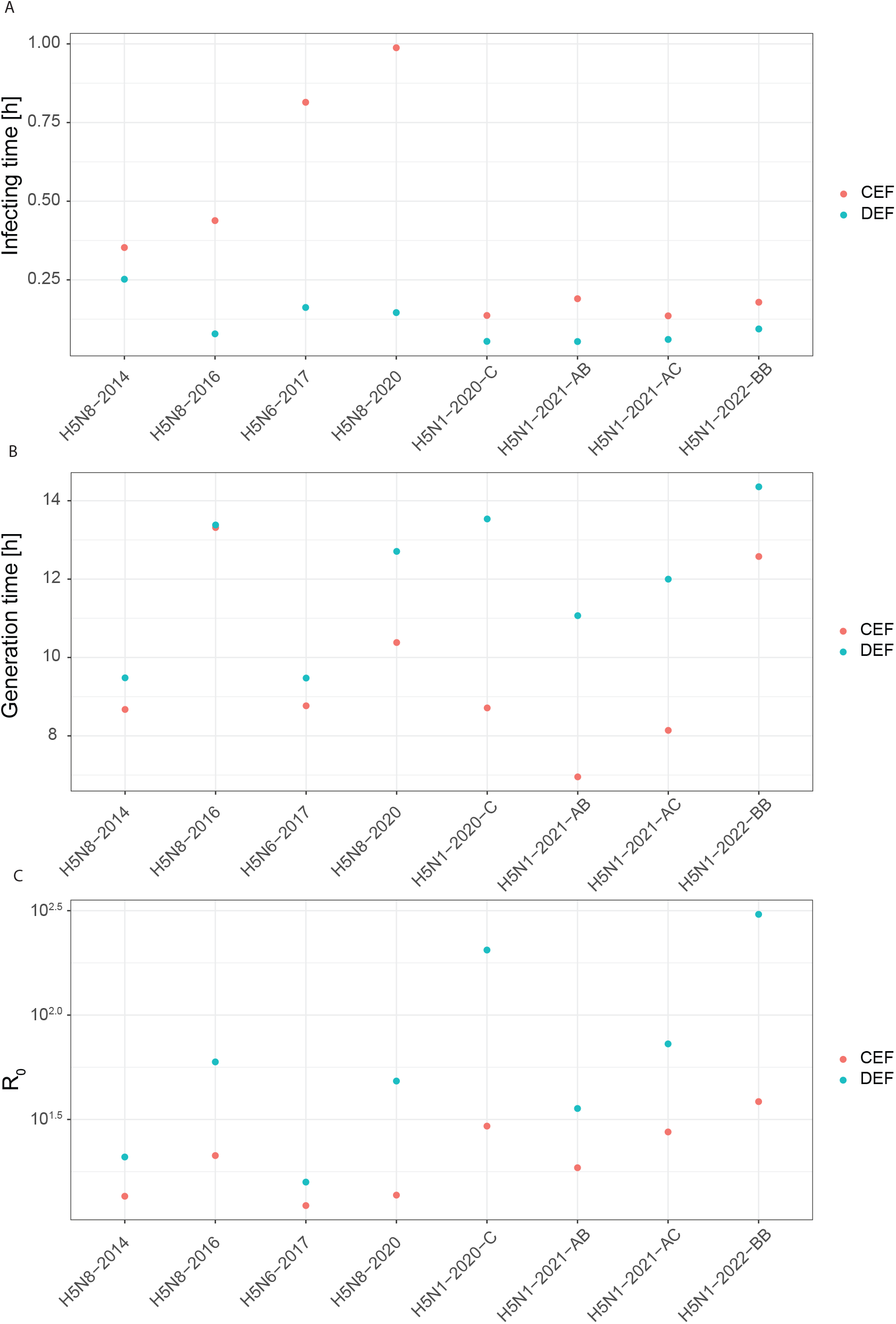
Model fit to cell index and infectious virus titre data on CEF and DEF cells. Dashed red lines and red dots indicate the raw data while the black lines indicate the model predictions.

We evaluated the identifiability of parameters by visually comparing the spread and shape of the posterior and prior distribution and by determining the correlation between parameter values (Section 6 of the Supplementary Information). In most cases, the estimated posterior distributions of model parameters were unimodal and more clustered compared to the uniform prior distributions. Exceptions are the clear bimodal shapes for the obtained distributions of the latent and infectious period for strain H5N8-2014 on CEF cells (Figure S36). The upper boundaries of the posterior distributions for the production rate of infectious virus particles by cells and for the lifetime of infectious virus particles were sometimes close or equal to the upper boundaries of the corresponding prior distributions (3 and 7 virus-host cell combinations, respectively). This suggests that there is not enough information in the data to identify the distributions for these parameters. For the lifetime of infectious virus particles (*τ*_V_), this is caused by the absence of a decrease in the measured virus titre even after most cells have died. These exceptions aside, the vast majority of parameters were identifiable based on the shape and spread of the posterior distributions compared to the prior distributions. The degree of correlation between model parameters was however almost always highly significant for any pair of parameters (including mechanistic traits) and any virus-host cell combination. This suggests that posterior distributions for individual parameters could have been narrower if more information would have been available in the data.

### Model output of mechanistic traits

Summaries of the posterior distributions of model parameters are shown in Figure S2. Derived median estimates of the mechanistic traits for the different combinations of virus strain and cell type are shown in Figure 3. The infecting time, generation time and basic reproductive number are mechanistic traits linked to viral replication dynamics in susceptible cell populations, and were compared across virus strains and between species specific cell types (Figure 3). The estimated infecting time for all virus strains tended to be shorter on DEF cells than those observed for CEF cells. In DEF cells, the estimated infecting time appeared to be similar across all virus strains, while in CEF cells estimated infecting times were notable high, relative to the other virus strains, for H5N8-2016, H5N6-2017 and H5N8-2020 (Figure 3A).

**Figure 3.**
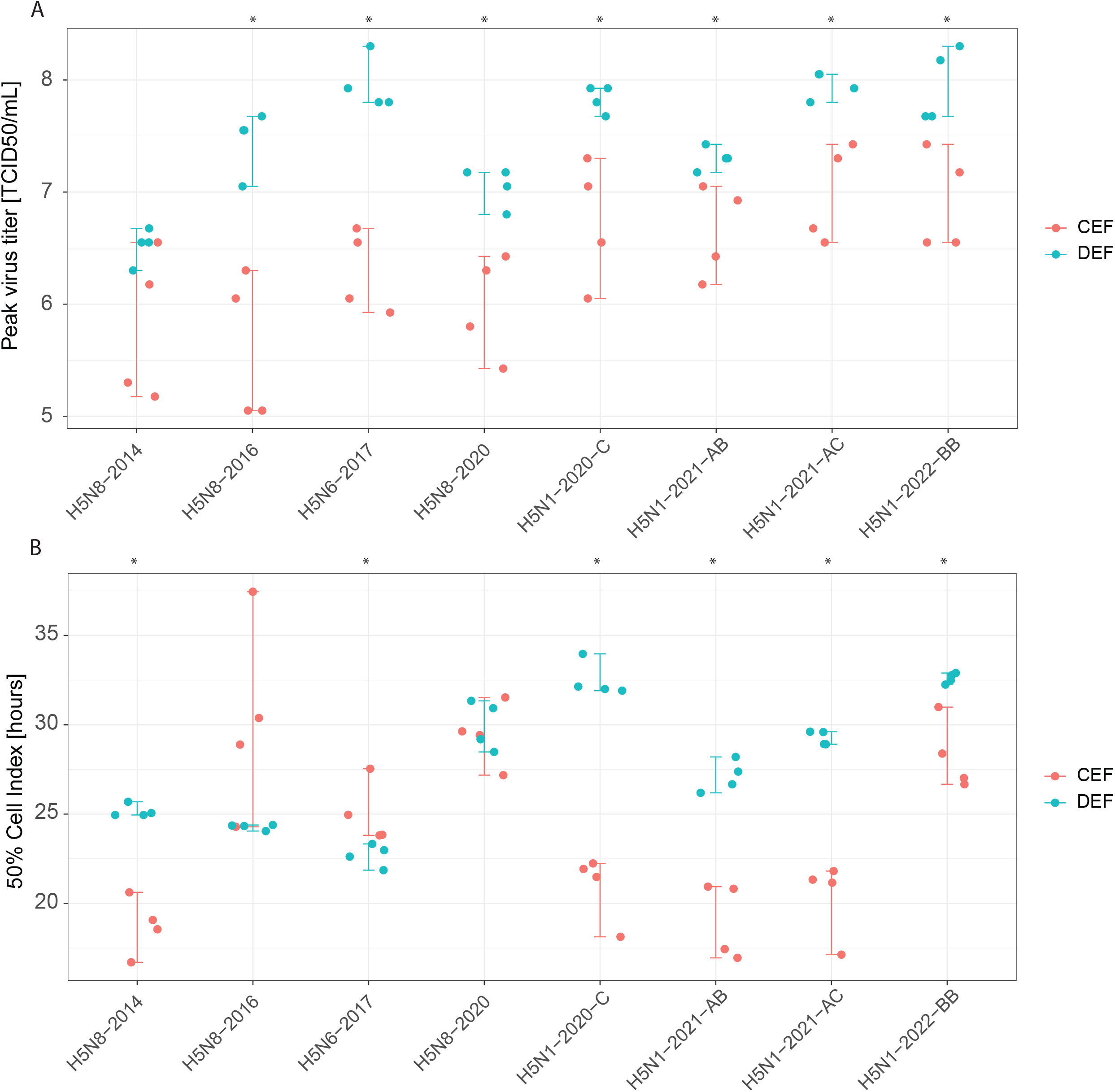
Mechanistic traits identified using mathematical modelling. The medians are displayed by the blue and red dots. A) Time to first secondary infection from start of infectious period *t*_*inf*_, B) Average time to secondary infection including latent period *t*_*gen*_, C) Number of cells infected by one infected cell R_0_.

The estimated generation time of the more recent H5 viruses (H5N8-2020 and the four H5N1 viruses) appeared to be longer on DEF cells than on CEF cells while the earlier H5 viruses showed similar generation times (Figure 3B).

The estimated R_0_ values in CEF cells appeared comparable across most virus strains, and were generally lower than those observed in DEF cells (Figure 3C). In contrast, a broader range of estimated R_0_ values was observed in DEF cells, with relatively low R_0_ values for H5N8-2014 and H5N6-2017, and notably high values for H5N1-2020-C and H5N1-2022-BB (Figure 3).

Other parameters from the mathematical model did not exhibit a trend as pronounced as those observed in the mechanistic traits. However, on CEF cells the infection rate constant (*β*) tended to be higher for the H5N1 viruses than the other viruses while the *β* was high across all viruses in DEF cells except the H5N6-2017 virus (Figure S2A). The eclipse period was similar for all viruses on CEF cells while wider variation was observed across viruses in DEF cells (Figure S2B). No clear trend could be observed for the infectious period (*τ*_I_) on CEF and DEF cells (Figure S2C). The production rate of infectious virus particles (*p*) was generally higher on DEF cells than on CEF cells except for the H5N8-2014 virus. Furthermore, the virus production rate was generally higher for the H5N1 viruses than the other viruses on CEF cells (Figure S2D). Lastly, the lifetime of released infectious virus particles (*τ*_*v*_) was similar on CEF and DEF cells for the H5N1 viruses but tended to be longer for the other viruses on DEF cells compared to CEF cells (Figure S2E).

## Discussion

In this study, we explored the potential of avian *in vitro* models combined with mathematical modelling to characterize and compare the replication dynamics of a reference panel of eight HPAI virus isolated during epizootics in the Netherlands between 2014 and 2024. In addition to obtaining “traditional” virological parameters such as peak virus replication and the time to 50% cell death, eight new parameters: the *t*_*inf*_, *t*_*gen*_, *R*_0_, *β, τ*_*E*_, *τ*_*I*_, *p* and *τ*_V_ were estimated by using a mathematical model to analyse the experimentally measured infectious virus titre and cytopathogenicity over time on primary chicken embryo fibroblasts (CEF) and duck embryo fibroblasts (DEF).

Although, this approach provides a representation of the phenotypic traits of the viruses on cells, the next challenge and focus of future research is to connect these parameters to *in vivo* within host disease dynamics parameters. Various approaches can be used to connect these *in vitro* and *in vivo* processes. One approach could be to use a phenomenological method, exploring the relationship between *in vitro* and *in vivo* parameters without a specific hypothesis regarding the underlying biological processes. An example of this approach is the exploration of direct links between *in vitro* parameters for cell cultures and parameters that describe disease dynamics at much higher biological levels such as that of an animal or human host. This approach was successfully applied in a mathematical model describing the *in vitro* dynamics of avian influenza viruses in human cell lines, where several parameters were found to correlate to some extent to replication fitness, transmission fitness, or virulence in ferrets [35]. For the HPAI viruses in our panel (H5N8-2014, H5N8-2016, H5N6-2017, H5N8-2020 and H5N1-2020-AB), virulence has been previously investigated following the intravenous pathogenicity index (IVPI) procedure in chickens and Pekin ducks resulting in a score from low virulence (0.00) to high virulence (3.00) [38]. Although the IVPI did not identify differences in virulence in chickens for these viruses (all IVPI scores 3) the IVPI scores in Pekin ducks were low for the viruses isolated from chickens (H5N8-2014 1.87, H5N8-2020 1.74 and H5N1-2021-AB 1.96) and high for the viruses isolated from Pekin ducks (H5N8-2016 2.99 and H5N6-2017 3.00). Interestingly, only the HPAI viruses isolated from Pekin ducks (H5N8-2016 and H5N6-2017) induced more rapid cell death in DEF cells than in CEF cells, suggesting a potential cellular correlate for virulence in poultry.

Replication fitness in chickens and Pekin ducks following intra respiratory challenge has been previously investigated for the H5N8-2014, H5N8-2016, H5N6-2017, H5N1-2020-C, and H5N1-2021-AB viruses, as well as a virus of the same genotype as H5N8-2020 (tested only in chickens)[8, 10, 36]. The shedding duration in poultry infected with HPAI is likely influenced by the virulence of the virus, which often results in high and rapid mortality. As a result, peak virus titre may provide a more accurate measure for assessing the replication dynamics of HPAI in poultry. Overall, peak virus shedding in chickens is slightly lower than in Pekin ducks [15]. A similar trend was observed *in vitro*, with lower peak virus titers in CEF cells than in DEF cells. Interestingly, this difference in median peak virus replication was wider for the H5N8-2016 and H5N6-2017 viruses isolated from ducks (2 and 1.56 log_10_ TCID_50_/mL higher in DEF than in CEF cells compared to a difference of < 1 log_10_ TCID_50_/mL for the other six HPAI viruses). This suggests that the host species from which a virus is isolated may influence its relative viral fitness. A similar host-dependent difference in virus replication was observed in embryonated chicken and Pekin duck eggs [37]. Similarly, the infecting time was generally longer on CEF cells than on DEF cells indicating a trend for faster replication dynamics on duck cells than on chicken cells.

When exploring the replication behavior of the different virus strains within chickens, the H5N6-2017 and H5N8-2020 viruses showed the lowest reported peak virus shedding, followed by the H5N8-2014 and H5N8-2016 viruses, while the highest peak shedding was reported for the H5N1-2020-C and H5N1-2021-AB viruses [8, 10, 36]. This pattern was generally reproduced in CEF cells, with the longest infecting times (indicative of slower replication) measured for H5N8-2020 and H5N6-2017 viruses and the shortest infecting times measured for the H5N1 strains. Although peak virus replication in CEF cells did not fully replicate the in vivo shedding trends, a general trend for apparent higher peak replication was observed for the H5N1 virus strains compared to the H5N6 and H5N8 strains. In ducks, challenge experiments in Pekin ducks, showed similar peak virus shedding titers across different H5Nx HPAI virus strains, with slightly lower titers measured for the H5N8-2014 virus [8, 10, 36]. Similarly, in DEF cells, the lowest peak replication was measured for the H5N8-2014 virus. Unlike peak virus replication, the estimated infecting time in DEF cells – also the longest for the H5N8-2014 virus - appeared to be similar across all HPAI viruses as it was observed for the peak virus shedding observed *in vivo*. In summary, replication trends observed in DEF and CEF cells suggest that the infecting time may be a good indicator of viral replication dynamics *in vivo*.

Inferring transmissibility fitness from *in vitro* experiments is challenging, as obtaining transmission data at the host level (*in vivo*) is difficult through both epidemiological and experimental studies. Relevant parameters describing the transmission dynamics in a host population (not in cells) would be the transmission rate (*β*\*), which is the number of new infections generated by one infectious individual per unit of time, the infectious period (*t*_*1*_*), which is the number of days an infected individual is contagious, and the basic reproduction number (R_0_*), which at host level is the average number of new infections an infectious individual generates in a susceptible population. Among the HPAI viruses in our panel, transmission experiments quantifying these parameters in chickens and Pekin ducks were calculated for viruses from the same genotype as the H5N8-2016 and the H5N1-2021-AC virus. For the H5N8 2016, *β*\*, *t*_*1*_ ∗ and R_0_* for chickens were 0.77 day^-1^, 2.3 days and 1.8 respectively (Gonzales, unpublished data) and for Pekin ducks 3.4 day^-1^ 4.9 days and 16.5 [38]. For the H5N1-2021-AC virus, *β*\*, *t*_*1*_ ∗ and R_0_* and for chickens were 1.13 day^-1^, 3.2 days and 3.64 respectively [39] and for Mule ducks 10.9 day^-1^, 8.1 days and 88 [40]. A common finding from these studies, along with several others evaluating transmission in these host species [41], is that *β*\*, *t*_*1*_* and R_0_* are higher in ducks than in chickens. A similar trend was observed in our *in vitro* study, with the infecting time being generally faster (potentially indicating a higher *β*\* the generation time being generally longer (potential indicating a longer *t*_*1*_*) and R_0_ being generally higher in duck cells than chicken cells. However, these are preliminary observations and transmission data for the other viruses in our panel would be required to better evaluate whether these *in vitro* parameters are useful cellular indicators for the relative transmissibility fitness of these viruses in chickens and ducks. It should be noted, that *in vitro* models cannot be used to infer absolute values for transmission dynamics in live birds.

This study has several limitations, key among those is the small number of assessed viruses and the much limited corresponding *in vivo* data, particularly regarding transmission, potentially compromising accurate *in vitro* to *in vivo* translation. Transmission experiments for all the genotypes included in the reference panel will be required to better determine which *in vitro* replication parameters most accurately reflect *in vivo* replication and transmission. Such data would increase the reliability of using this panel to benchmark novel viruses for risk assessment. Furthermore, in the context of low pathogenic avian influenza (LPAI) primary embryo fibroblasts may be a limited model for assessing transmissibility compared to tissue specific models (such as respiratory or digestive tract cells). However, HPAI viruses are fundamentally pantropic. Therefore, embryo fibroblasts may reflect the dynamics of a systemic infection, which is a critical determinant of replication fitness, virulence and transmission fitness in poultry. Alternatively, primary embryo kidney cells, intestinal organoids or airway organoids can be used to expand the model to LPAI as these cells endogenously express trypsin-like proteases [42-44]. For Pekin ducks, *in vitro* virulence may be better assessed in endothelial cell models, as endothelial tropism is a key determinant of HPAIV pathogenesis [45]. One other limitation is that the *in vivo* data from our reference panel only includes poultry isolates. As replication kinetics appear to be influenced by the source host (e.g., Pekin duck or chicken) from which the virus was isolated, the reference panel must be expanded to include wild bird isolates. Such an expansion is essential to enhance *in vitro* to *in vivo* translation for the risk assessment of HPAI viruses circulating in wild bird populations.

From a mathematical modelling perspective, another limitation is the lack of model validation. Although the model provided a good fit to the viral load and cell data, this does not guarantee that model-derived quantities such as the R_0_, infecting time and generation time, are accurately estimated. Other mathematical models may reproduce the data equally well, while yielding different estimates of these summary measures [46]. It is therefore important to validate the model by comparing its predictions for these summary measures with independent experimentally measured values. This is experimentally challenging, since it would require introducing a small number of infectious cells in a susceptible cell population and tracking the secondary infections from these cells over time. An alternative approach would be to evaluate the model’s predictive power by simulating independent cell culture experiments with more complex viral dynamics, for example in the presence of a second competing virus strain, antibodies or antiviral compounds, and comparing the predicted and observed infection dynamics [47]. Future studies should focus on validation strategies to better establish the predictive value of the model.

In this study, we demonstrate that the combination of *in vitro* experimentation and mathematical modeling can provide a more complete, quantitative characterization (described by 10 parameters) of the virus replication dynamics in avian cells representative of our target hosts: chickens and ducks. We also suggest how these *in vitro* parameters could possibly translate to *in vivo* conditions. In this way, we characterized a reference panel of closely related H5Nx viruses to benchmark unknown (future) viruses, therefore assisting in the rapid assessment of HPAI infection risks across different poultry species. By generating several parameters, our approach identified differences in the virus replication dynamics of HPAI viruses on chicken cells that could not be identified *in vivo*. This *in vitro* model approach could be extended to encompass wild bird or mammalian cells. Therefore, this study builds towards rapid characterization and risk assessment of HPAI viruses protecting both animal health and public health.

## Acknowledgements

This work was funded by the Dutch Ministry of Agriculture, Nature and Food Quality, projects WOT-01-003-096 and KB-37-003-047-WBVR with additional funding by Discovery Grants RGPIN/3774-2022 (C.A.A.B.) from the Natural Sciences and Engineering Research Council of Canada (NSERC, www.nserc-crsng.gc.ca), and by the RIKEN Center for Interdisciplinary Theoretical and Mathematical Sciences (iTHEMS, ithems.riken.jp) (C.A.A.B.).

## Notes

### Competing Interest Statement

The authors have declared no competing interest.

### Summary of Updates

Peer reviewed manuscript and ORCID ID from author added

