## Supplementary Information for "Mathematical modelling of in vitro replication dynamics for multiple highly pathogenic avian influenza clade 2.3.4.4 viruses in chicken and duck cells"

**Contents**

1. Definitions, dimensions and initial values or prior distributions for state variables and parameters in the mathematical model
  - Tables S1 and S2
2. Genetic analysis and potential host shift adaptations
  - Tables S3 and S4
3. Replication dynamics without mathematical modelling
  - Figure S1
4. Summary of the posterior distributions of model parameters for each combination of virus strain and host cell
  - Figure S2.
5. Fine-tuning of the ABC-SMC algorithm
  - Figures S3 and S4
6. Assessment of the convergence of posterior distributions and parameter identifiability for each combination of virus-strain and host-cell
    6.1. H5N1-AB on CEF cells | Figures S5 to S7
    6.2. H5N1-AB on DEF cells | Figures S8 to S10
    6.3. H5N1-AC on CEF cells | Figures S11 to S13
    6.4. H5N1-AC on DEF cells | Figures S14 to S16
    6.5. H5N1-BB on CEF cells | Figures S17 to S19
    6.6. H5N1-BB on DEF cells | Figures S20 to S22
    6.7. H5N1-C on CEF cells | Figures S23 to S25
    6.8. H5N1-C on DEF cells | Figures S26 to S28
    6.9. H5N6-2017 on CEF cells | Figures S29 to S31
    6.10. H5N6-2017 on DEF cells | Figures S32 to S34
    6.11. H5N8-2014 on CEF cells | Figures S35 to S37
    6.12. H5N8-2014 on DEF cells | Figures S38 to S40
    6.13. H5N8-2016 on CEF cells | Figures S41 to S43
    6.14. H5N8-2016 on DEF cells | Figures S44 to S46
    6.15. H5N8-2020 on CEF cells | Figures S47to S49
    6.16. H5N8-2020 on DEF cells | Figures S50 to S52

**1. Definitions, dimensions and initial values or prior distributions for state variables and parameters in the mathematical model**

Table S1. The definitions, dimensions and initial values of state variables in the model.

| State variable | Definition | Dimension | Initial value |
|---|---|---|---|
| $S$ | Concentration of susceptible cells | cells/ml | 175,000 |
| $E$ | Concentration of cells in eclipse phase | cells/ml | 0 |
| $D$ | Concentration of dead cells | cells/ml | 0 |
| $I$ | Concentration of infectious cells | cells/ml | 0 |
| $V$ | Concentration of infectious virus particles | TCID50/ml | 175 |

Table S2. The definitions, dimensions and posterior distributions of model parameters.

| Parameter | Definition | Dimension | Prior distribution[1, 2] |
|---|---|---|---|
| $\beta$ | Transmission parameter | ml/(TCID50*h) | $U_c(-9,1)$ on $\log_{10}$ scale |
| $\tau_E$ | Eclipse period cells | h | $U_c(0,24)$ |
| $\tau_I$ | Infectious period cells | h | $U_c(0.01,24)$ |
| $n$ | Shape parameter of the gamma distribution for the lengths of the eclipse and infectious periods | - | $U_d(1,30)$ |
| $p$ | Virus replication rate by infectious cells | TCID50/(cell*h) | $U_c(-2,3)$ on $\log_{10}$ scale |
| $\alpha$ | Infectious virus particles lost from the supernatant due to the infection of a cell. | TCID50/cell | $U_c(-2,2)$ on $\log_{10}$ scale |
| $\tau_V$ | Lifetime of an infectious virus particle | h | $U_c(-1,2)$ on $\log_{10}$ scale |

[1]) Symbols $U_c$ and $U_d$ represent the continuous and discrete uniform distributions, respectively.

[2]) The prior distribution of some parameters was specified on a $\log_{10}$ instead of linear scale in order to increase the probability of sampling parameter values over a range spanning several orders of magnitude.

### 2. Genetic analysis and potential host shift adaptations

Table S3. Virulence factors that differ between the eight HPAI viruses. Coloured cells indicates the presence of the marker in the first column. Increased replication in mammalian cells (red), increased virulence in chickens (yellow), decreased virulence in mice (green), decreased replication in mammalian cells (blue) and disruption of the second sialic acid binding site (light green) is indicated by the colours.

| Marker | Effect | H5N8-2014 | H5N8-2016 | H5N6-2017 | H5N8-2020 | H5N1-2020-C | H5N1-2021-AB | H5N1-2021-AC | H5N1-2022-BB |
|---|---|---|---|---|---|---|---|---|---|
| NS-1:P42S | Increased replication in mammalian cells | | | | | | | | |
| NS-1:C138F,NS-1:K55E,NS-1:K66E | Increased replication in mammalian cells | | | | | | | | |
| NP:M105V | increased virulence in chickens | | | | | | | | |
| PB2:I292V | Increased replication in mammalian cells | | | | | | | | |
| PB2:K389R | Increased replication in mammalian cells | | | | | | | | |
| PB2:L89V,PB2:G309D | Increased replication in mammalian cells | | | | | | | | |
| PB2:L89V,PB2:G309D,PB2:T339K,PB2:R477G,PB2:I495V,PB2:K627E,PB2:A676T | Increased replication in mammalian cells | | | | | | | | |
| PB1-F2:N66S | Increased replication in mammalian cells | | | | | | | | |
| PA:K158R | Increased replication in mammalian cells | | | | | | | | |
| PA:Q400P | Decreased virulence in mice | | | | | | | | |
| NP:Y52N | Decreased replication in mammalian cells | | | | | | | | |
| NA-1:A369I | Distruption of the second sialic acid binding site (2SBS) | | | | | | | | |
| NA-1:K432E | Distruption of the second sialic acid binding site (2SBS) | | | | | | | | |

Table S4. Phylogenetic grouping of the selected HPAI viruses based on each genome segment. Different colors indicate sequences with p-distance ≤ 0.015 and is identified for each genome segment separately.

| Abbreviation | PB2 | PB1 | PA | HA | NP | NA | MP | NS |
|---|---|---|---|---|---|---|---|---|
| H5N8-2014 | | | | | | | | |
| H5N8-2016 | | | | | | | | |
| H5N6-2017 | | | | | | | | |
| H5N8-2020 | | | | | | | | |
| H5N1-2020-C | | | | | | | | |
| H5N1-2021-AB | | | | | | | | |
| H5N1-2021-AC | | | | | | | | |
| H5N1-2022-BB | | | | | | | | |

#### 3. Replication dynamics without mathematical modelling

Figure S1. Chicken embryo fibroblasts stained with DAPI. Complete removal of the monolayer due to cytopathogenicity is observed at 34hpi except for minor positive staining for H5N1-2022-BB.

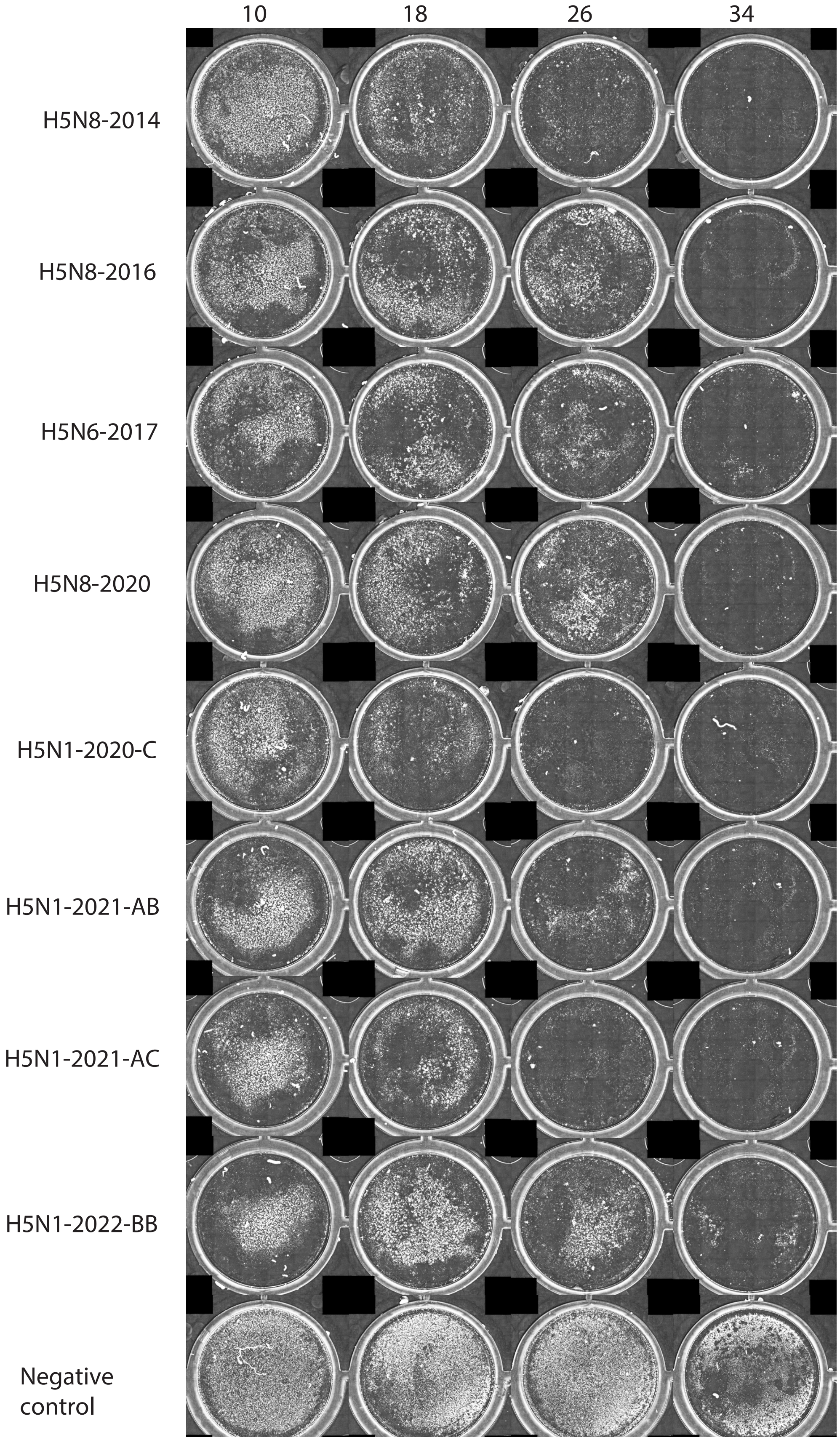

**4. Summary of the posterior distributions of model parameters for each combination of virus strain and host cell**

Figure S2. Additional model output parameters. The medians are displayed by the blue and red dots.

A

Infection rate constant ($Log_{10}$) β

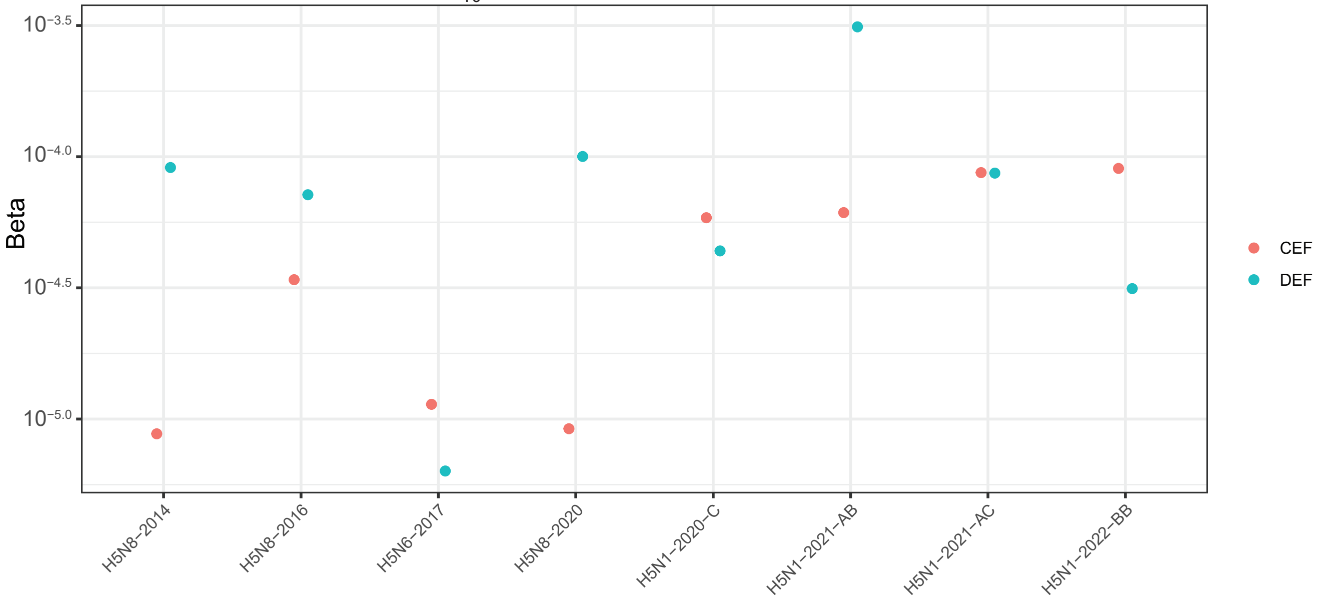


B

Eclipse period $t_E$

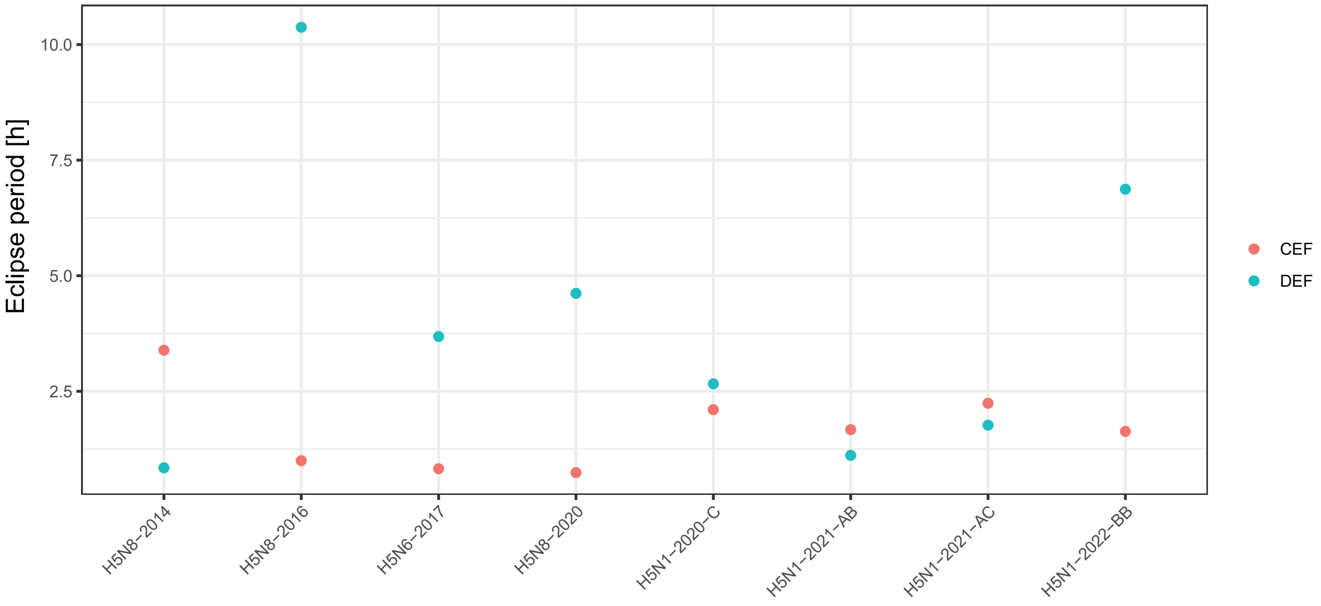

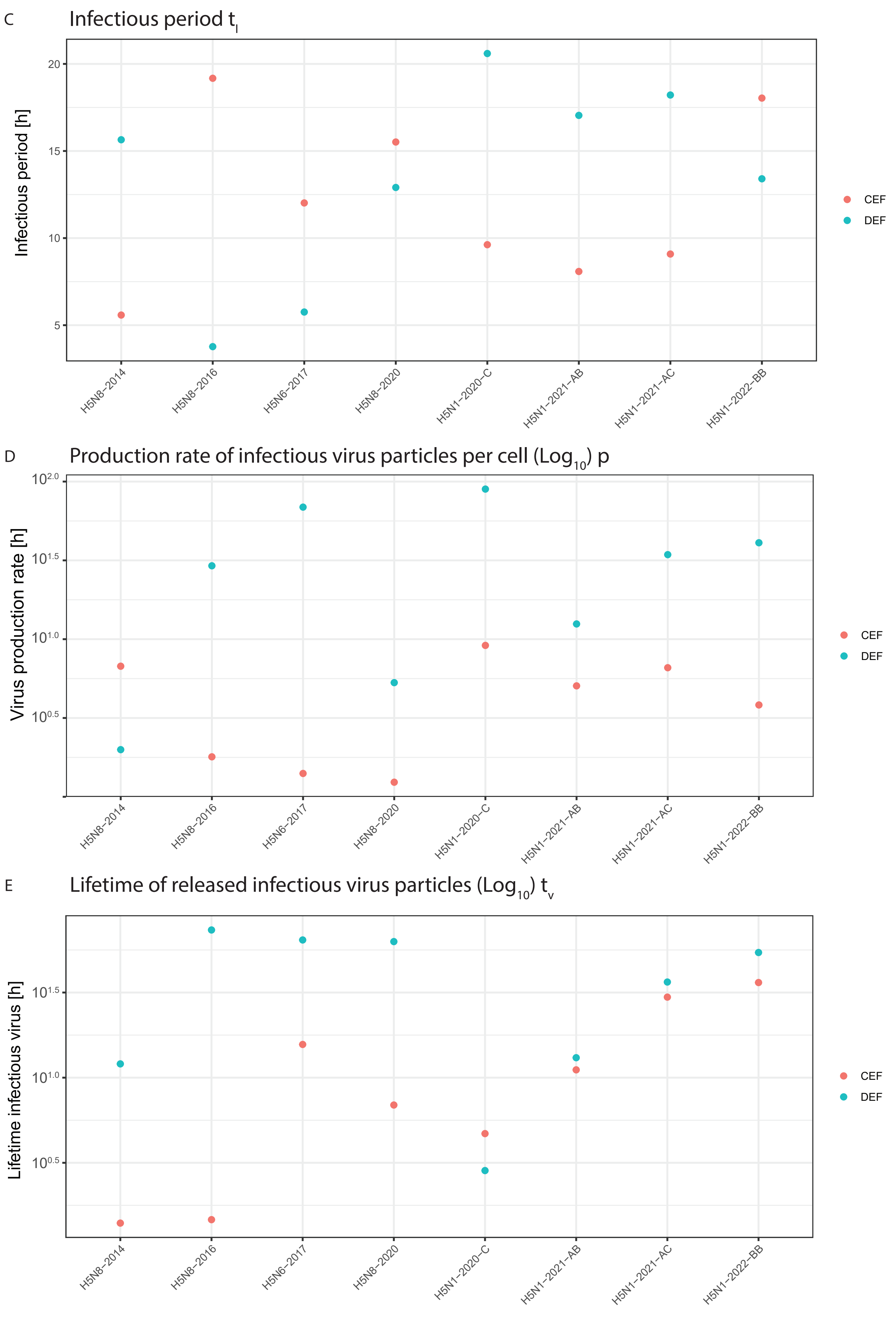

C
Infectious period $t_I$
Infectious period [h]
20
15
10
5
H5N8-2014
H5N8-2016
H5N6-2017
H5N8-2020
H5N1-2020-C
H5N1-2021-AB
H5N1-2021-AC
H5N1-2022-BB
CEF
DEF
D
Production rate of infectious virus particles per cell ($Log_{10}$) p
Virus production rate [h]
$10^{2.0}$
$10^{1.5}$
$10^{1.0}$
$10^{0.5}$
H5N8-2014
H5N8-2016
H5N6-2017
H5N8-2020
H5N1-2020-C
H5N1-2021-AB
H5N1-2021-AC
H5N1-2022-BB
CEF
DEF
E
Lifetime of released infectious virus particles ($Log_{10}$) $t_v$
Lifetime infectious virus [h]
$10^{1.5}$
$10^{1.0}$
$10^{0.5}$
H5N8-2014
H5N8-2016
H5N6-2017
H5N8-2020
H5N1-2020-C
H5N1-2021-AB
H5N1-2021-AC
H5N1-2022-BB
CEF
DEF

**5. Fine-tuning of the ABC-SMC algorithm**

Figure S3. The median and 95% credible interval (CI) of the posterior distribution of model parameters fitted to data for strain H5N1-AB on CEF cells for increasing values of the particle population size.

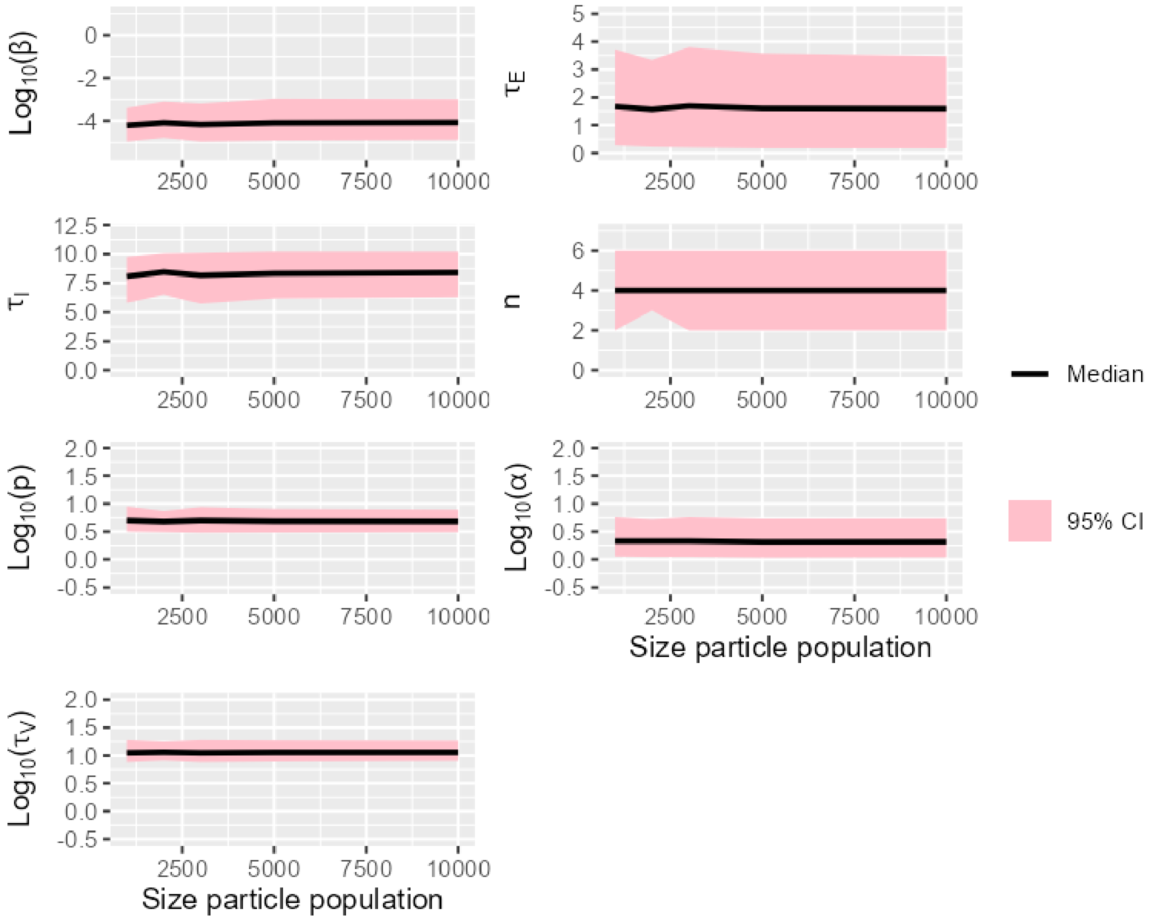

Figure S4. The median and 95% credible interval (CI) of the posterior distributions of model parameters fitted to data for strain H5N1-AB on CEF cells for increasing values of parameter $\theta$, which influences the width of the perturbation kernel for continous parameters.

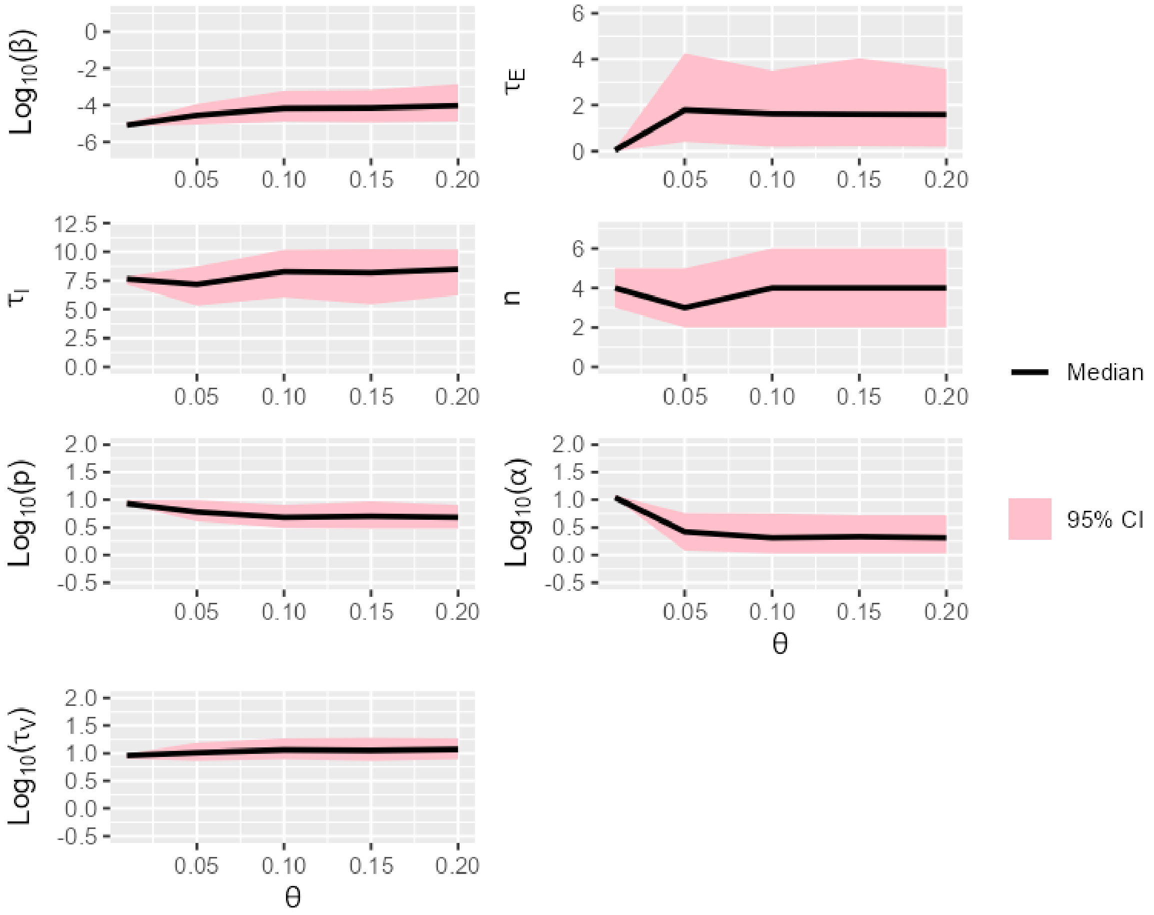

**6. Assessment of the convergence of posterior distributions and parameter identifiability for each combination of virus-strain and host-cell**

6.1 H5N1-AB on CEF cells

Figure S5. The shifts in the distributions of model parameters during the ABC-SMC analysis of data from strain H5N1-AB on CEF cells, when moving from the first particle population (containing accepted particles sampled from the prior distribution) to the posterior distribution via intermediate particle populations.

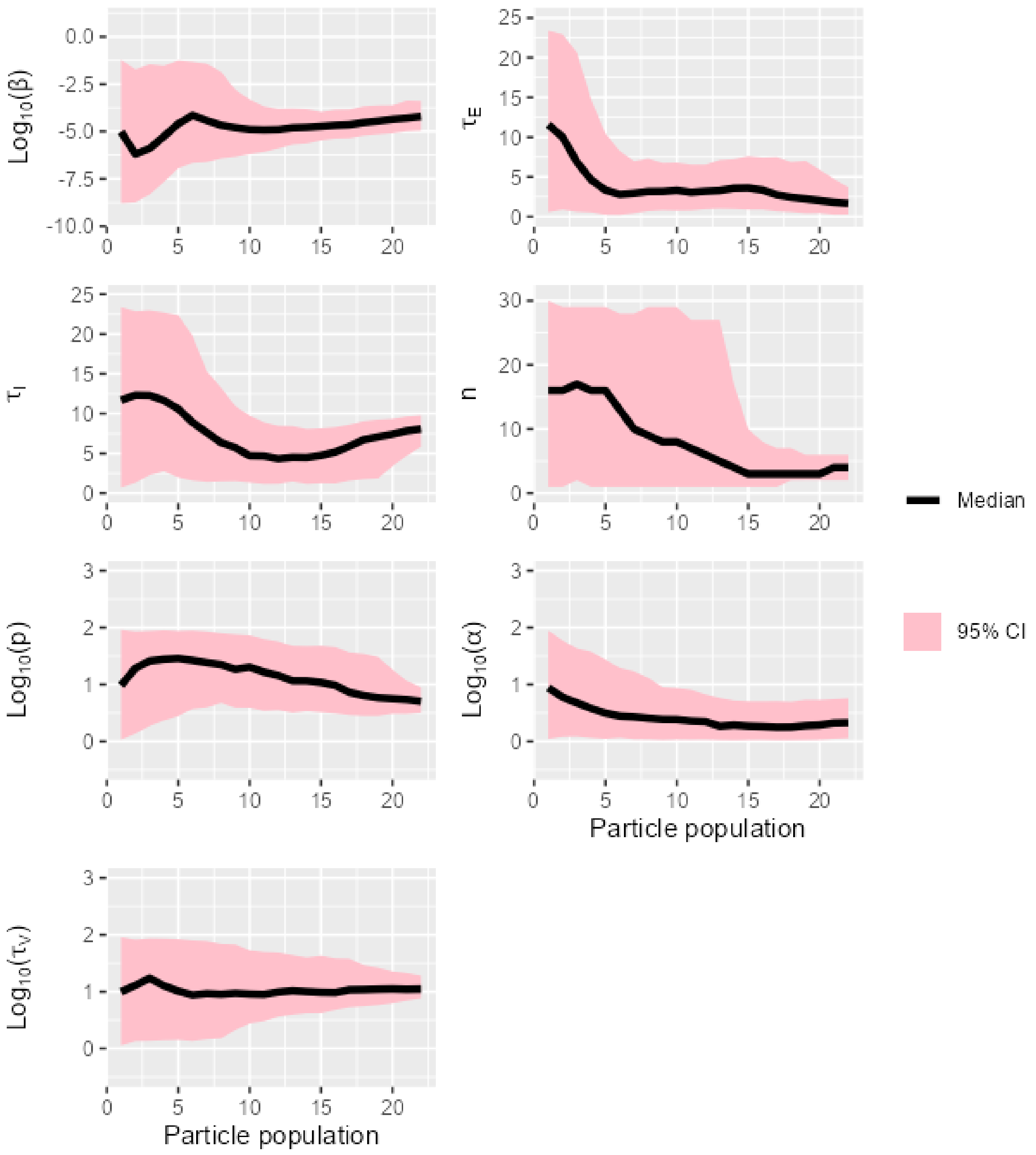

Figure S6. The overlap between prior and posterior distributions of model parameters for strain H5N1-AB on CEF cells.

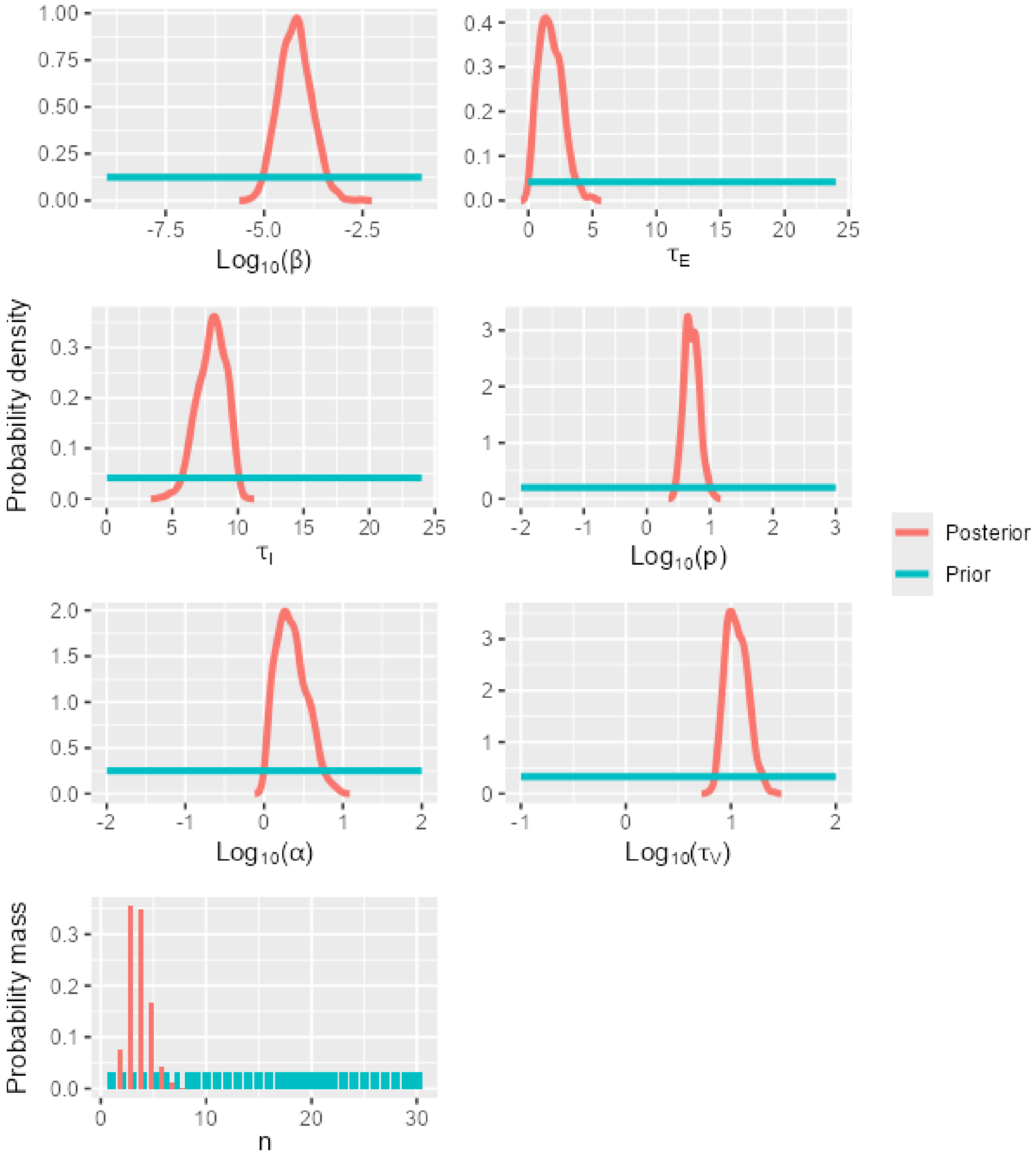

Figure S7. A matrix showing Kendall's rank correlation coefficient and the significance of the correlation (0.05 level, indicated by *) for pairs of model parameters calibrated using data for strain H5N1-AB on CEF cells.

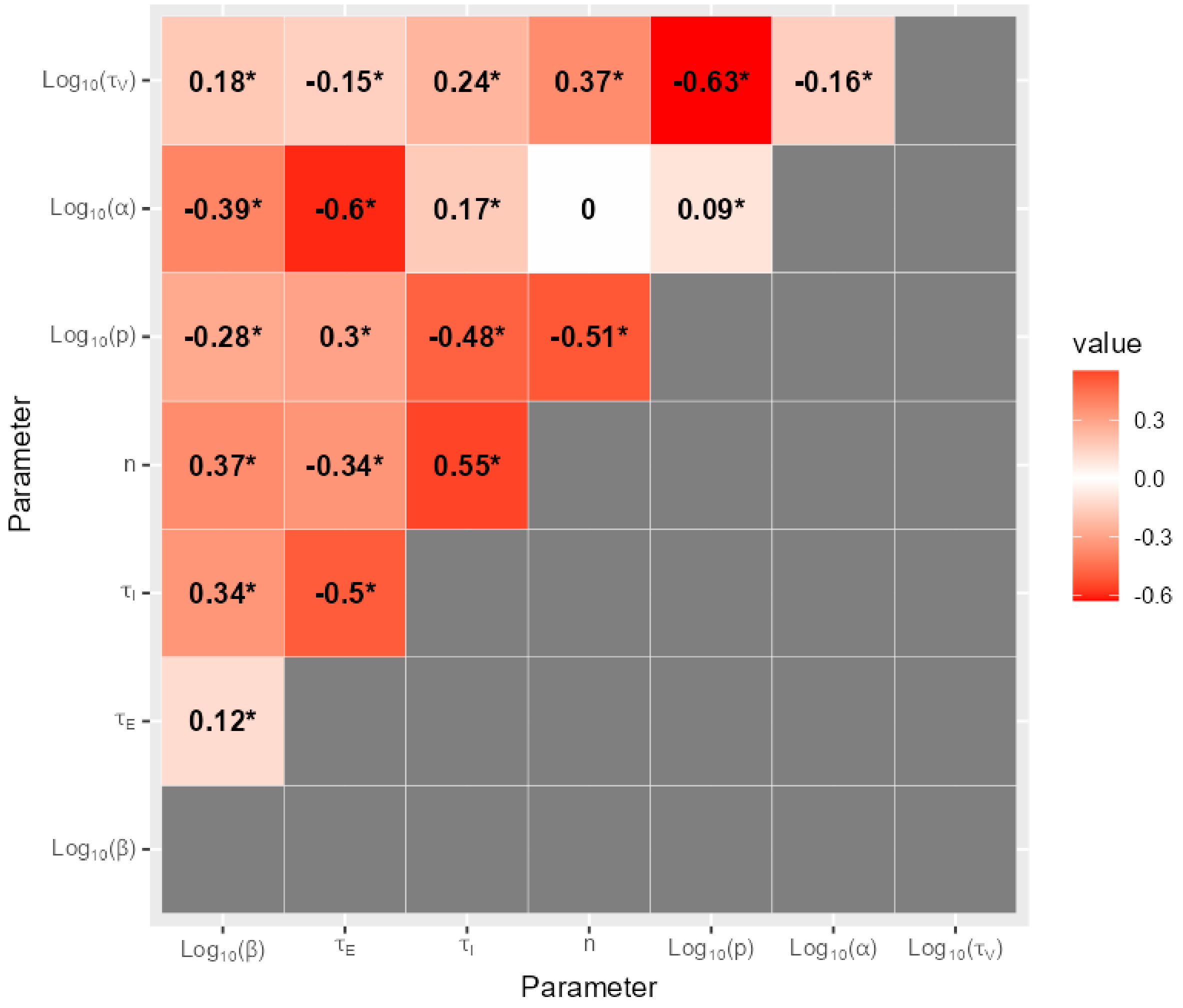

6.2 H5N1-AB on DEF cells

Figure S8. The shifts in the distributions of model parameters during the ABC-SMC analysis of data from strain H5N1-AB on DEF cells, when moving from the first particle population (containing accepted particles sampled from the prior distribution) to the posterior distribution via intermediate particle populations.

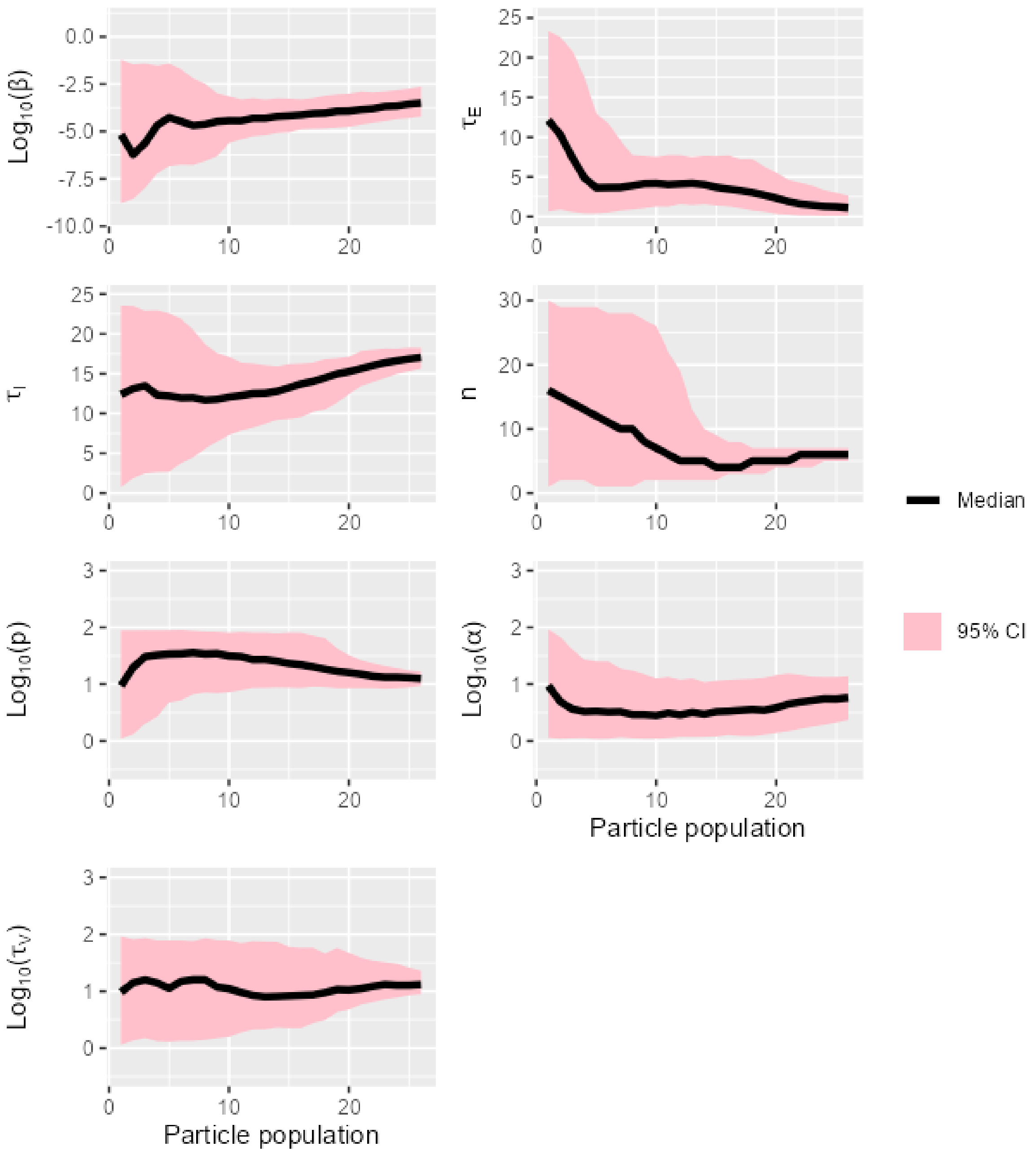

Figure S9. The overlap between prior and posterior distributions of model parameters for strain H5N1-AB on DEF cells.

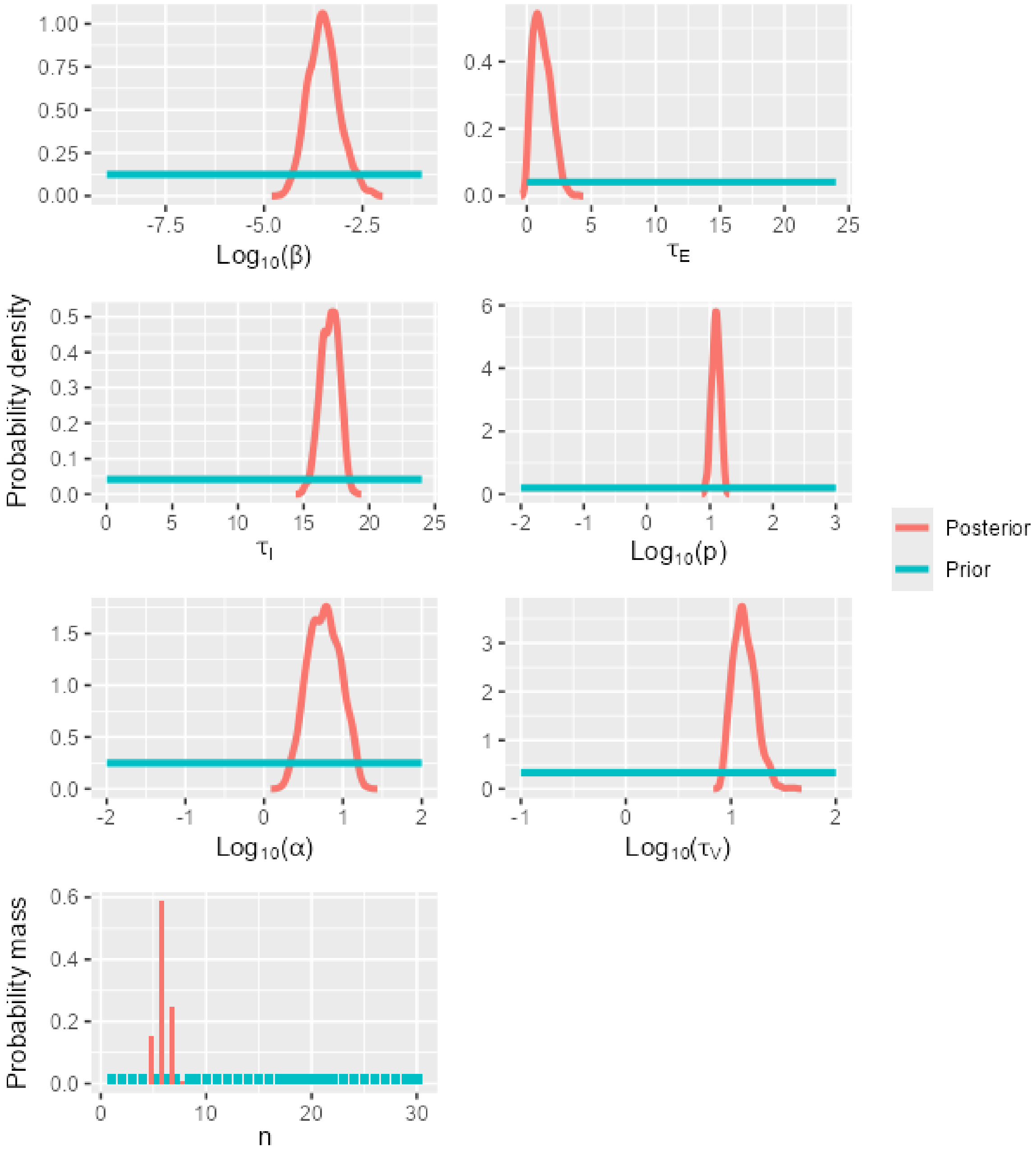

Figure S10. A matrix showing Kendall's rank correlation coefficient and the significance of the correlation (0.05 level, indicated by *) for pairs of model parameters calibrated using data for strain H5N1-AB on DEF cells.

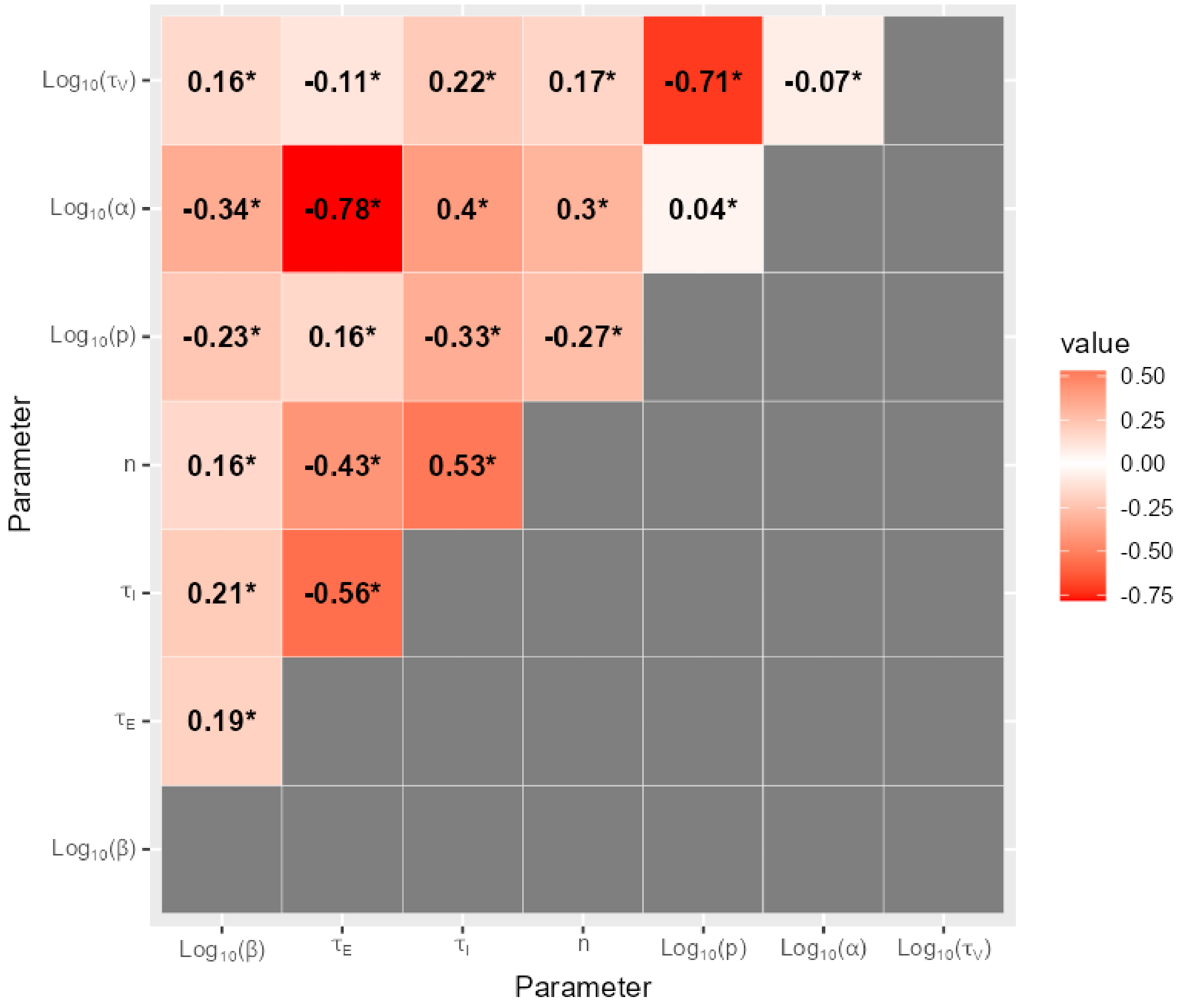

6.3 H5N1-AC on CEF cells

Figure S11. The shifts in the distributions of model parameters during the ABC-SMC analysis of data from strain H5N1-AC on CEF cells, when moving from the first particle population (containing accepted particles sampled from the prior distribution) to the posterior distribution via intermediate particle populations.

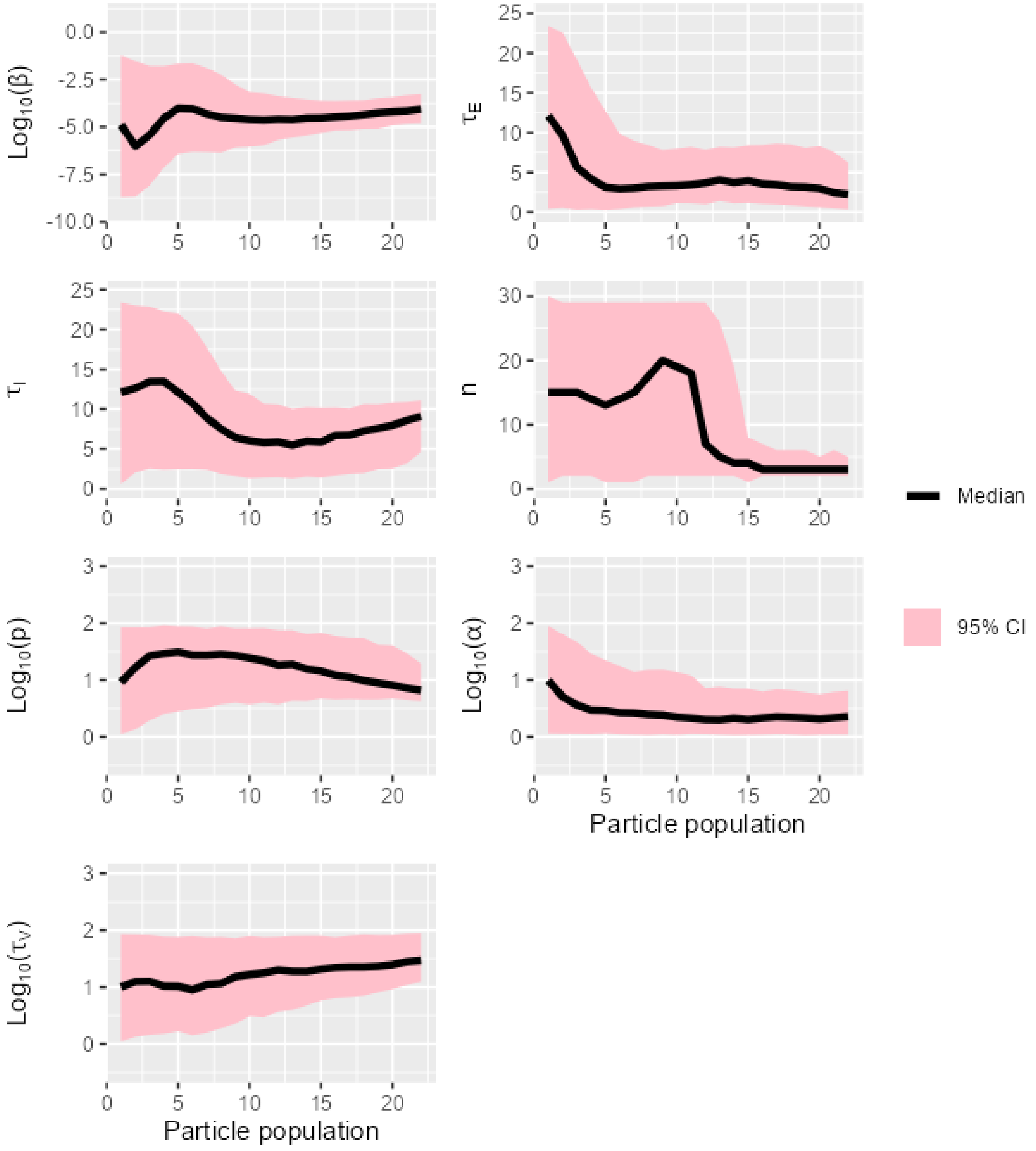

Figure S12. The overlap between prior and posterior distributions of model parameters for strain H5N1-AC on CEF cells.

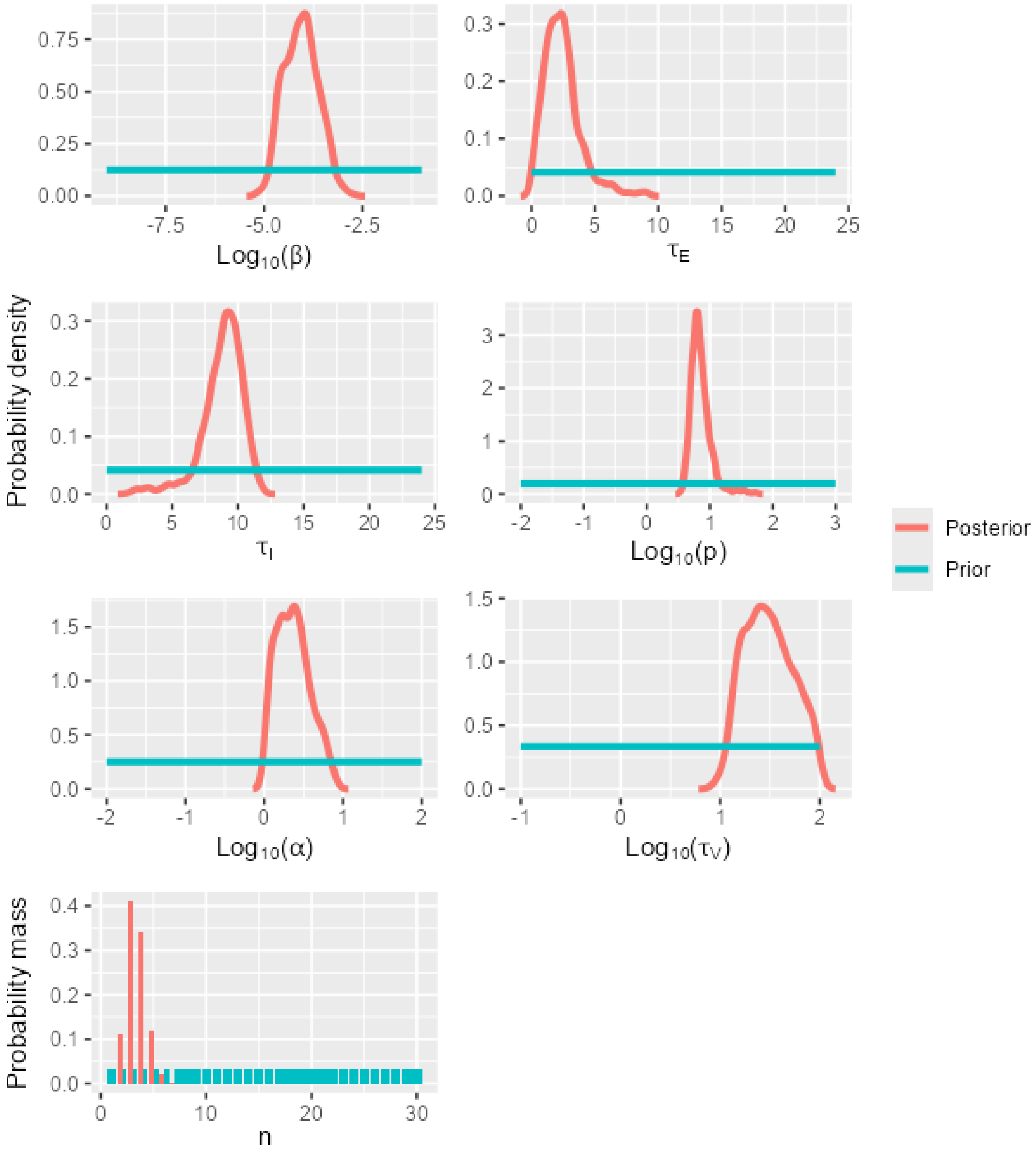

Figure S13. A matrix showing Kendall's rank correlation coefficient and the significance of the correlation (0.05 level, indicated by *) for pairs of model parameters calibrated using data for strain H5N1-AC on CEF cells.

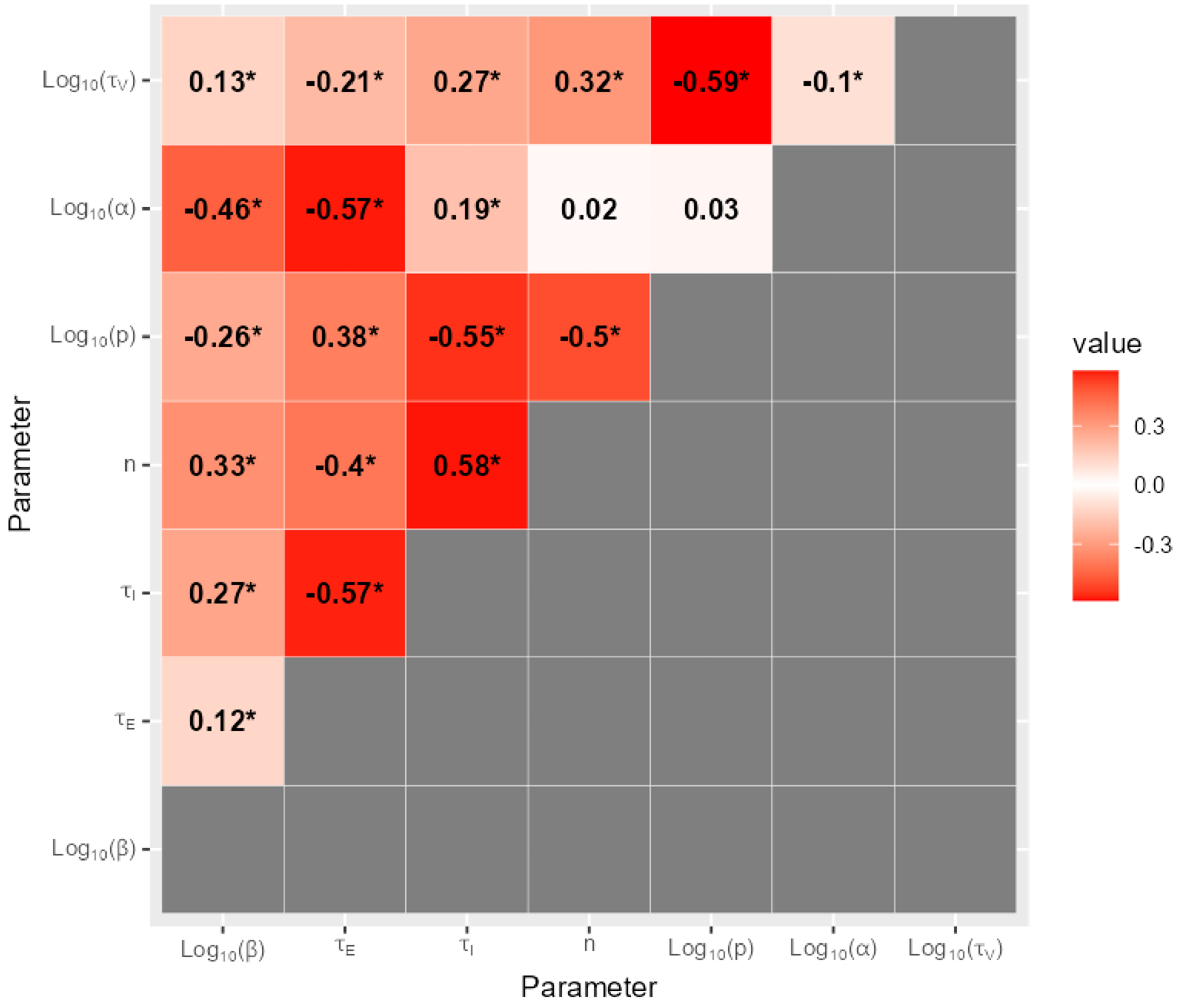

6.4 H5N1-AC on DEF cells

Figure S14. The shifts in the distributions of model parameters during the ABC-SMC analysis of data from strain H5N1-AC on DEF cells, when moving from the first particle population (containing accepted particles sampled from the prior distribution) to the posterior distribution via intermediate particle populations.

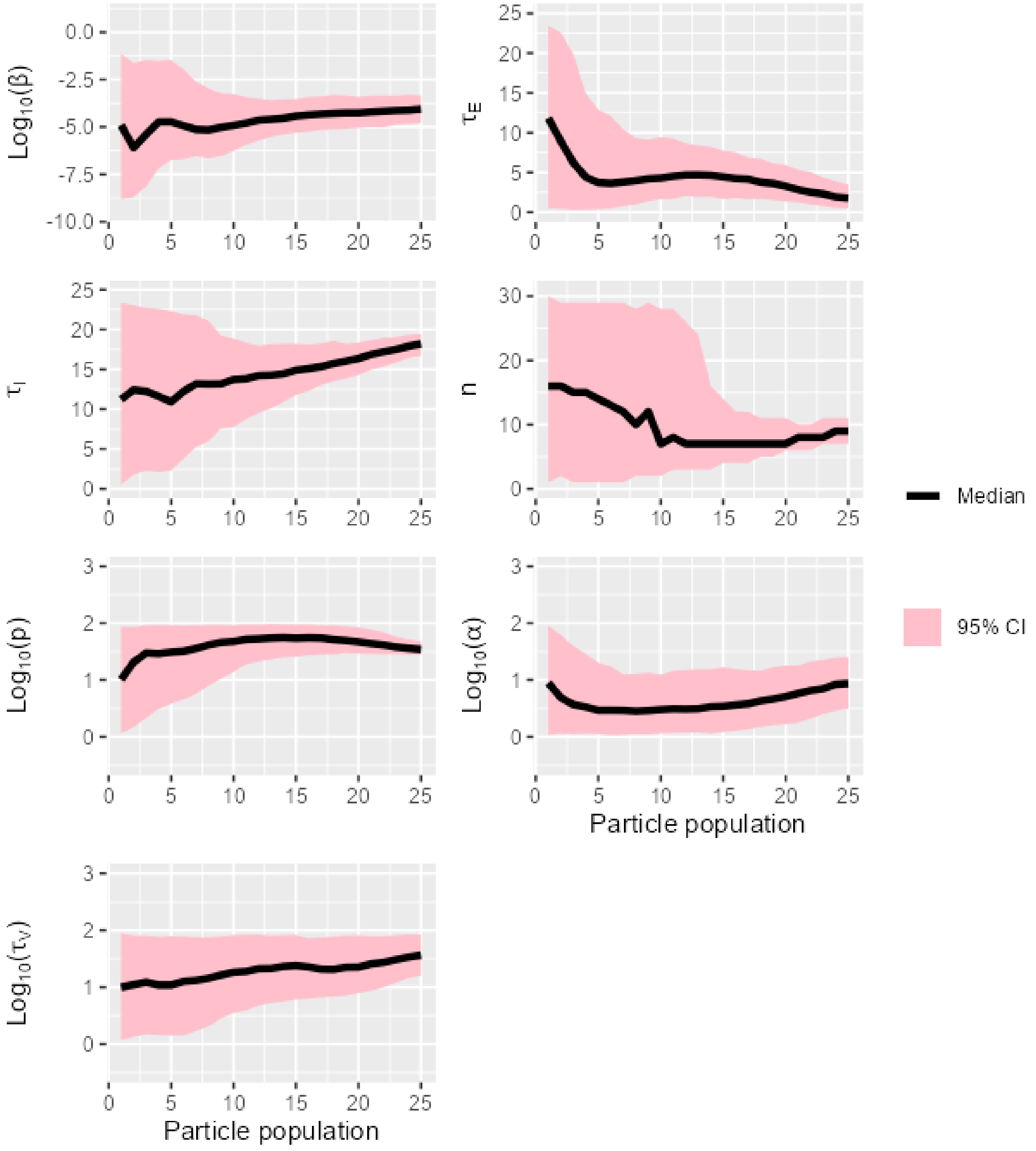

Figure S15. The overlap between prior and posterior distributions of model parameters for strain H5N1-AC on DEF cells.

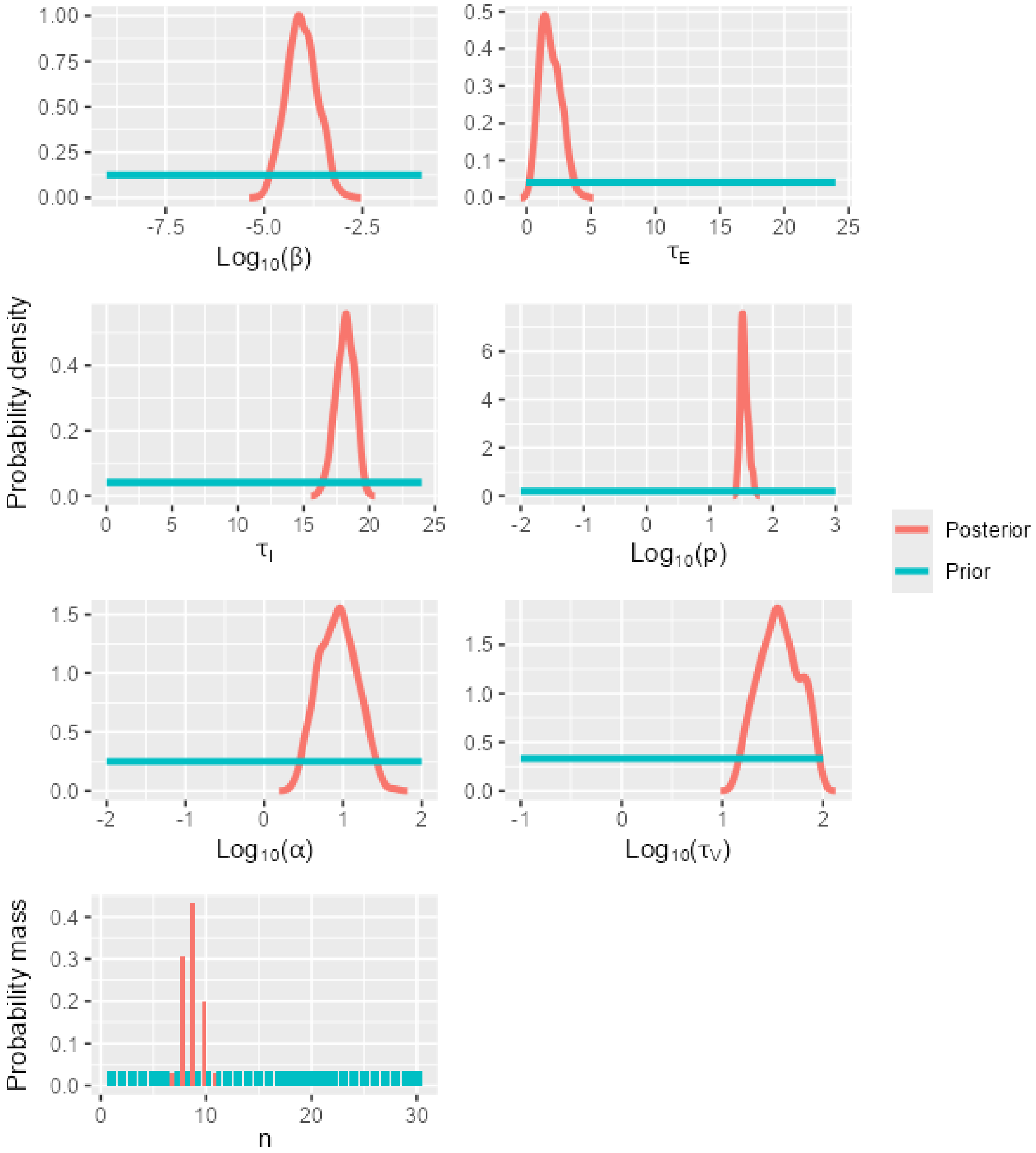

Figure S16. A matrix showing Kendall's rank correlation coefficient and the significance of the correlation (0.05 level, indicated by *) for pairs of model parameters calibrated using data for strain H5N1-AC on DEF cells.

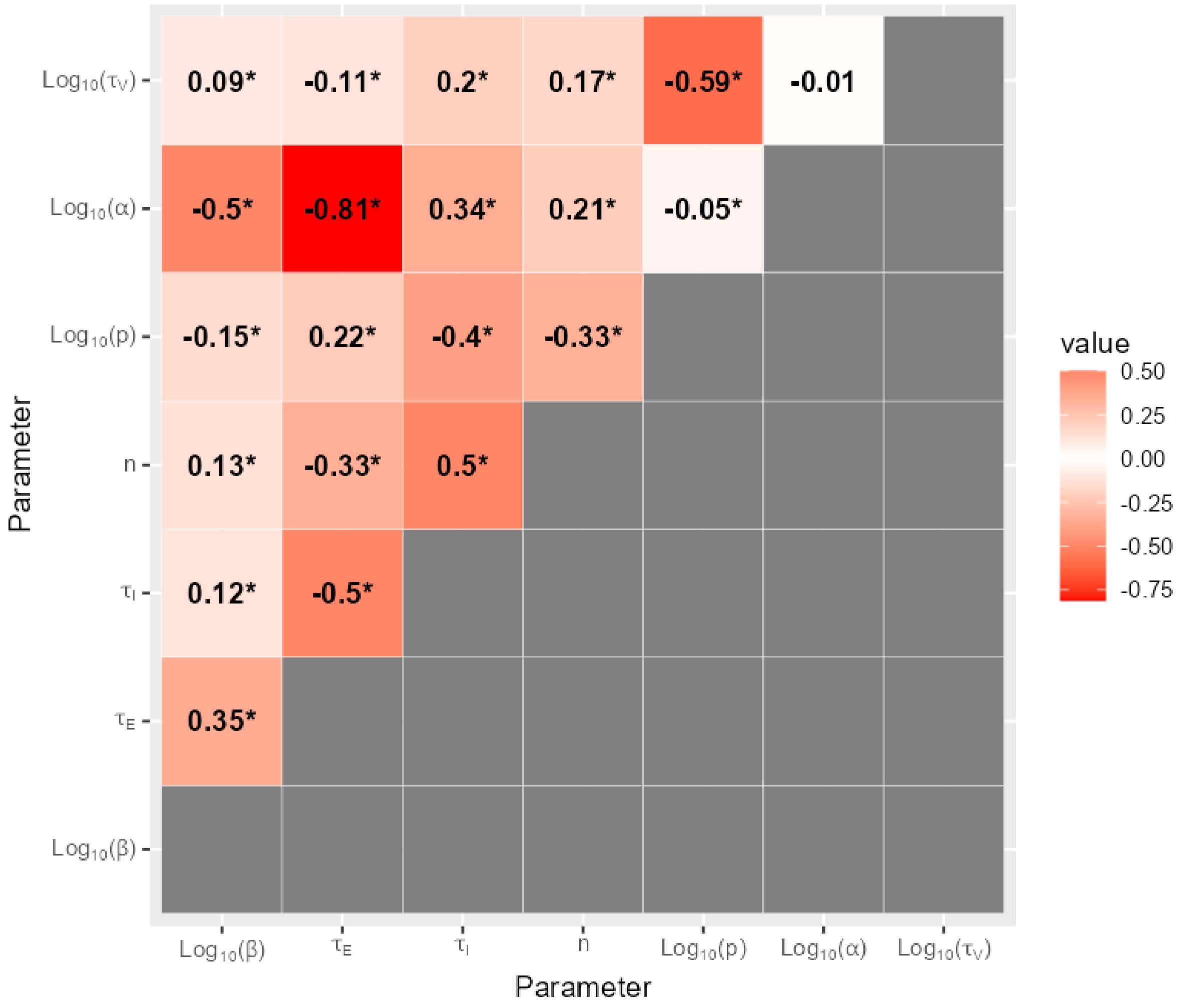

6.5 H5N1-BB on CEF cells

Figure S17. The shifts in the distributions of model parameters during the ABC-SMC analysis of data from strain H5N1-BB on CEF cells, when moving from the first particle population (containing accepted particles sampled from the prior distribution) to the posterior distribution via intermediate particle populations.

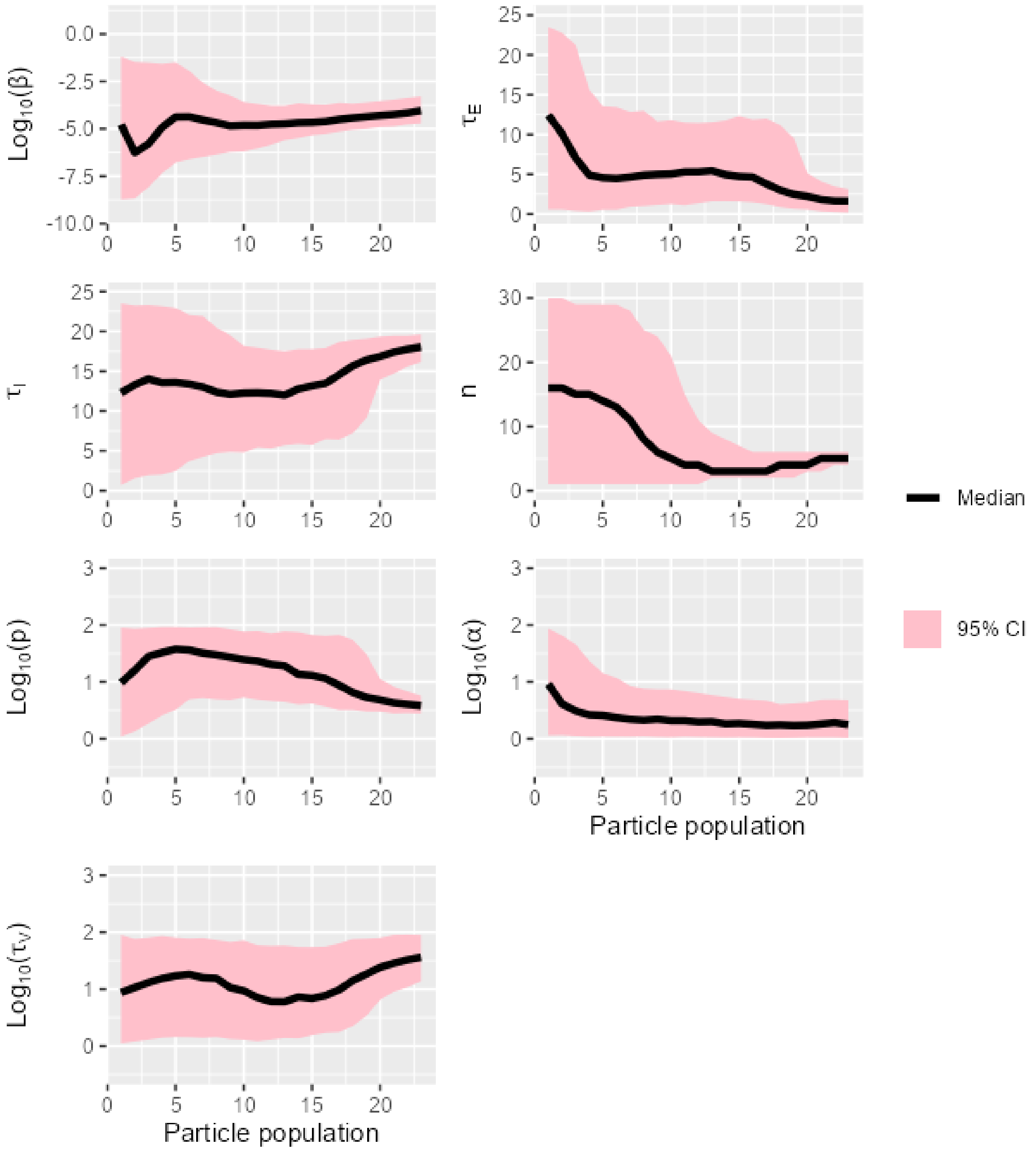

Figure S18. The overlap between prior and posterior distributions of model parameters for strain H5N1-BB on CEF cells.

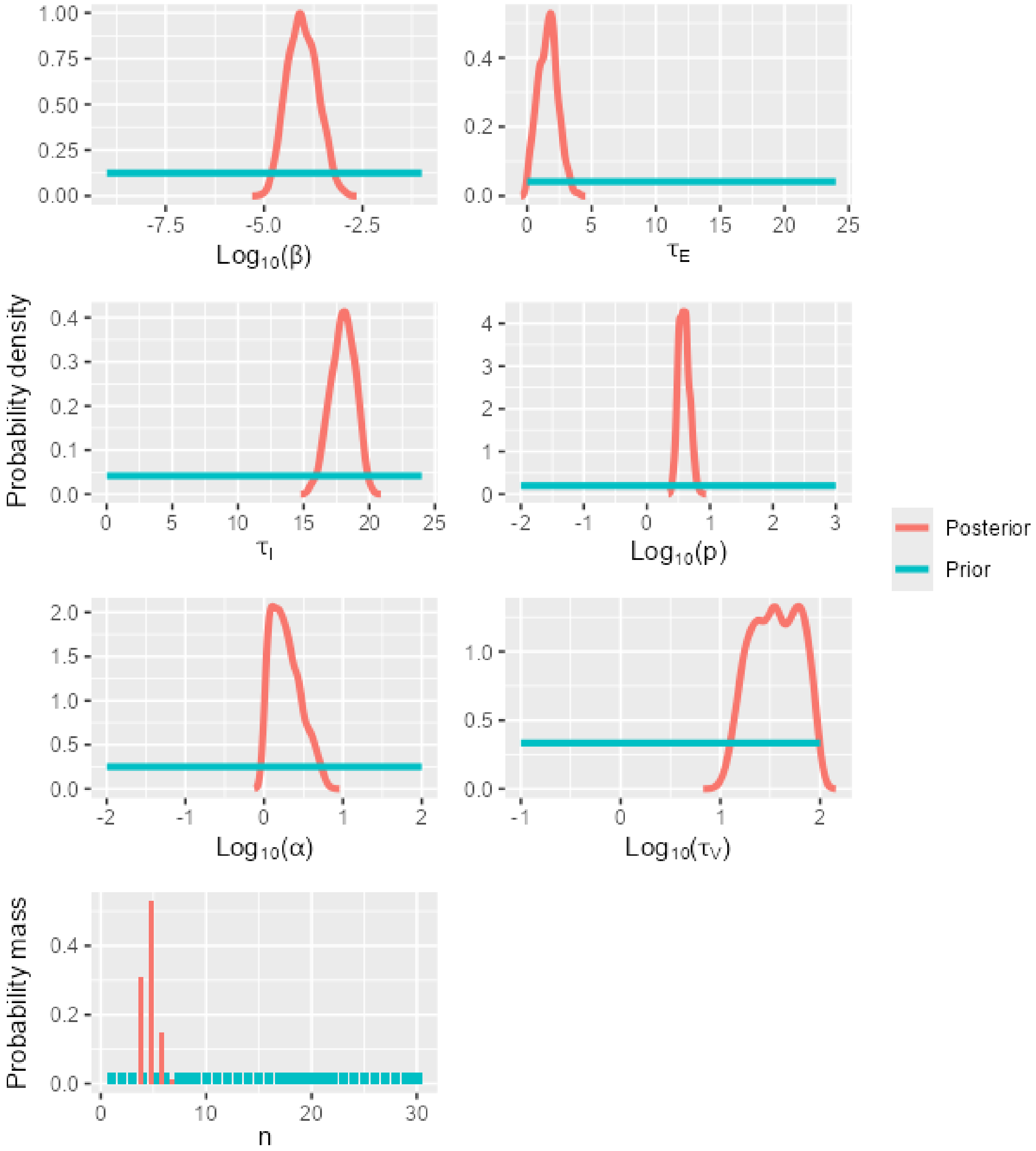

Figure S19. A matrix showing Kendall's rank correlation coefficient and the significance of the correlation (0.05 level, indicated by *) for pairs of model parameters calibrated using data for strain H5N1-BB on CEF cells.

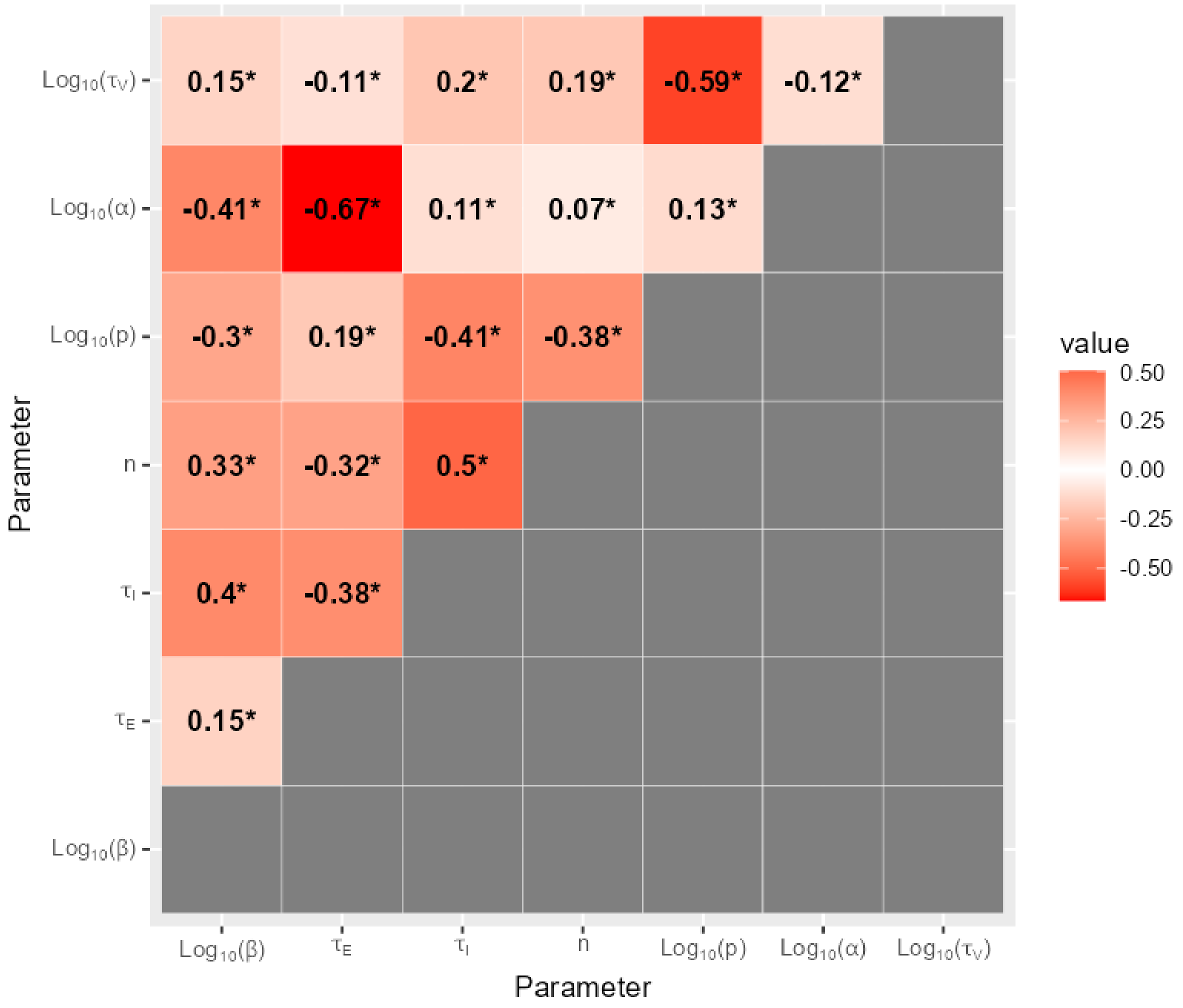

6.6 H5N1-BB on DEF cells

Figure S20. The shifts in the distributions of model parameters during the ABC-SMC analysis of data from strain H5N1-BB on DEF cells, when moving from the first particle population (containing accepted particles sampled from the prior distribution) to the posterior distribution via intermediate particle populations.

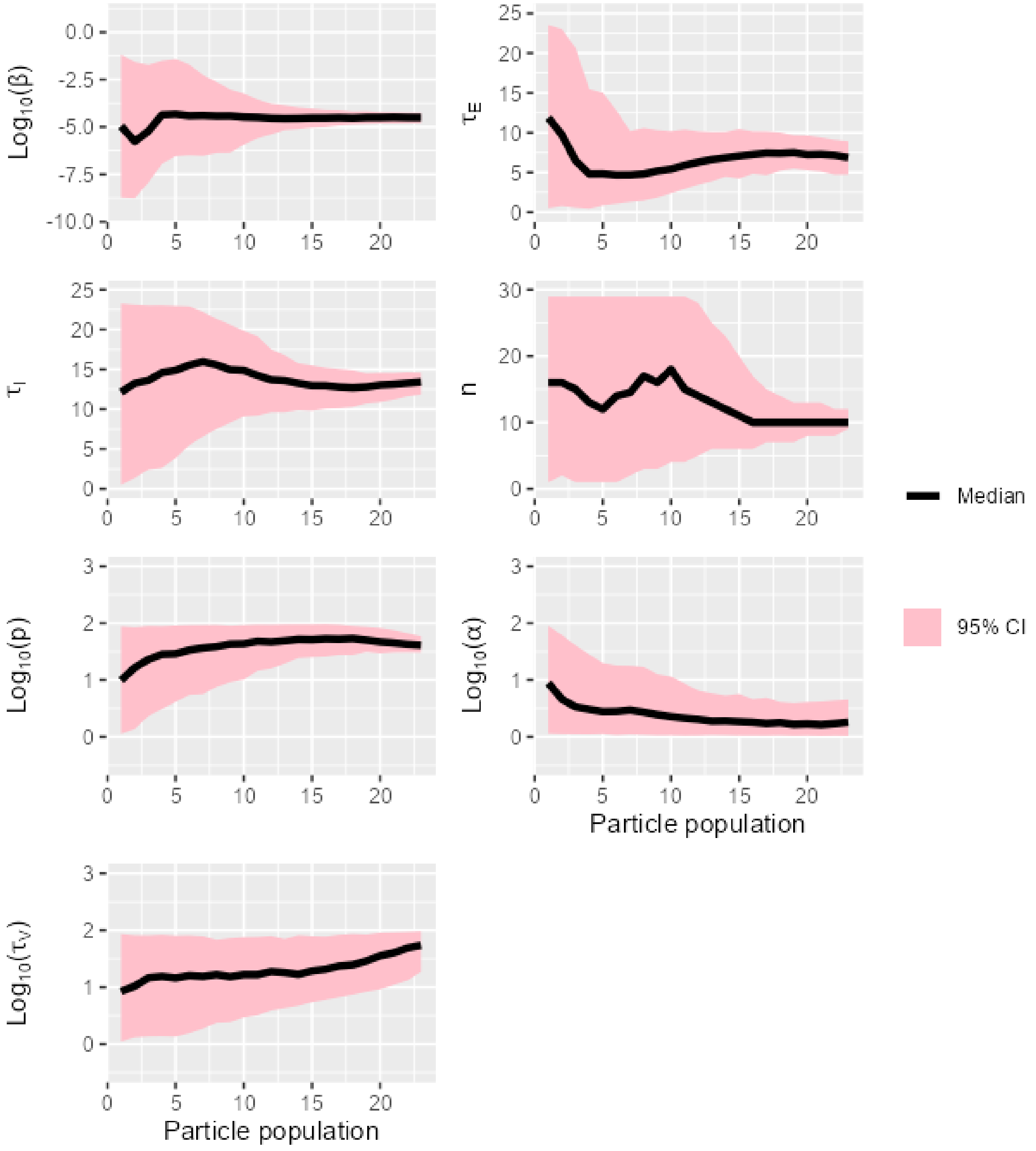

Figure S21. The overlap between prior and posterior distributions of model parameters for strain H5N1-BB on DEF cells.

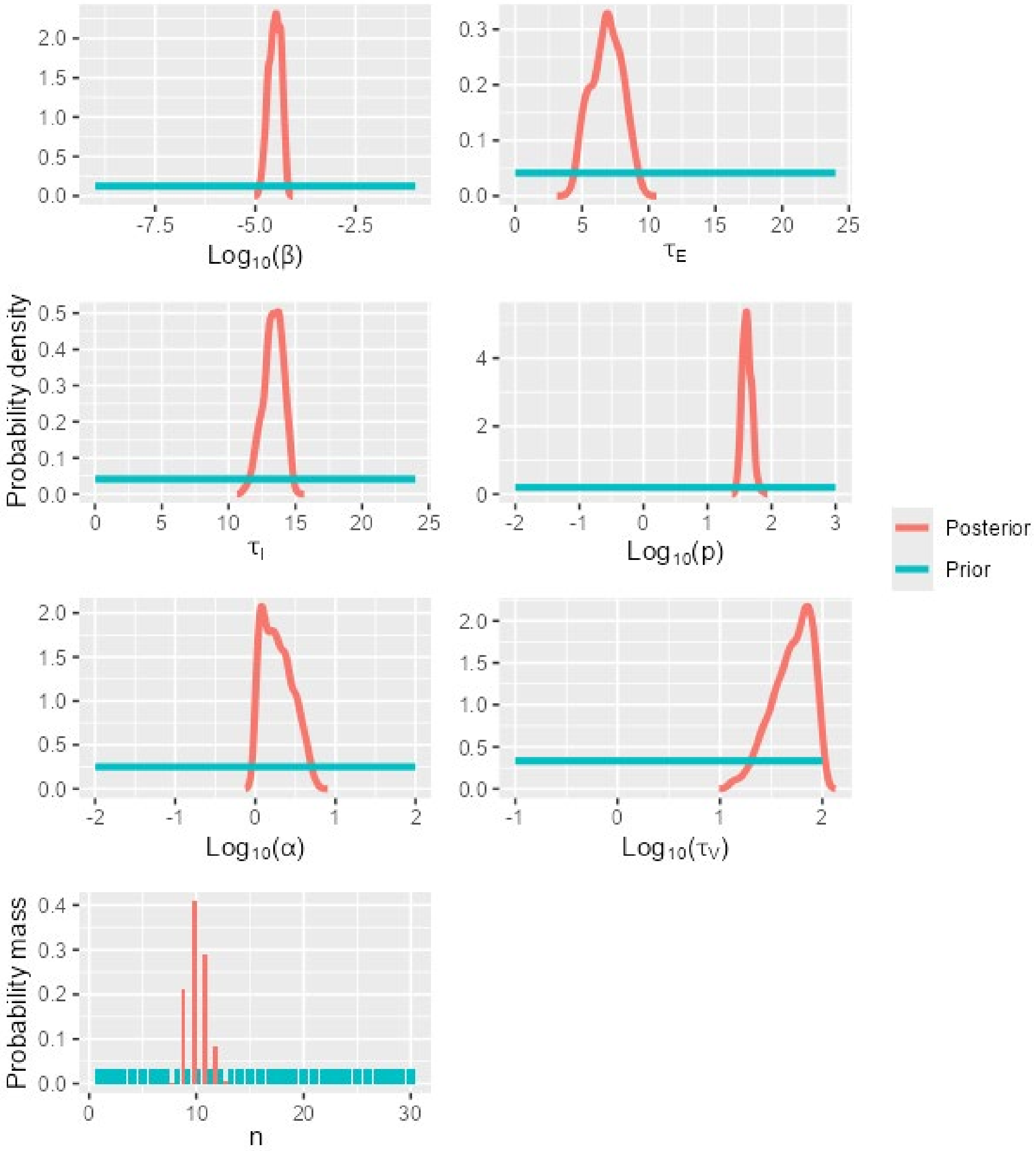

Figure S22. A matrix showing Kendall's rank correlation coefficient and the significance of the correlation (0.05 level, indicated by *) for pairs of model parameters calibrated using data for strain H5N1-BB on DEF cells.

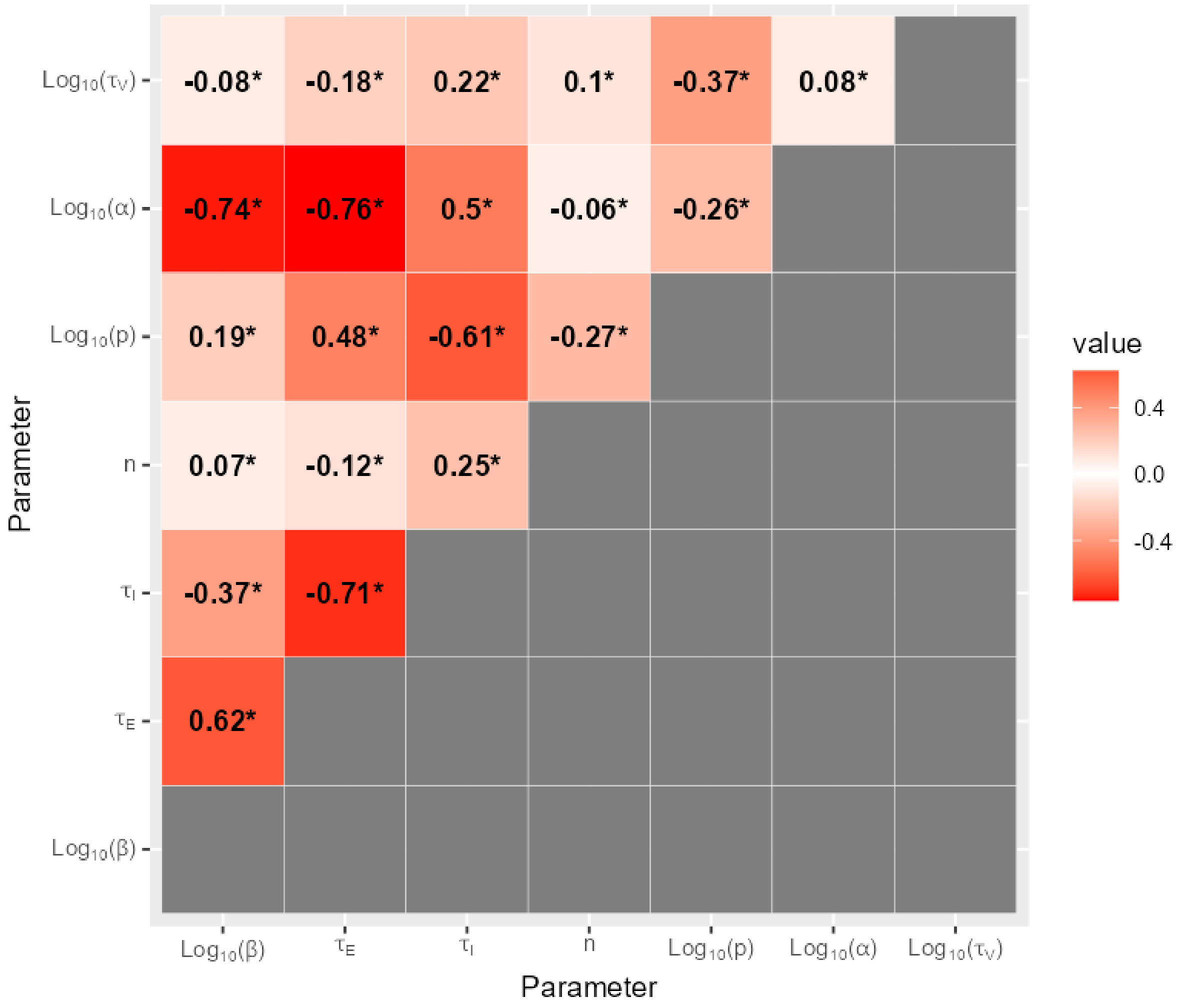

6.7 H5N1-C on CEF cells

Figure S23. The shifts in the distributions of model parameters during the ABC-SMC analysis of data from strain H5N1-C on CEF cells, when moving from the first particle population (containing accepted particles sampled from the prior distribution) to the posterior distribution via intermediate particle populations.

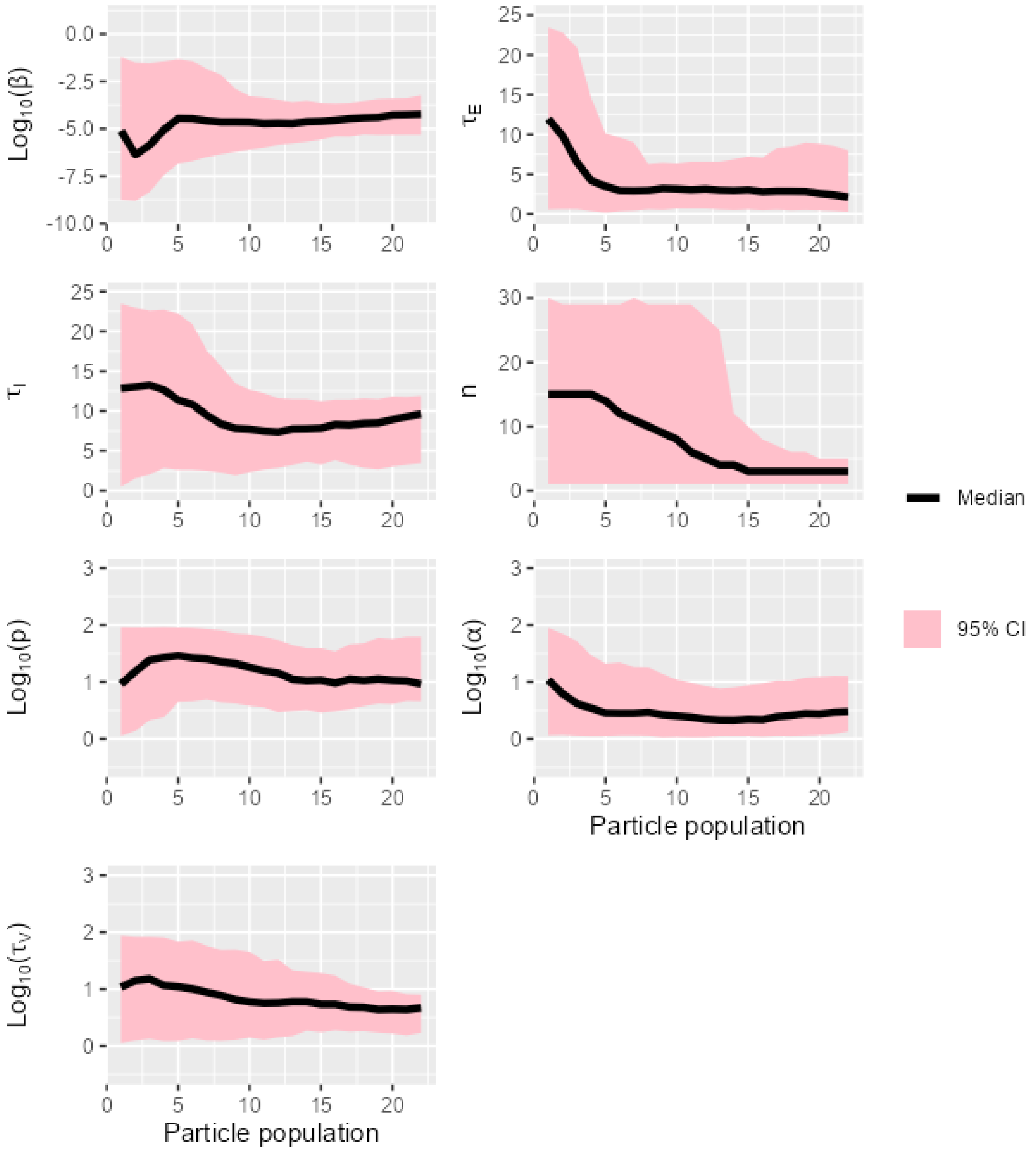

Figure S24. The overlap between prior and posterior distributions of model parameters for strain H5N1-C on CEF cells.

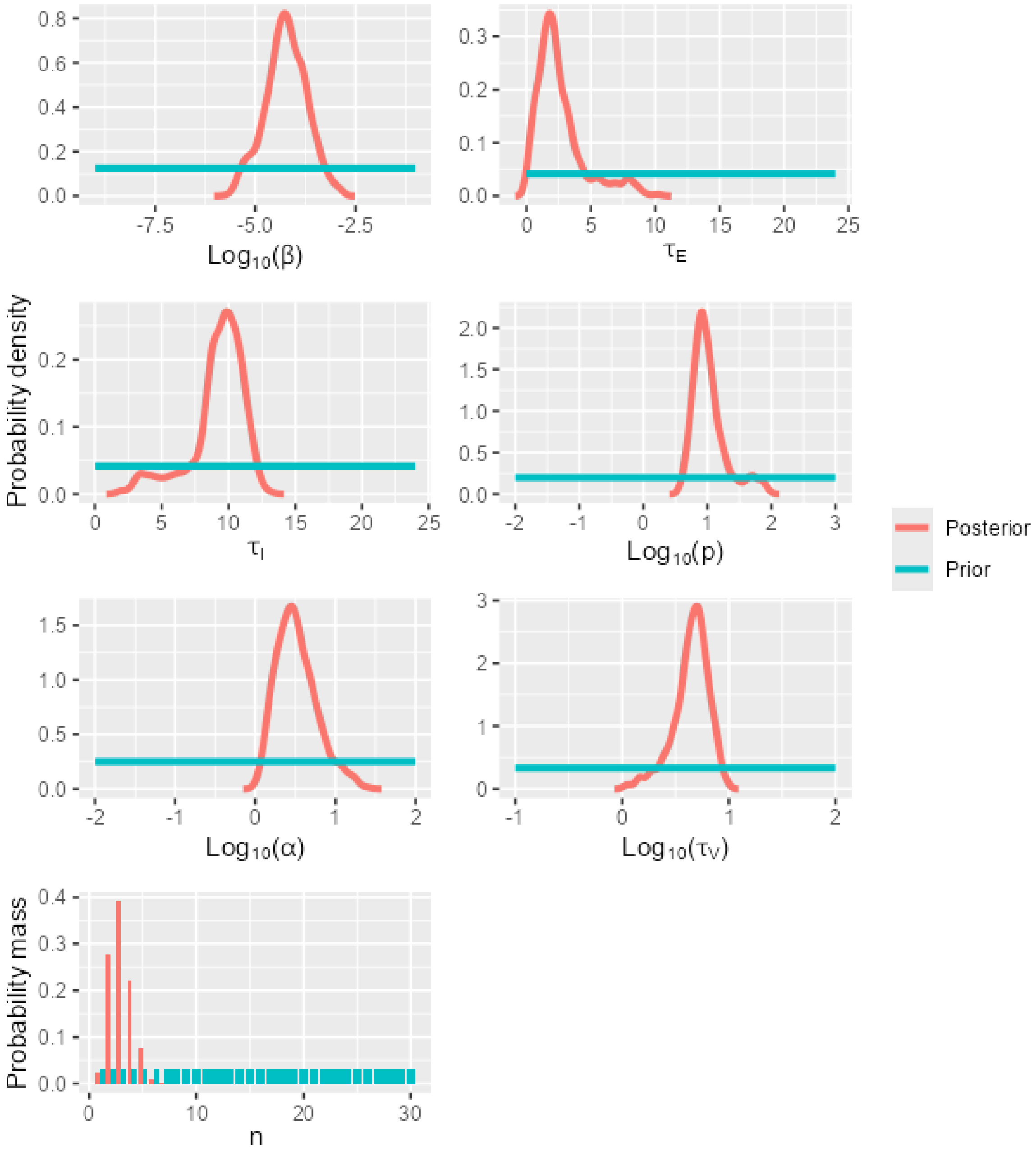

Figure S25. A matrix showing Kendall's rank correlation coefficient and the significance of the correlation (0.05 level, indicated by *) for pairs of model parameters calibrated using data for strain H5N1-C on CEF cells.

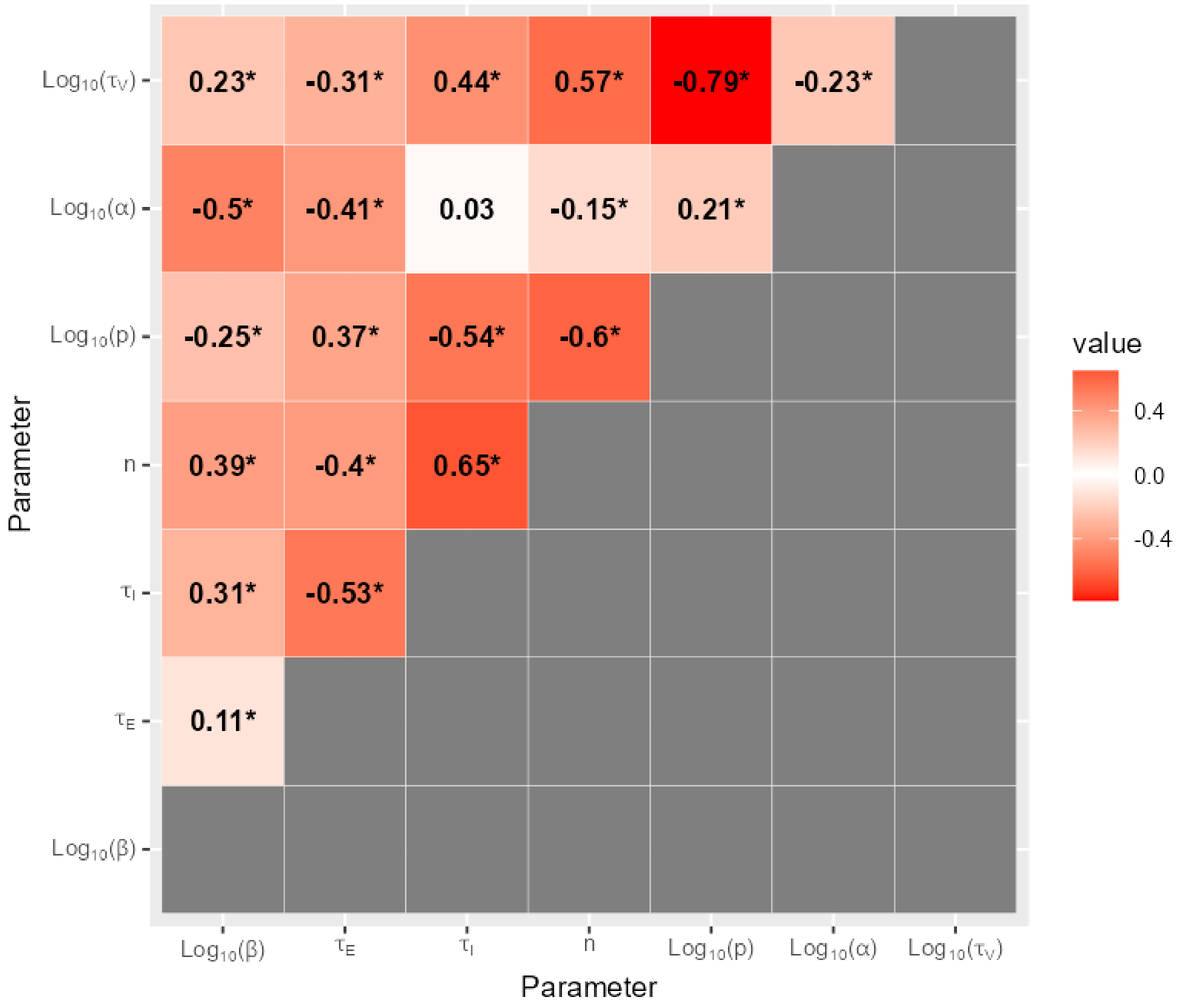

6.8 H5N1-C on DEF cells

Figure S26. The shifts in the distributions of model parameters during the ABC-SMC analysis of data from strain H5N1-C on DEF cells, when moving from the first particle population (containing accepted particles sampled from the prior distribution) to the posterior distribution via intermediate particle populations.

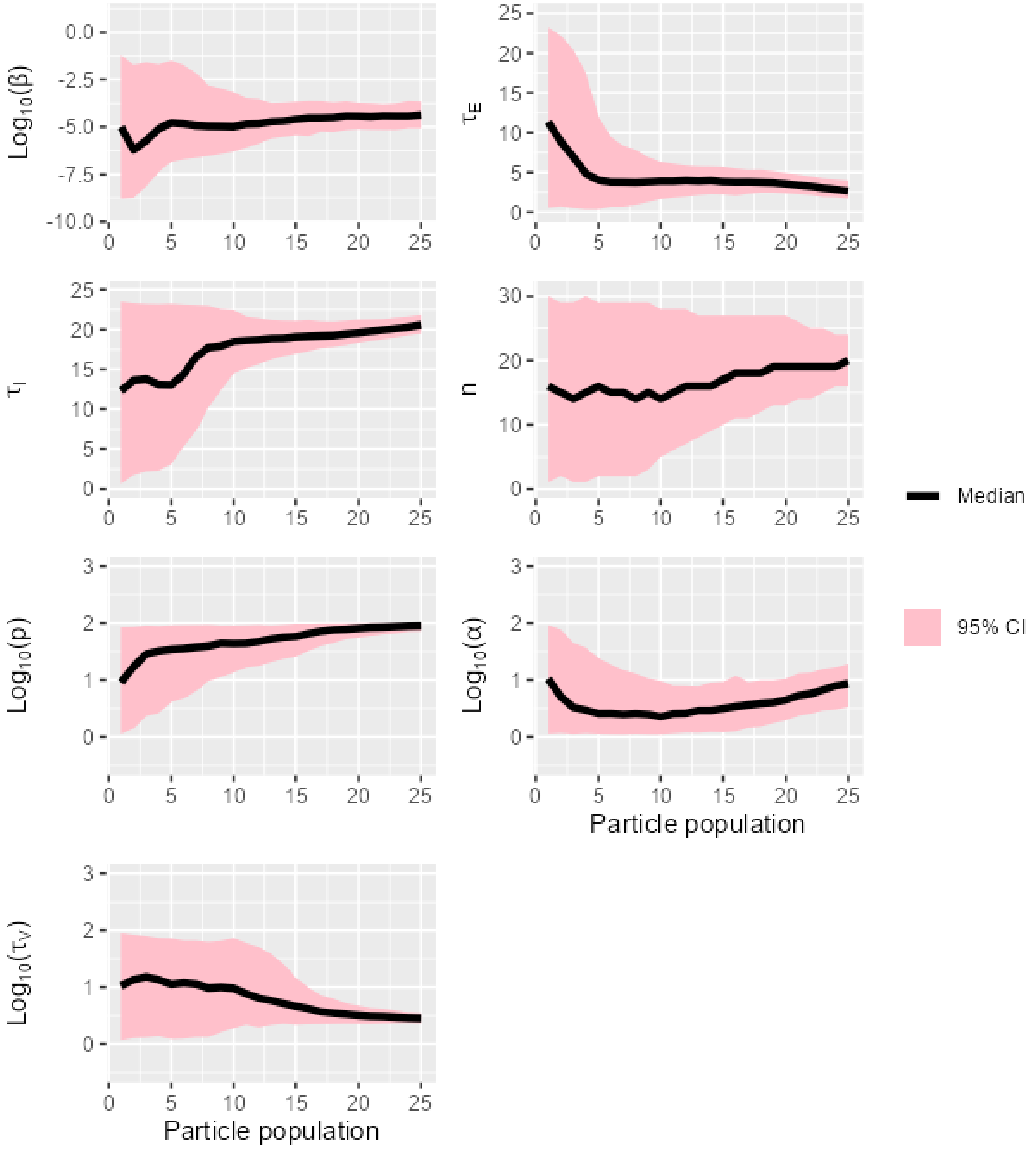

Figure S27. The overlap between prior and posterior distributions of model parameters for strain H5N1-C on DEF cells.

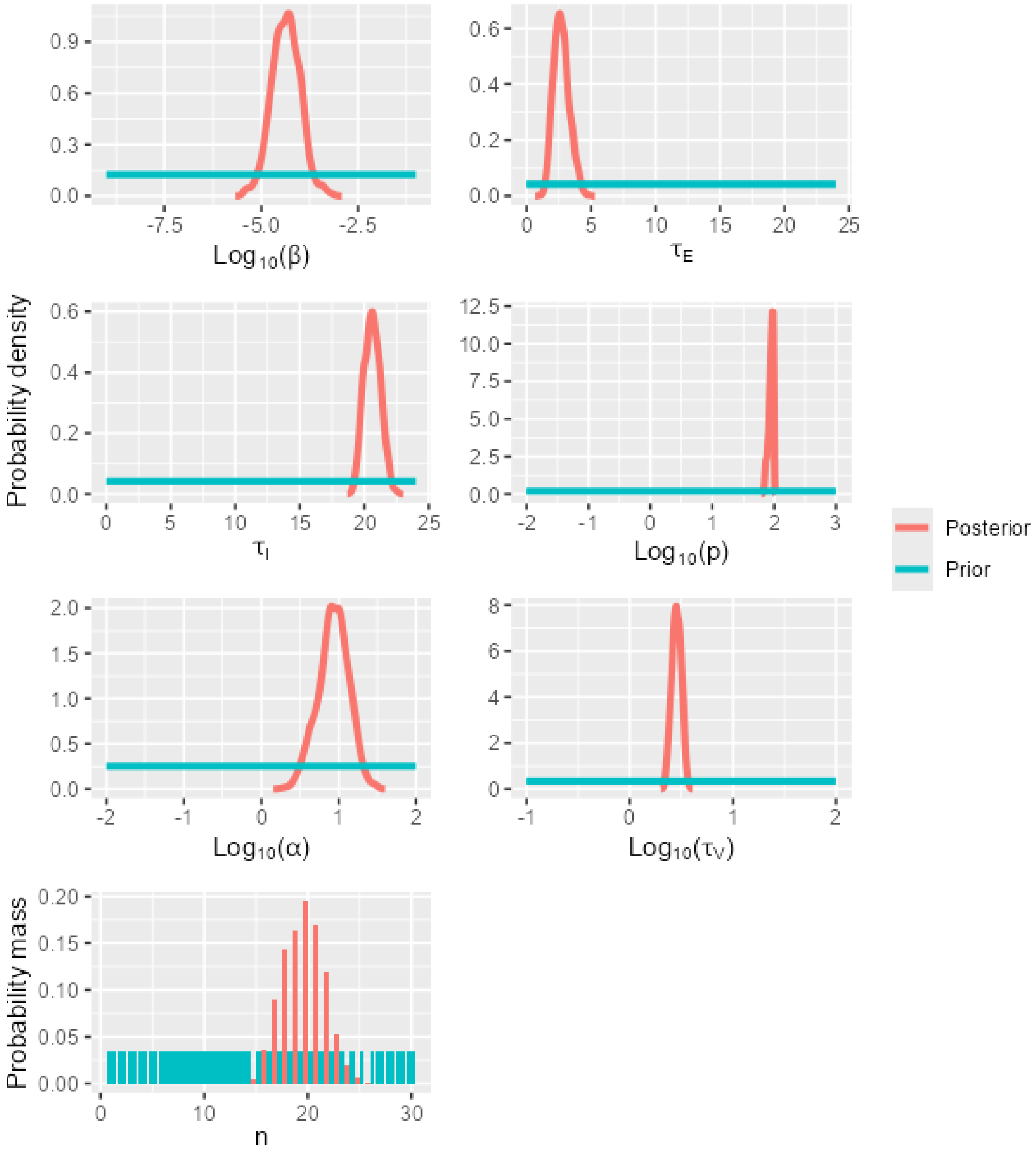

Figure S28. A matrix showing Kendall's rank correlation coefficient and the significance of the correlation (0.05 level, indicated by *) for pairs of model parameters calibrated using data for strain H5N1-C on DEF cells.

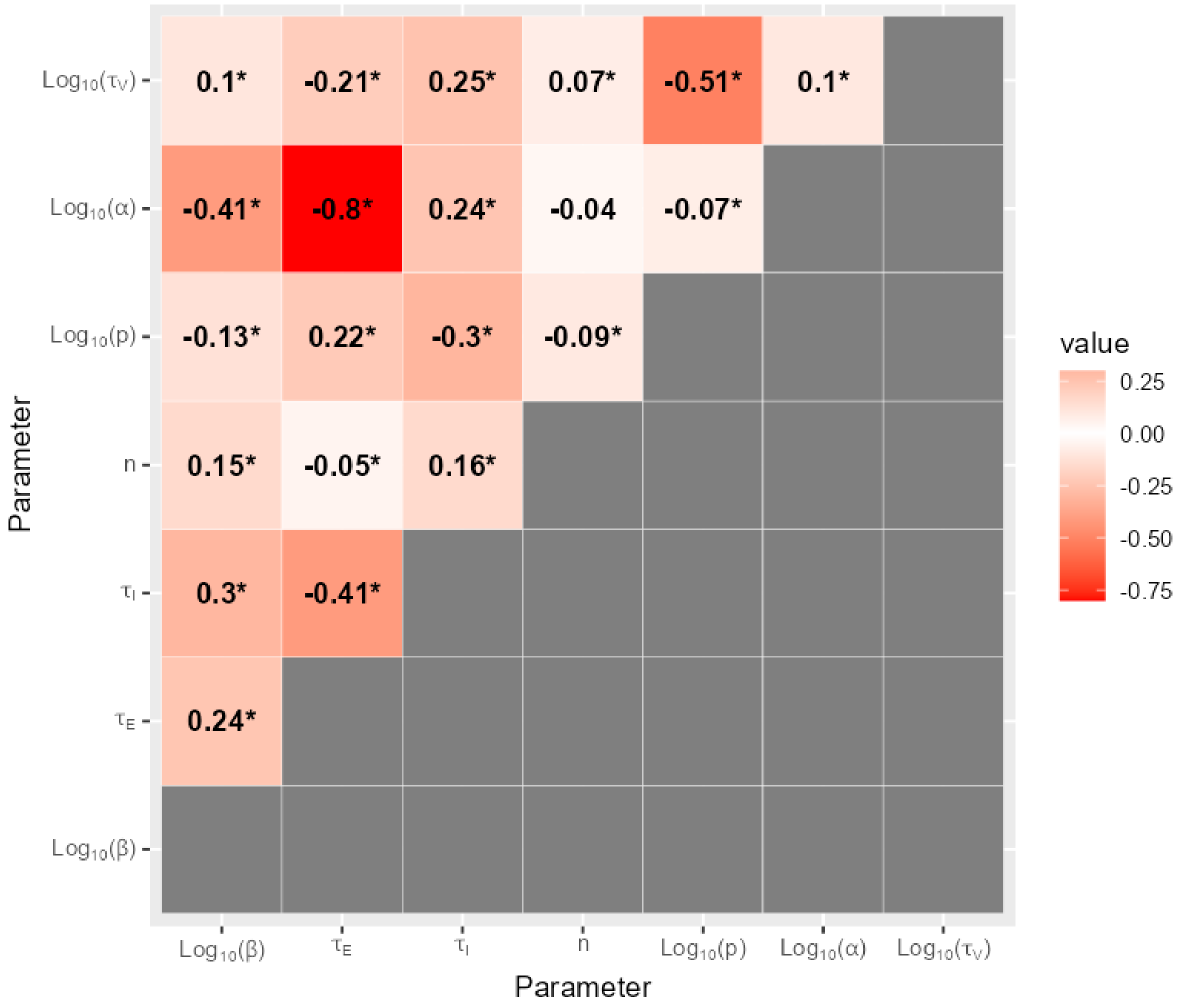

6.9 H5N6-2017 on CEF cells

Figure S29. The shifts in the distributions of model parameters during the ABC-SMC analysis of data from strain H5N6-2017 on CEF cells, when moving from the first particle population (containing accepted particles sampled from the prior distribution) to the posterior distribution via intermediate particle populations.

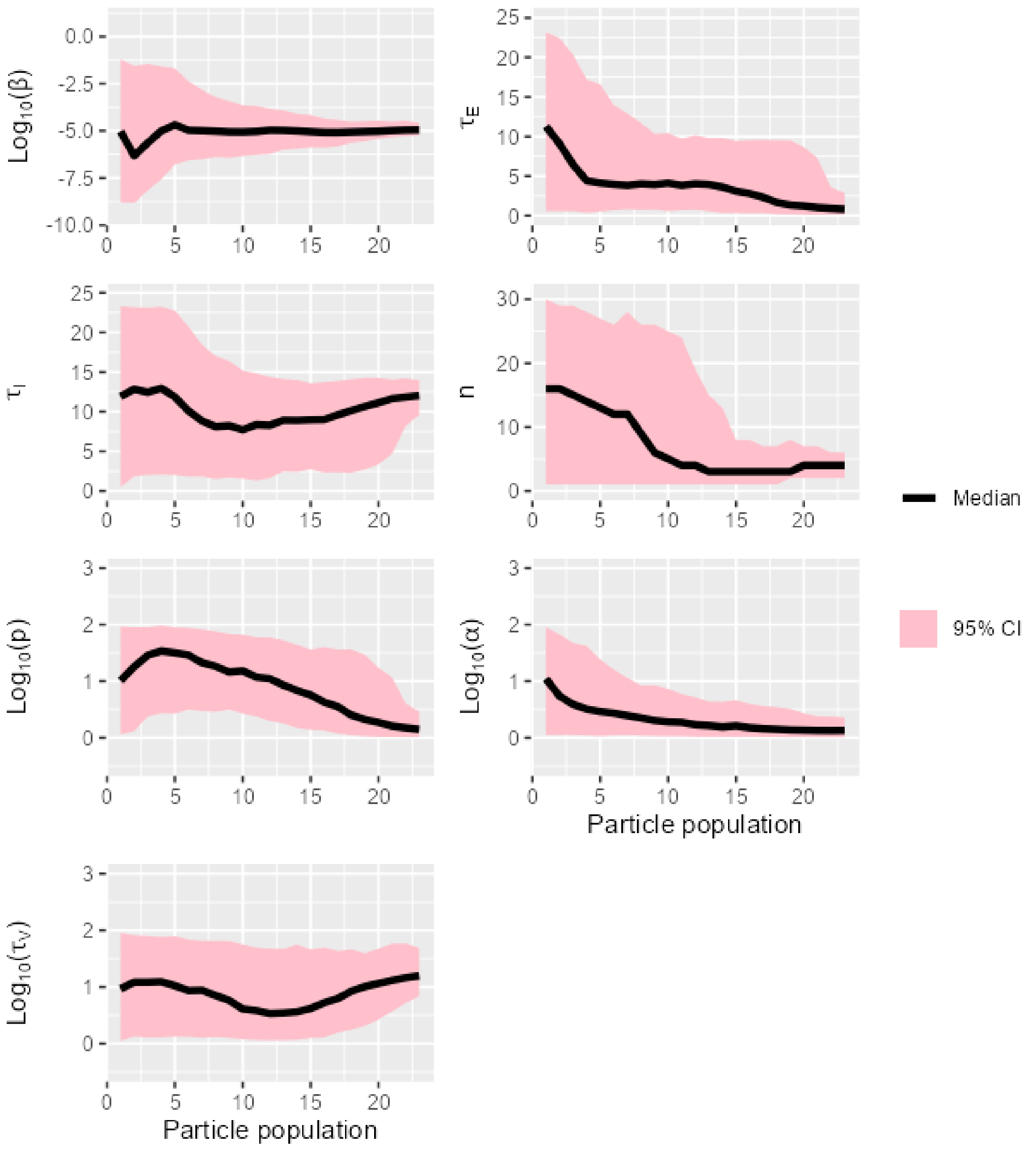

Figure S30. The overlap between prior and posterior distributions of model parameters for strain H5N6-2017 on CEF cells.

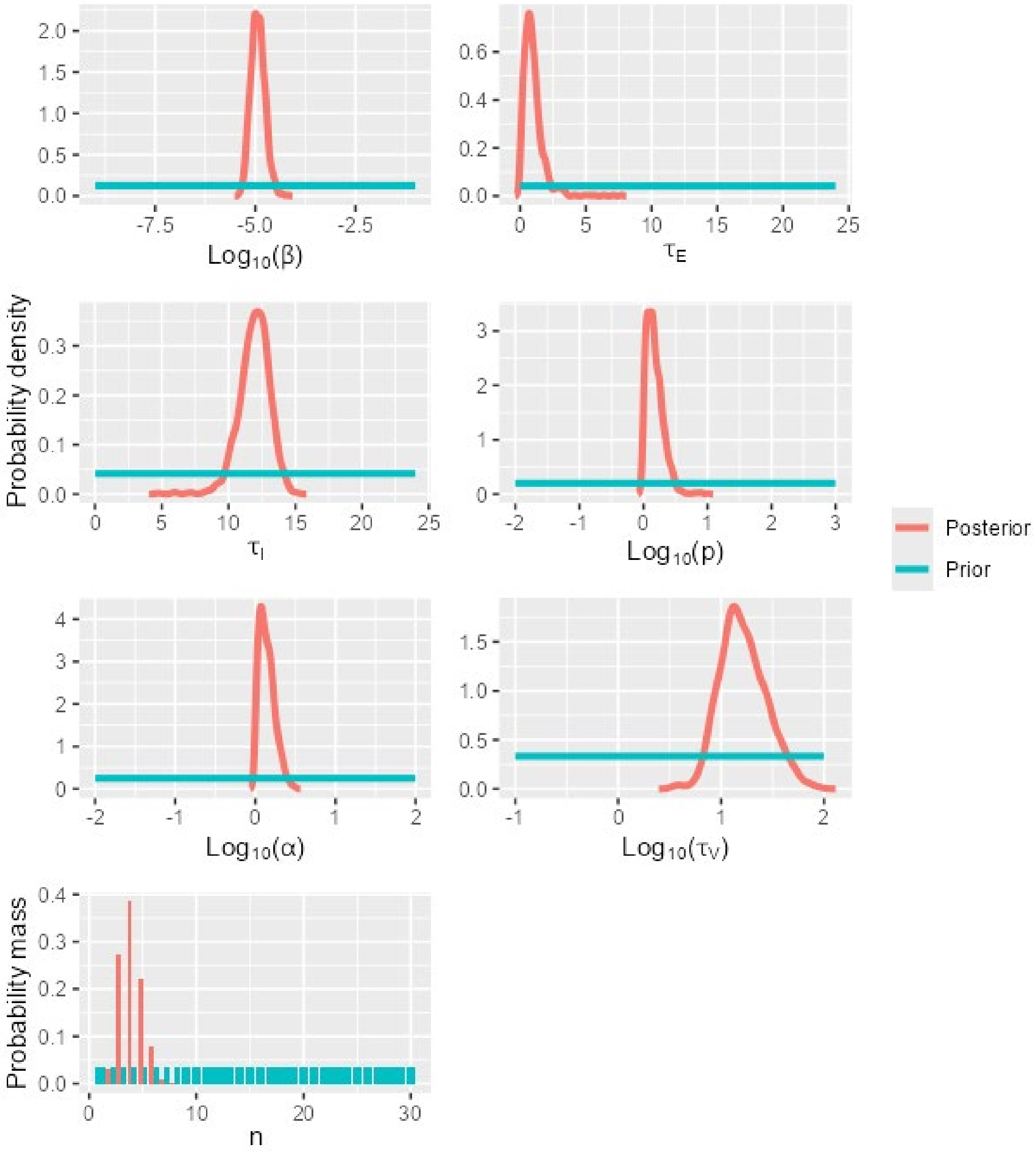

Figure S31. A matrix showing Kendall's rank correlation coefficient and the significance of the correlation (0.05 level, indicated by *) for pairs of model parameters calibrated using data for strain H5N6-2017 on CEF cells.

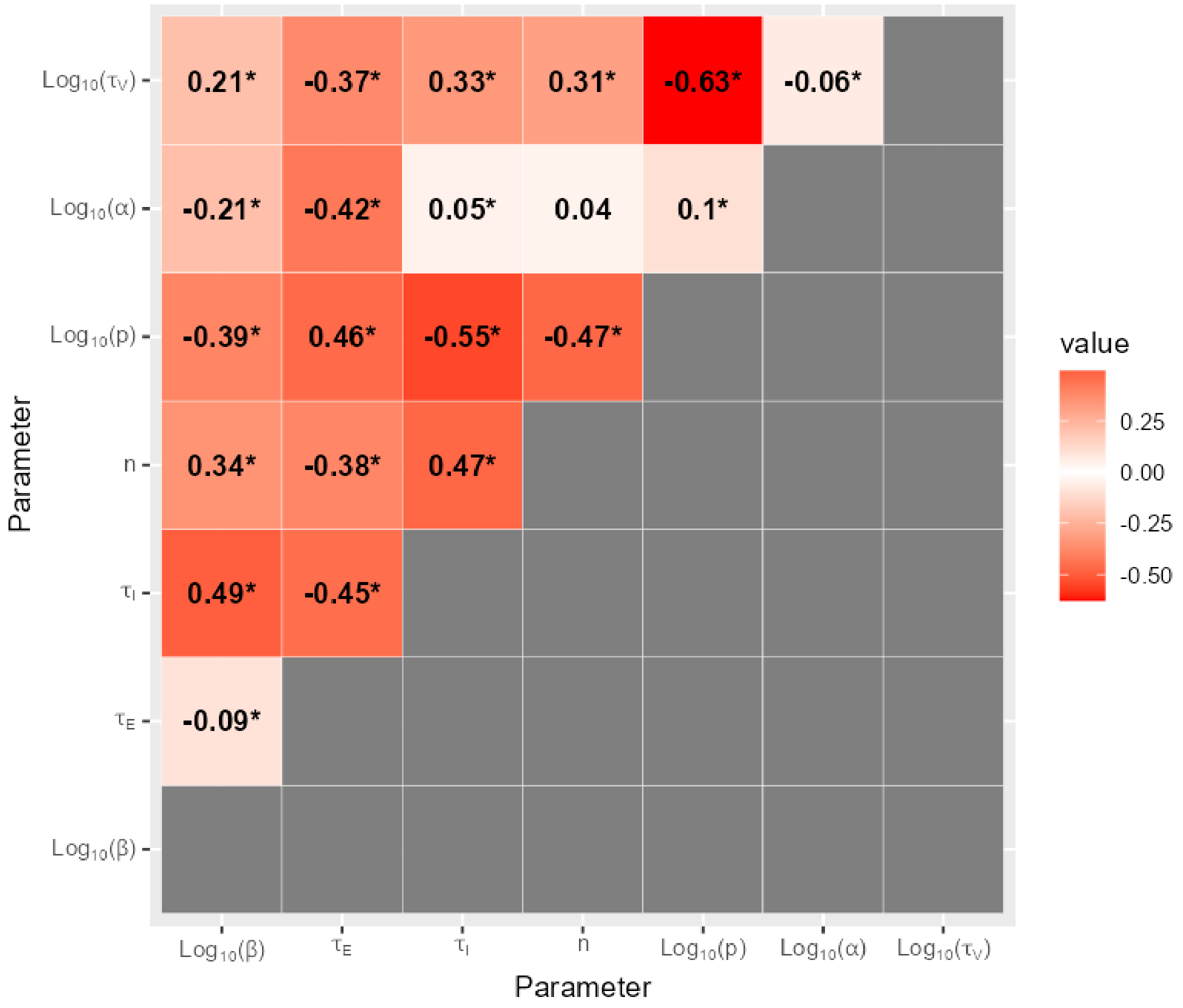

6.10 H5N6-2017 on DEF cells

Figure S32. The shifts in the distributions of model parameters during the ABC-SMC analysis of data from strain H5N6-2017 on DEF cells, when moving from the first particle population (containing accepted particles sampled from the prior distribution) to the posterior distribution via intermediate particle populations.

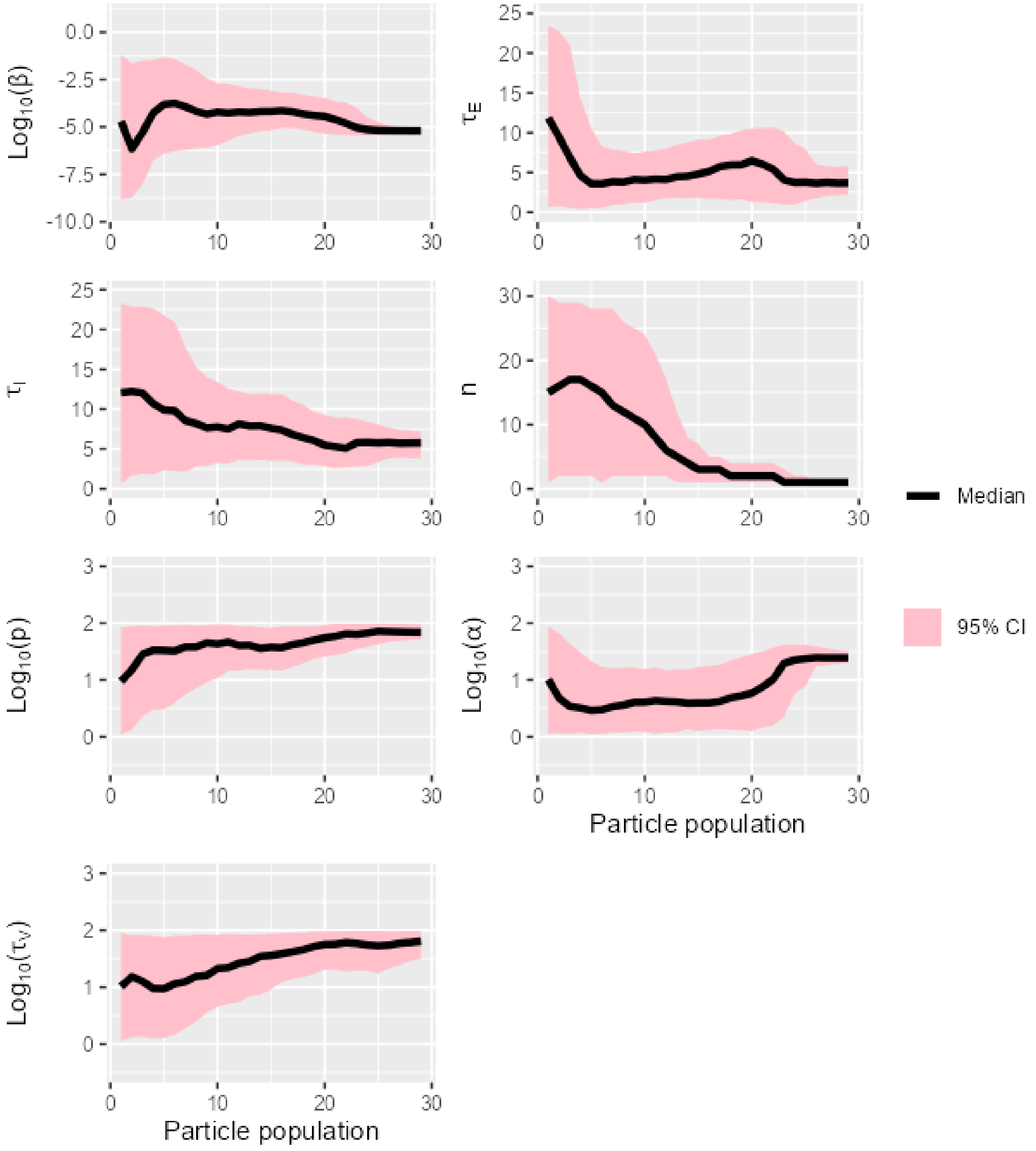

Figure S33. The overlap between prior and posterior distributions of model parameters for strain H5N6-2017 on DEF cells.

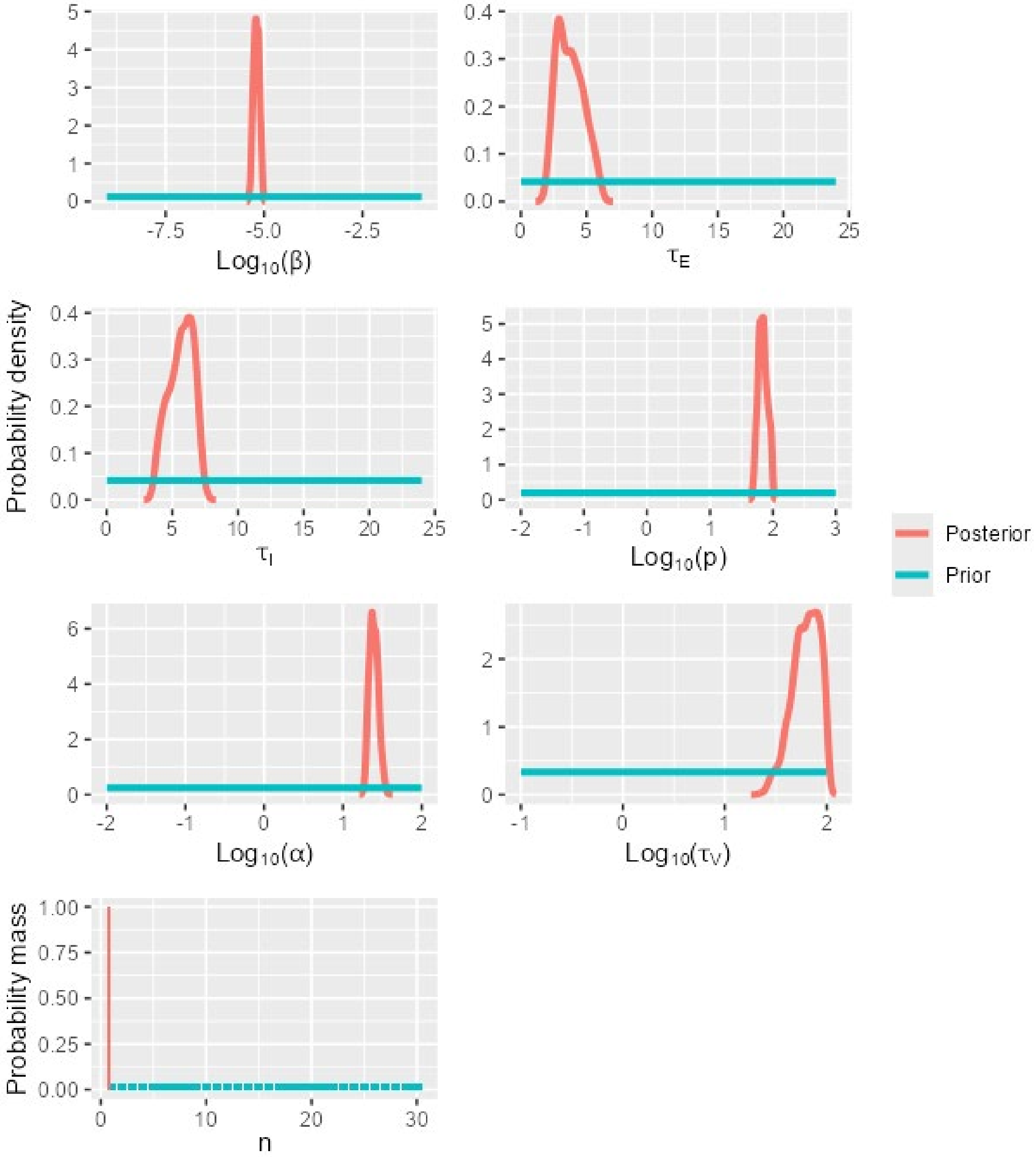

Figure S34. A matrix showing Kendall's rank correlation coefficient and the significance of the correlation (0.05 level, indicated by *) for pairs of model parameters calibrated using data for strain H5N6-2017 on DEF cells.

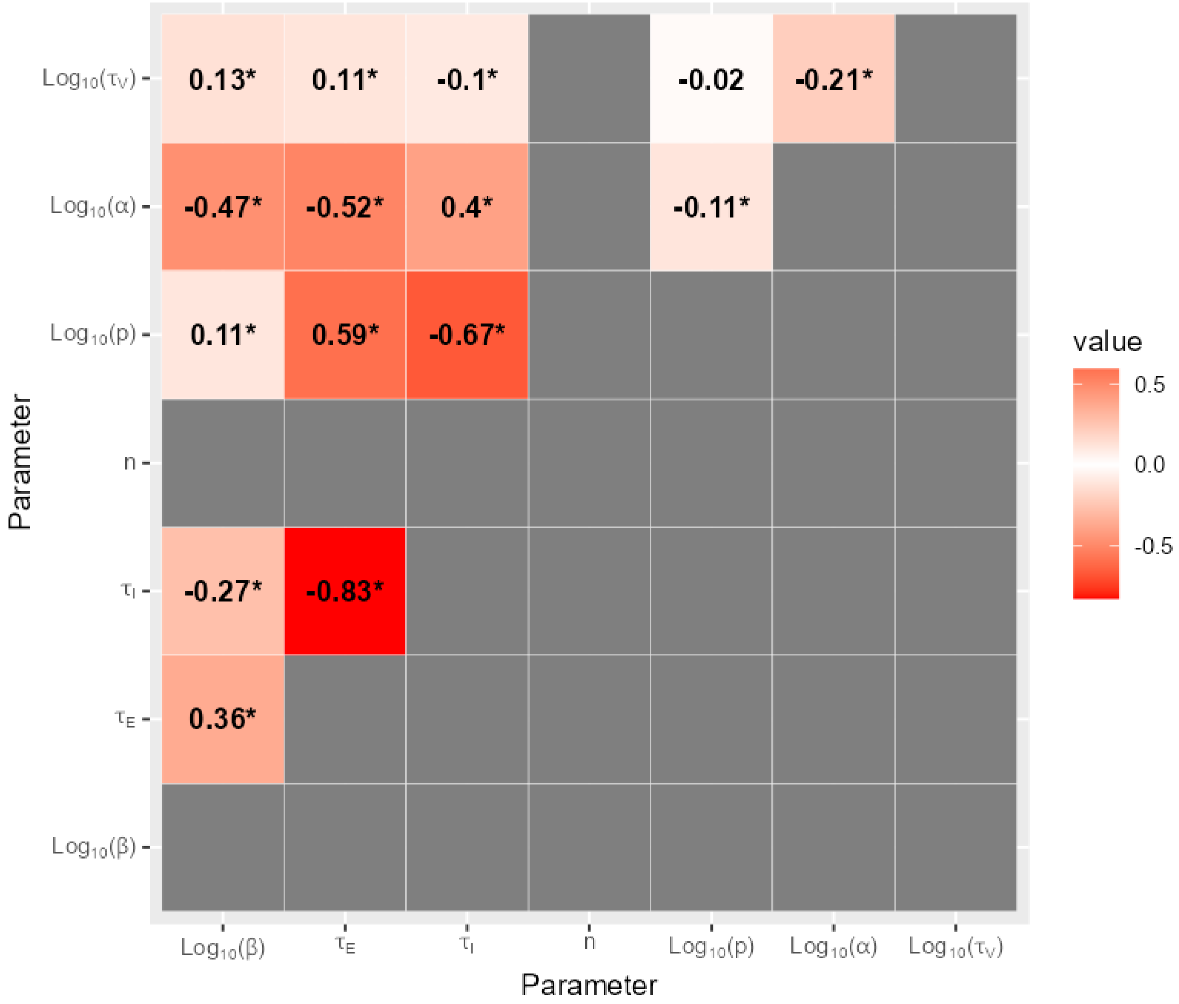

6.11 H5N8-2014 on CEF cells

Figure S35. The shifts in the distributions of model parameters during the ABC-SMC analysis of data from strain H5N8-2014 on CEF cells, when moving from the first particle population (containing accepted particles sampled from the prior distribution) to the posterior distribution via intermediate particle populations.

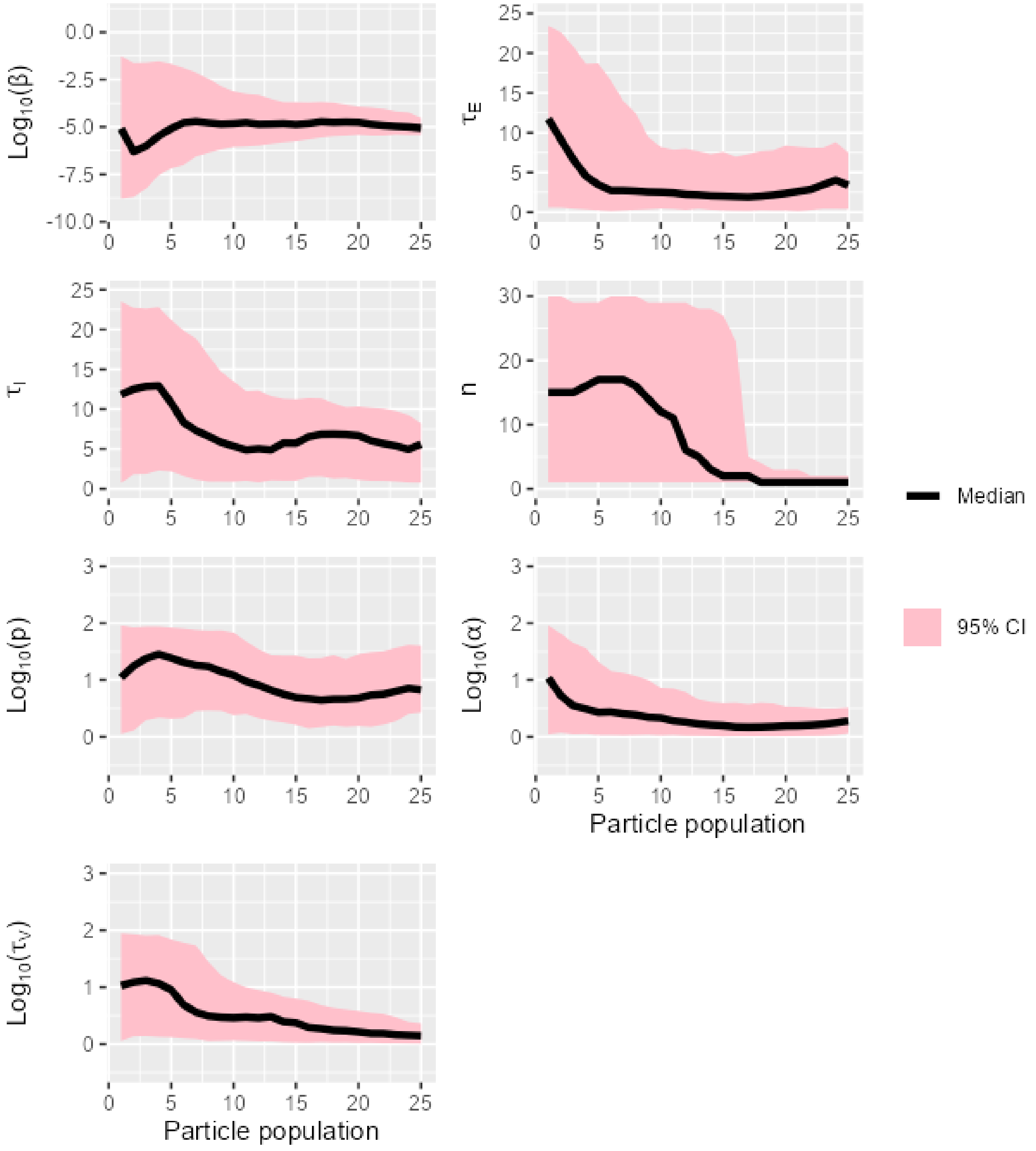

Figure S36. The overlap between prior and posterior distributions of model parameters for strain H5N8-2014 on CEF cells.

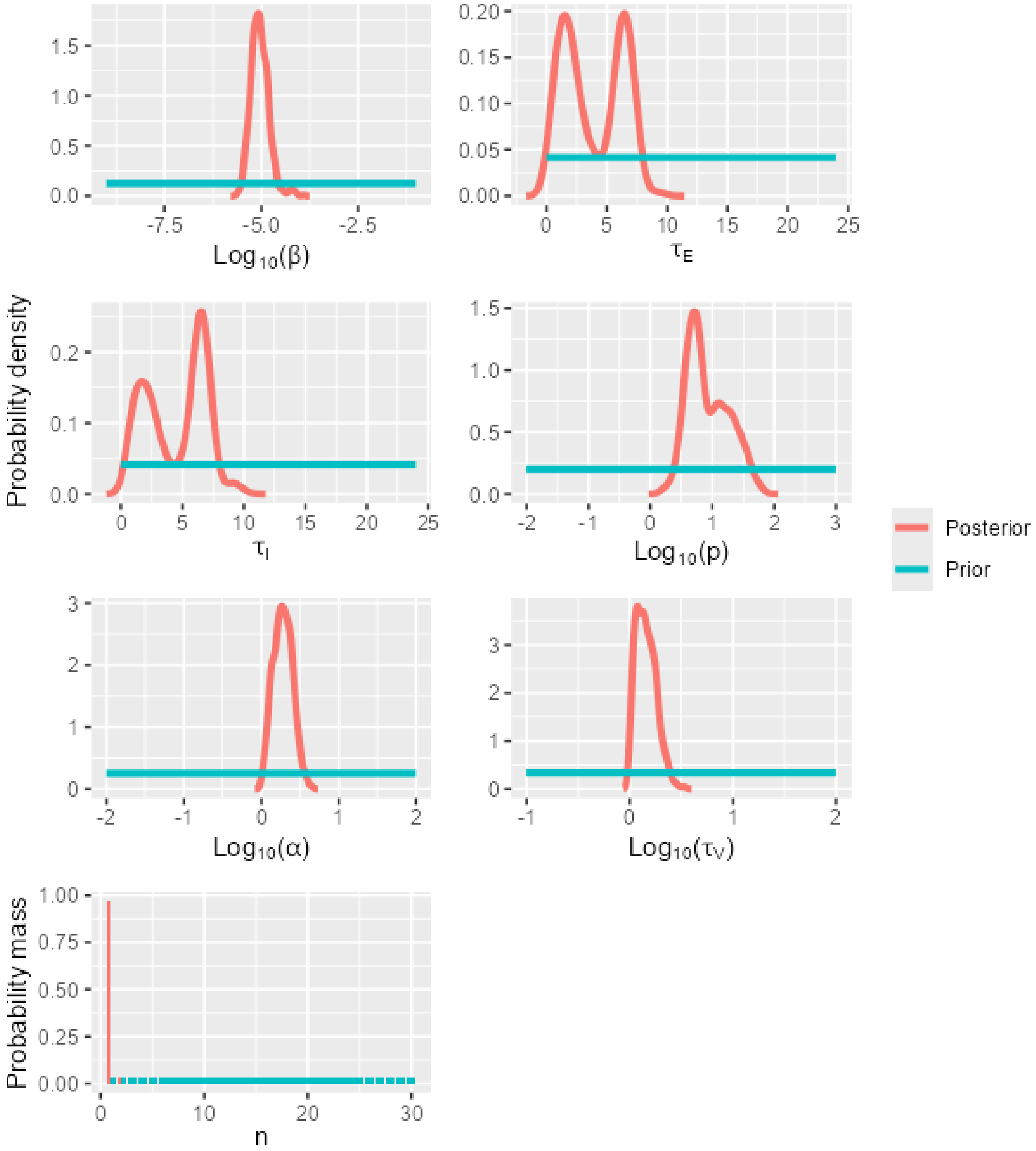

Figure S37. A matrix showing Kendall's rank correlation coefficient and the significance of the correlation (0.05 level, indicated by *) for pairs of model parameters calibrated using data for strain H5N8-2016 on CEF cells.

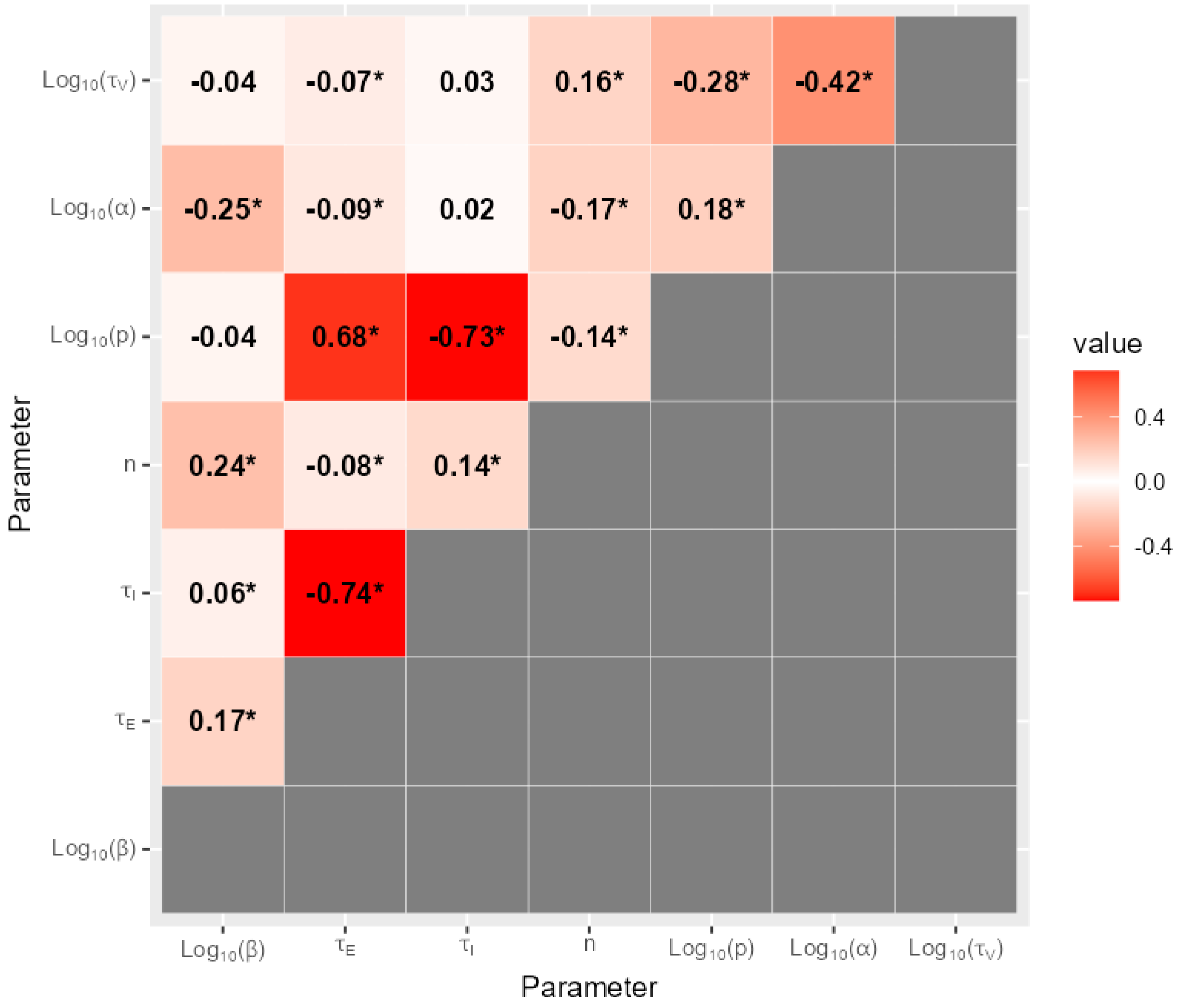

6.12 H5N8-2014 on DEF cells

Figure S38. The shifts in the distributions of model parameters during the ABC-SMC analysis of data from strain H5N8-2014 on DEF cells, when moving from the first particle population (containing accepted particles sampled from the prior distribution) to the posterior distribution via intermediate particle populations.

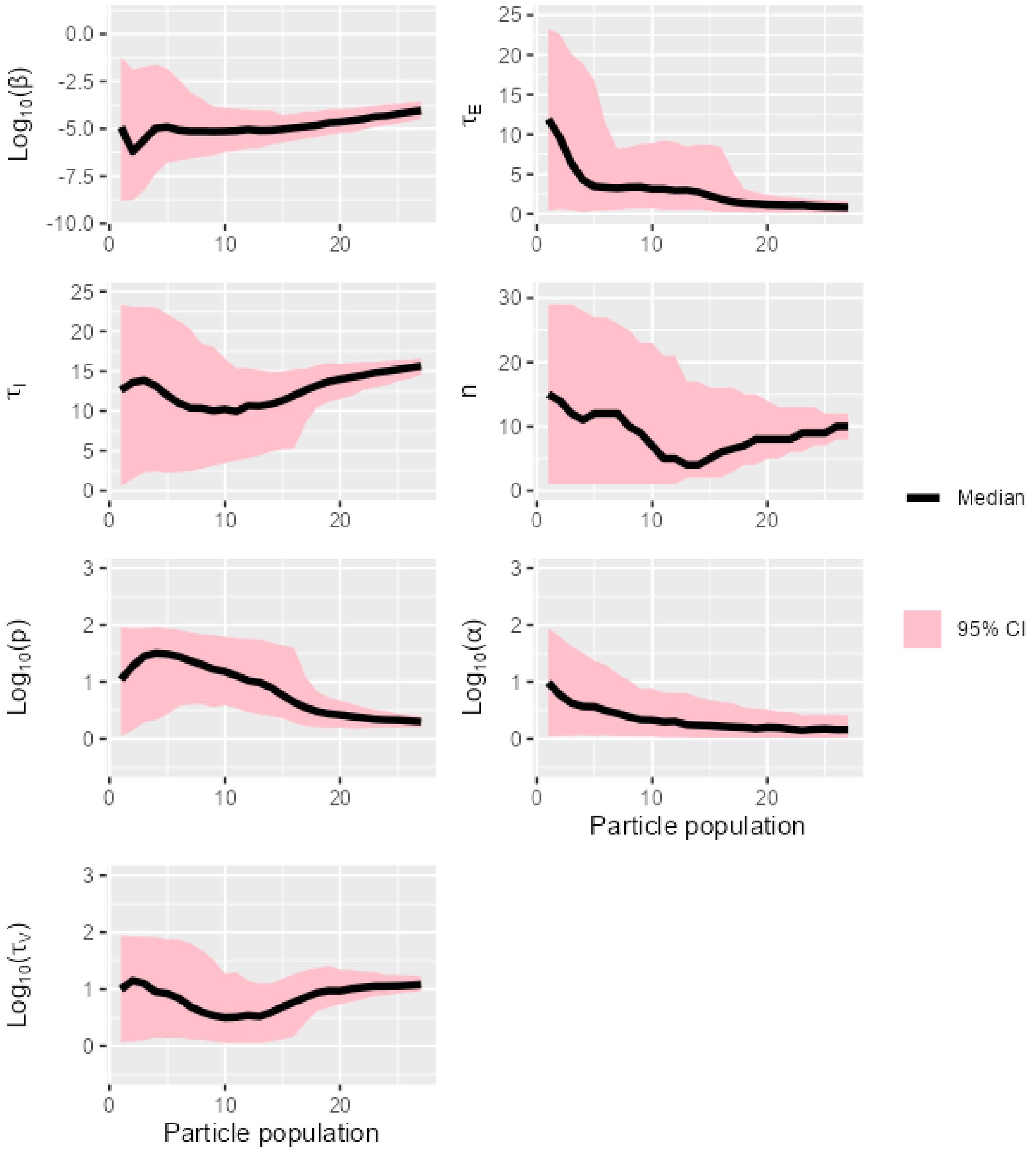

Figure S39. The overlap between prior and posterior distributions of model parameters for strain H5N8-2014 on DEF cells.

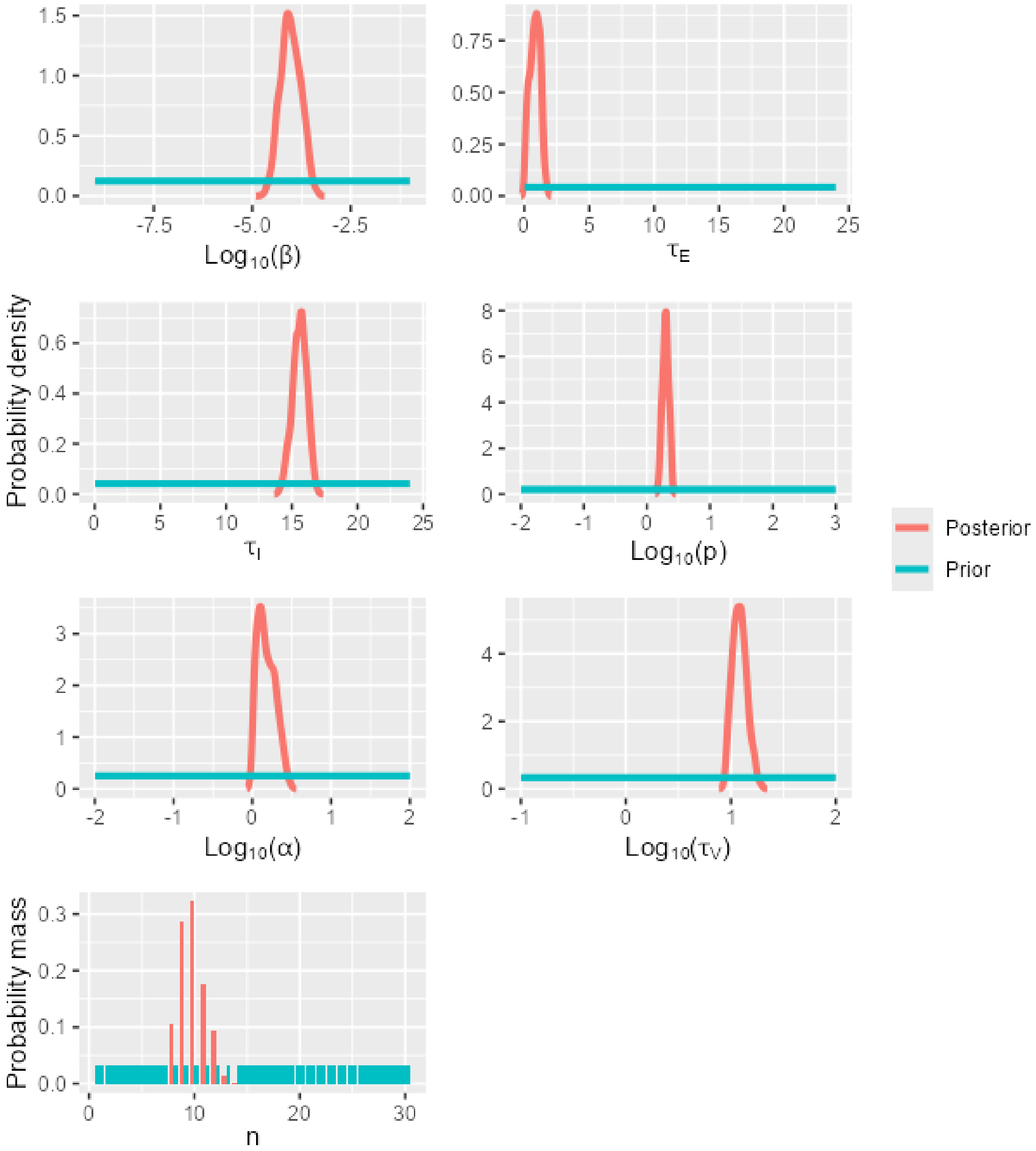

Figure S40. A matrix showing Kendall's rank correlation coefficient and the significance of the correlation (0.05 level, indicated by *) for pairs of model parameters calibrated using data for strain H5N8-2014 on DEF cells.

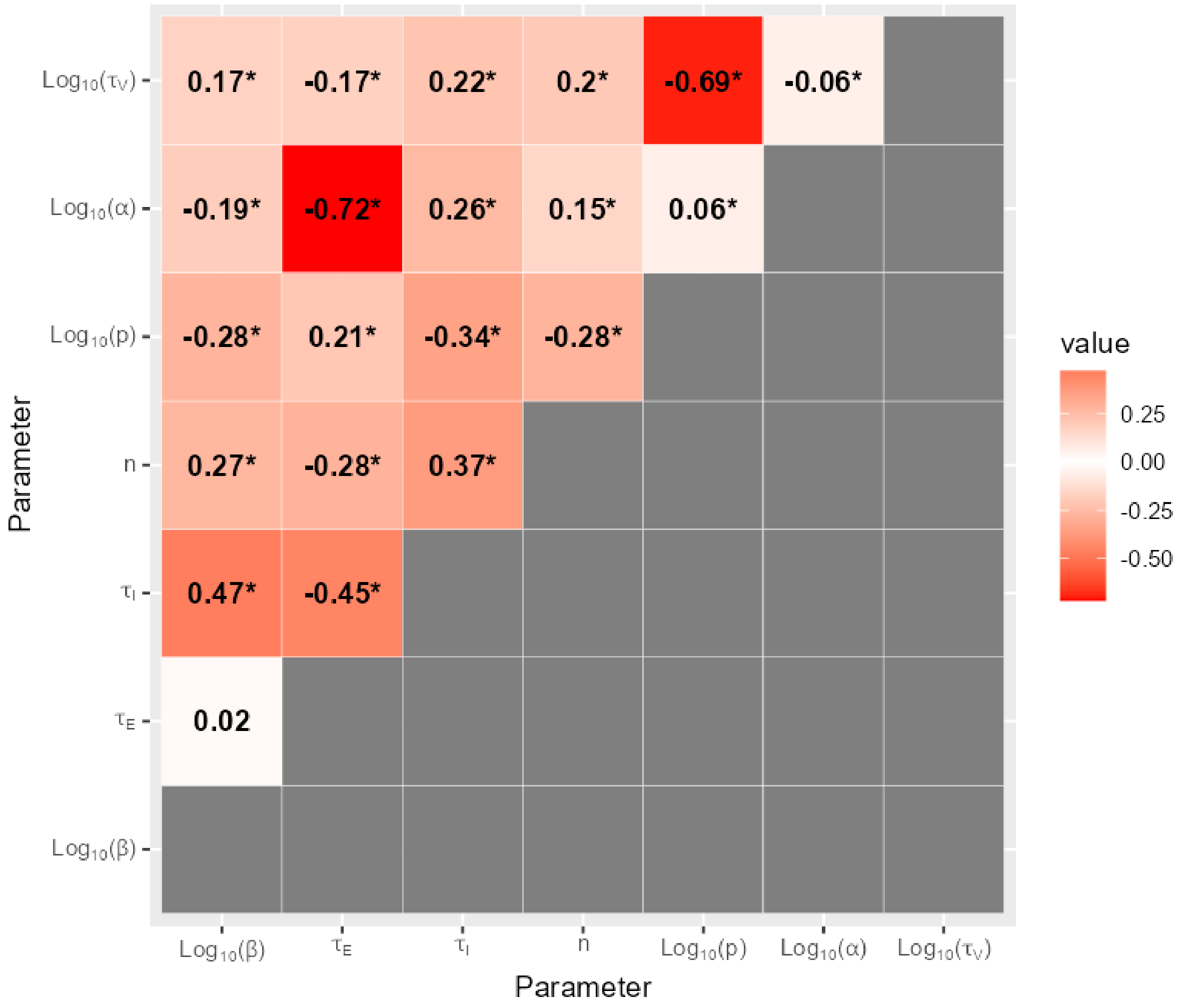

6.13 H5N8-2016 on CEF cells

Figure S41. The shifts in the distributions of model parameters during the ABC-SMC analysis of data from strain H5N8-2016 on CEF cells, when moving from the first particle population (containing accepted particles sampled from the prior distribution) to the posterior distribution via intermediate particle populations.

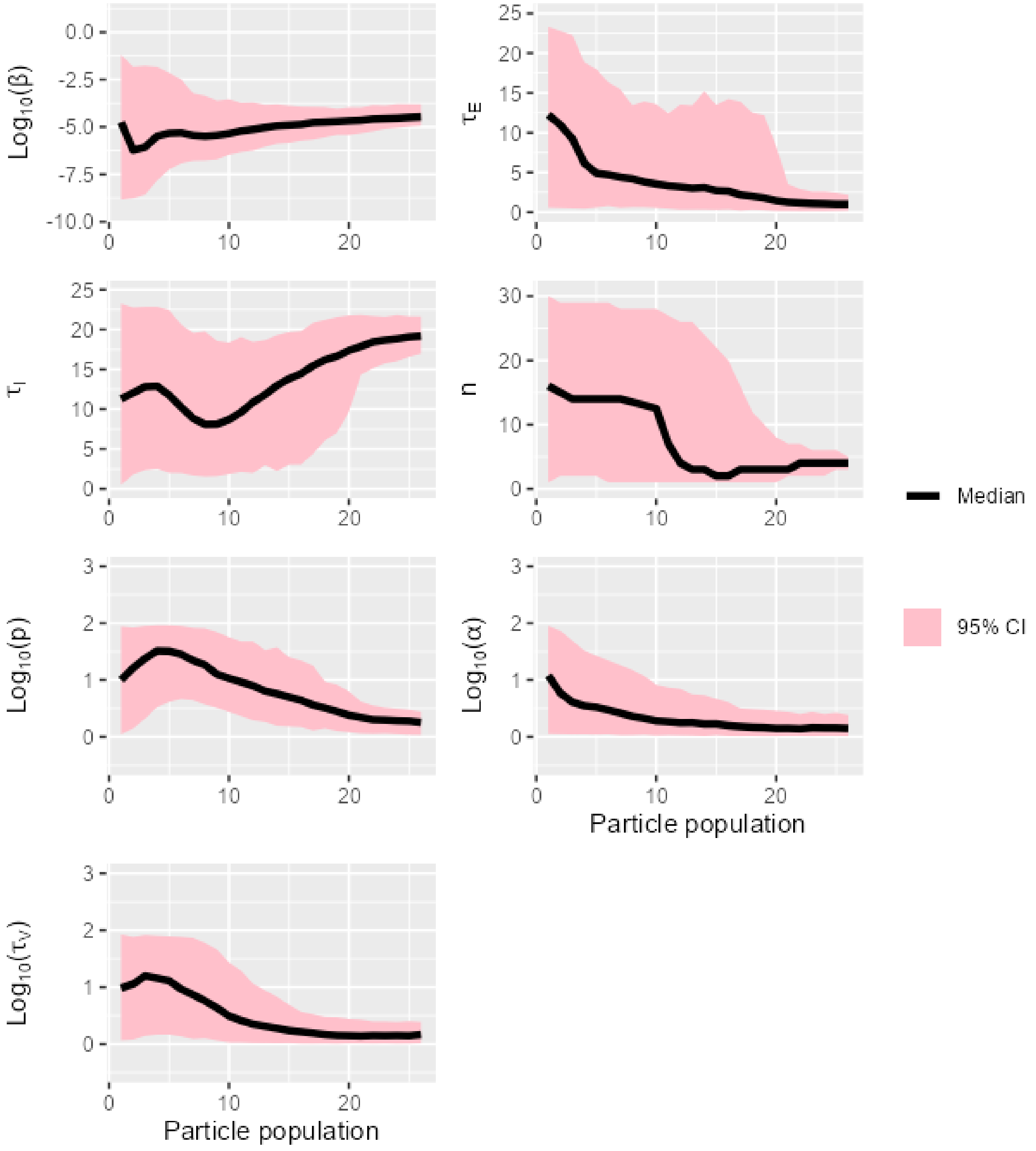

Figure S42. The overlap between prior and posterior distributions of model parameters for strain H5N8-2016 on CEF cells.

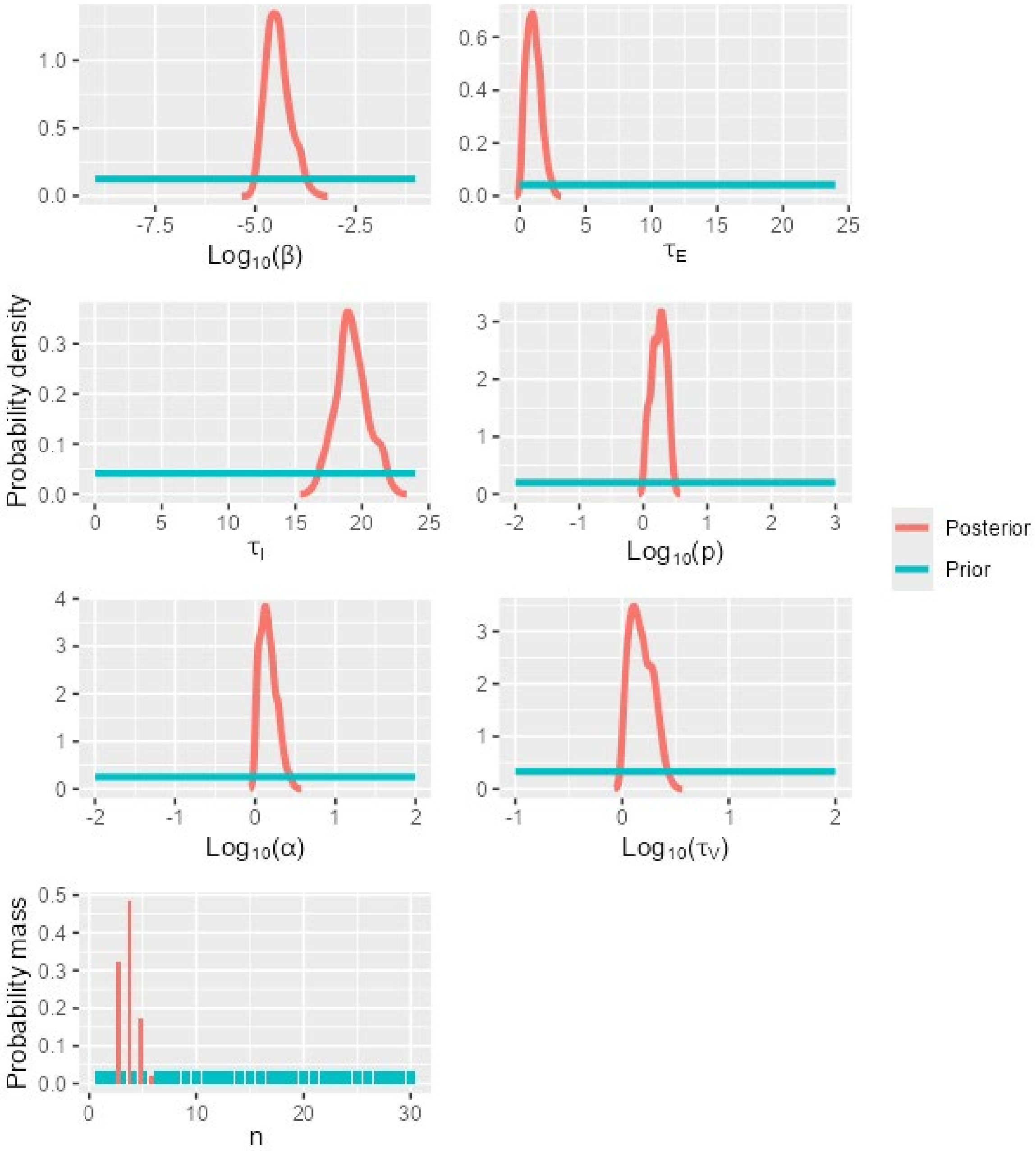

Figure S43. A matrix showing Kendall's rank correlation coefficient and the significance of the correlation (0.05 level, indicated by *) for pairs of model parameters calibrated using data for strain H5N8-2016 on CEF cells.

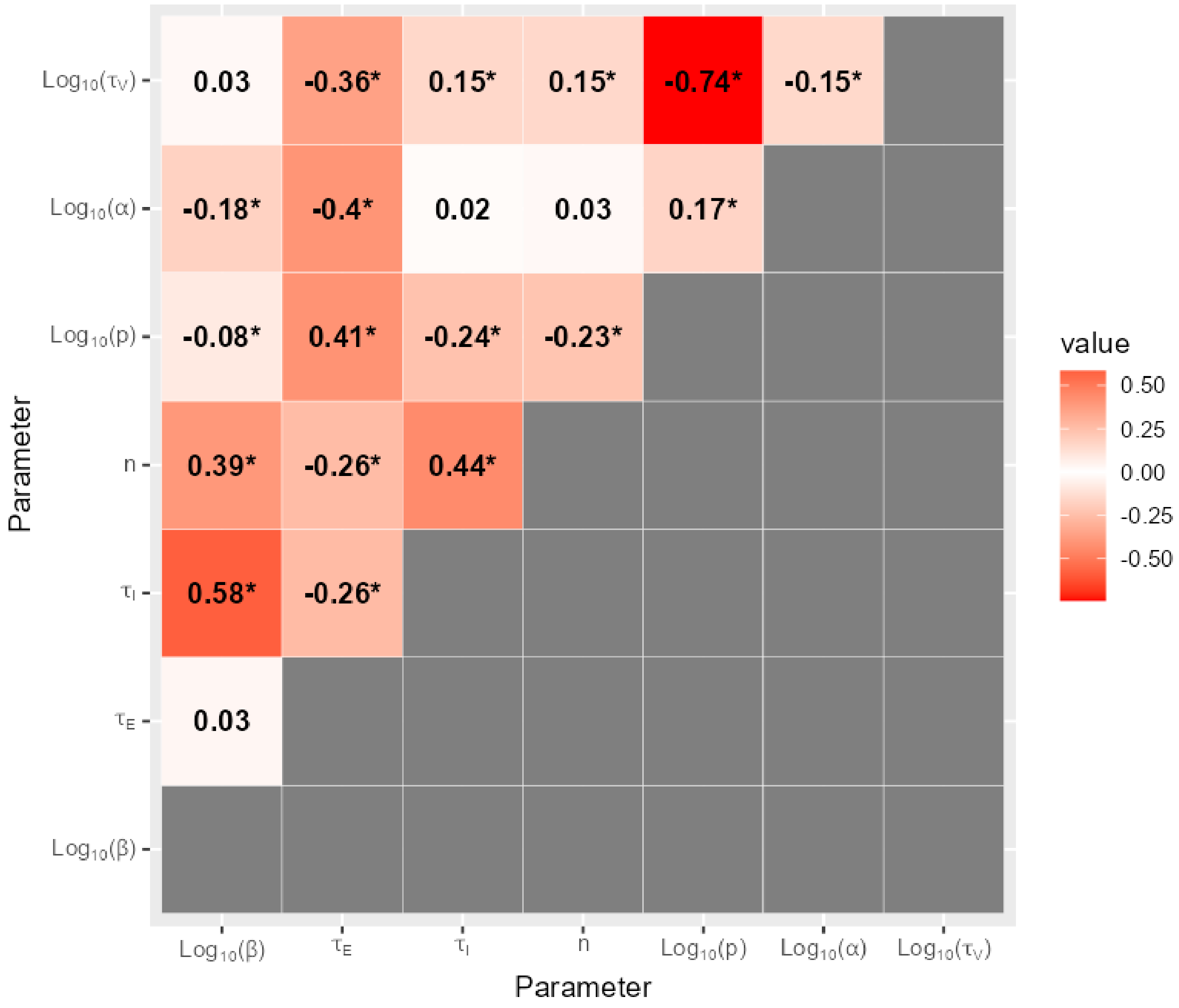

6.14 H5N8-2016 on DEF cells

Figure S44. The shifts in the distributions of model parameters during the ABC-SMC analysis of data from strain H5N8-2016 on DEF cells, when moving from the first particle population (containing accepted particles sampled from the prior distribution) to the posterior distribution via intermediate particle populations.

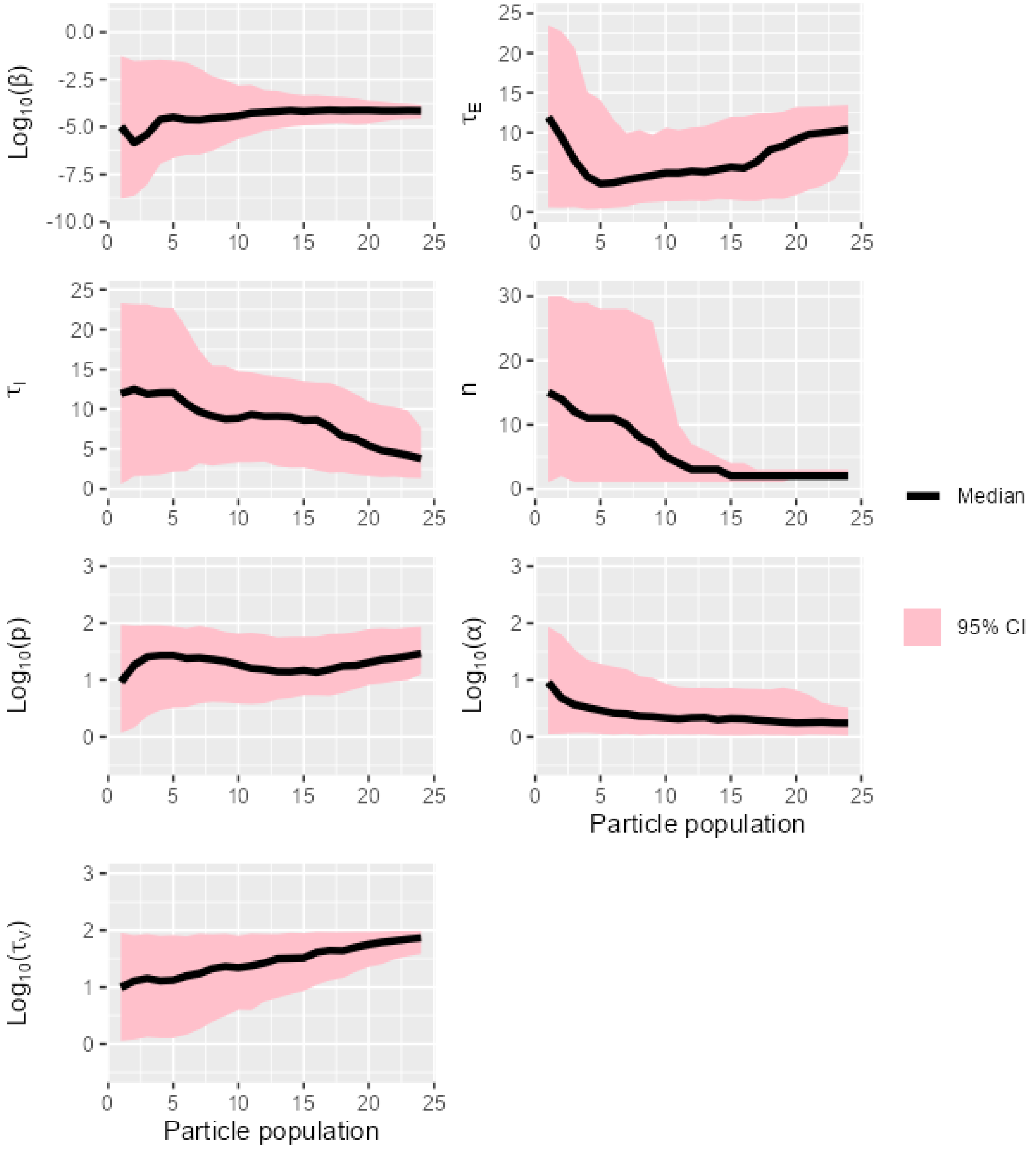

Figure S45. The overlap between prior and posterior distributions of model parameters for strain H5N8-2016 on DEF cells.

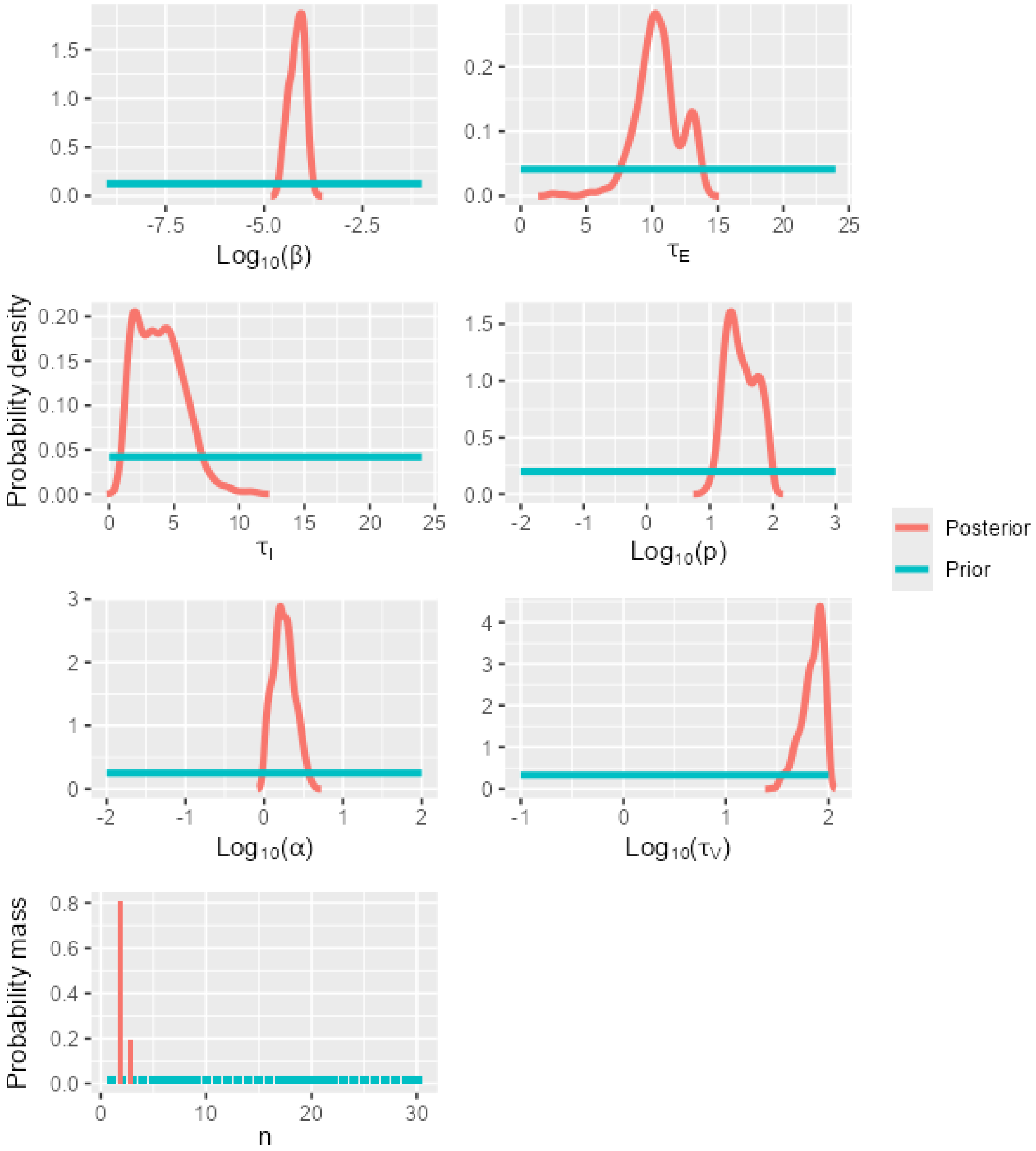

Figure S46. A matrix showing Kendall's rank correlation coefficient and the significance of the correlation (0.05 level, indicated by *) for pairs of model parameters calibrated using data for strain H5N8-2016 on DEF cells.

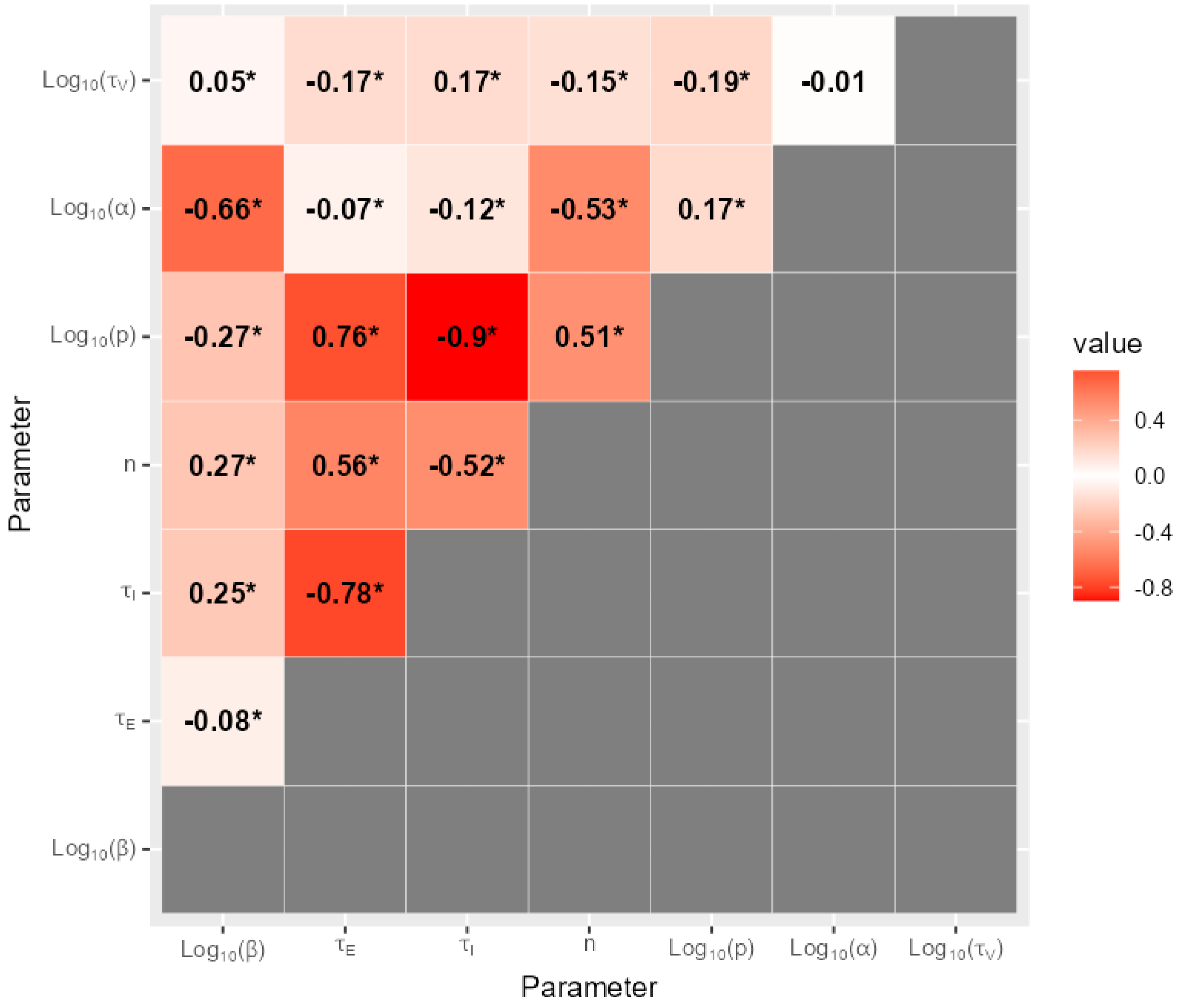

6.15 H5N8-2020 on CEF cells

Figure S47. The shifts in the distributions of model parameters during the ABC-SMC analysis of data from strain H5N8-2020 on CEF cells, when moving from the first particle population (containing accepted particles sampled from the prior distribution) to the posterior distribution via intermediate particle populations.

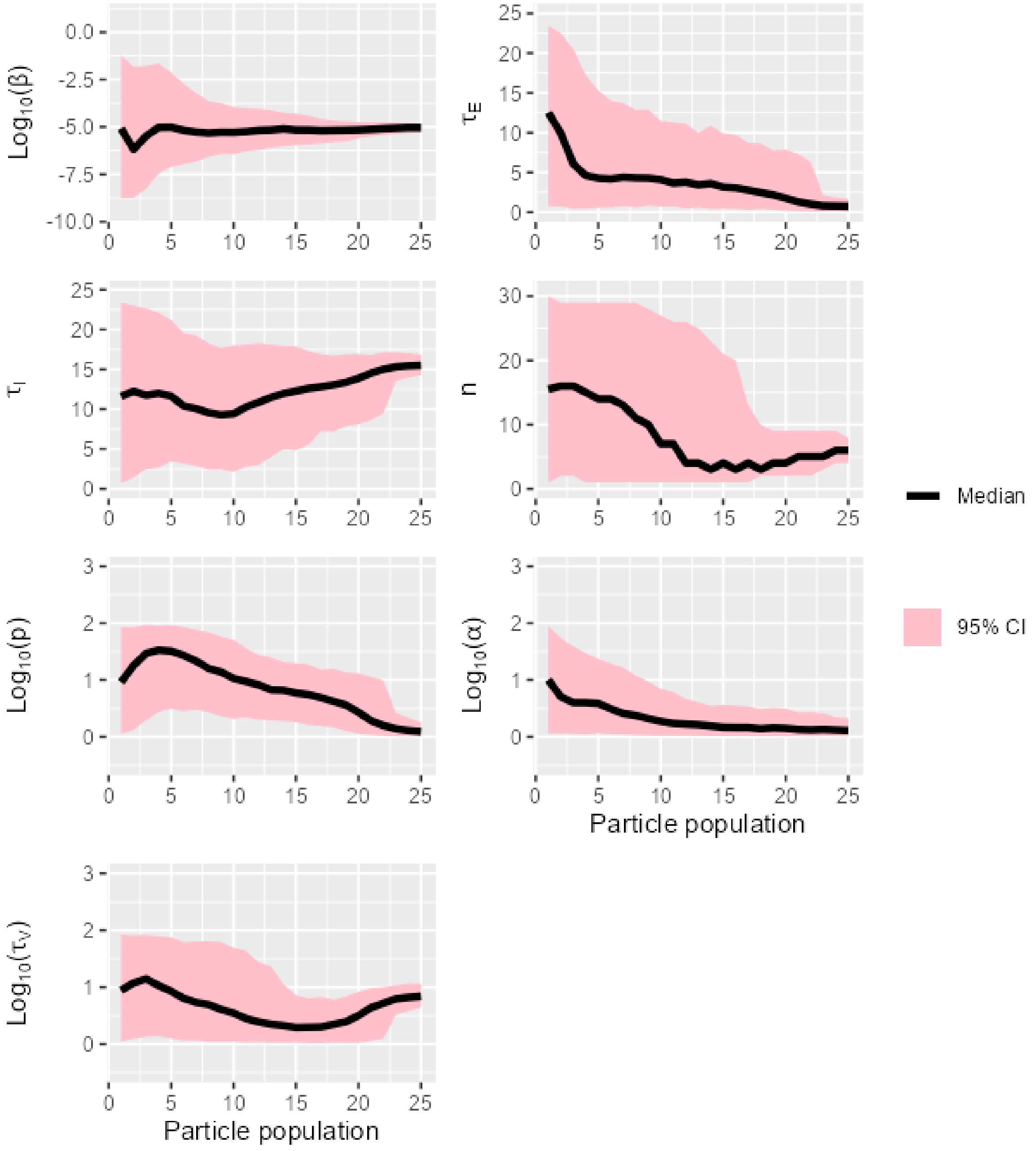

Figure S48. The overlap between prior and posterior distributions of model parameters for strain H5N8-2020 on CEF cells.

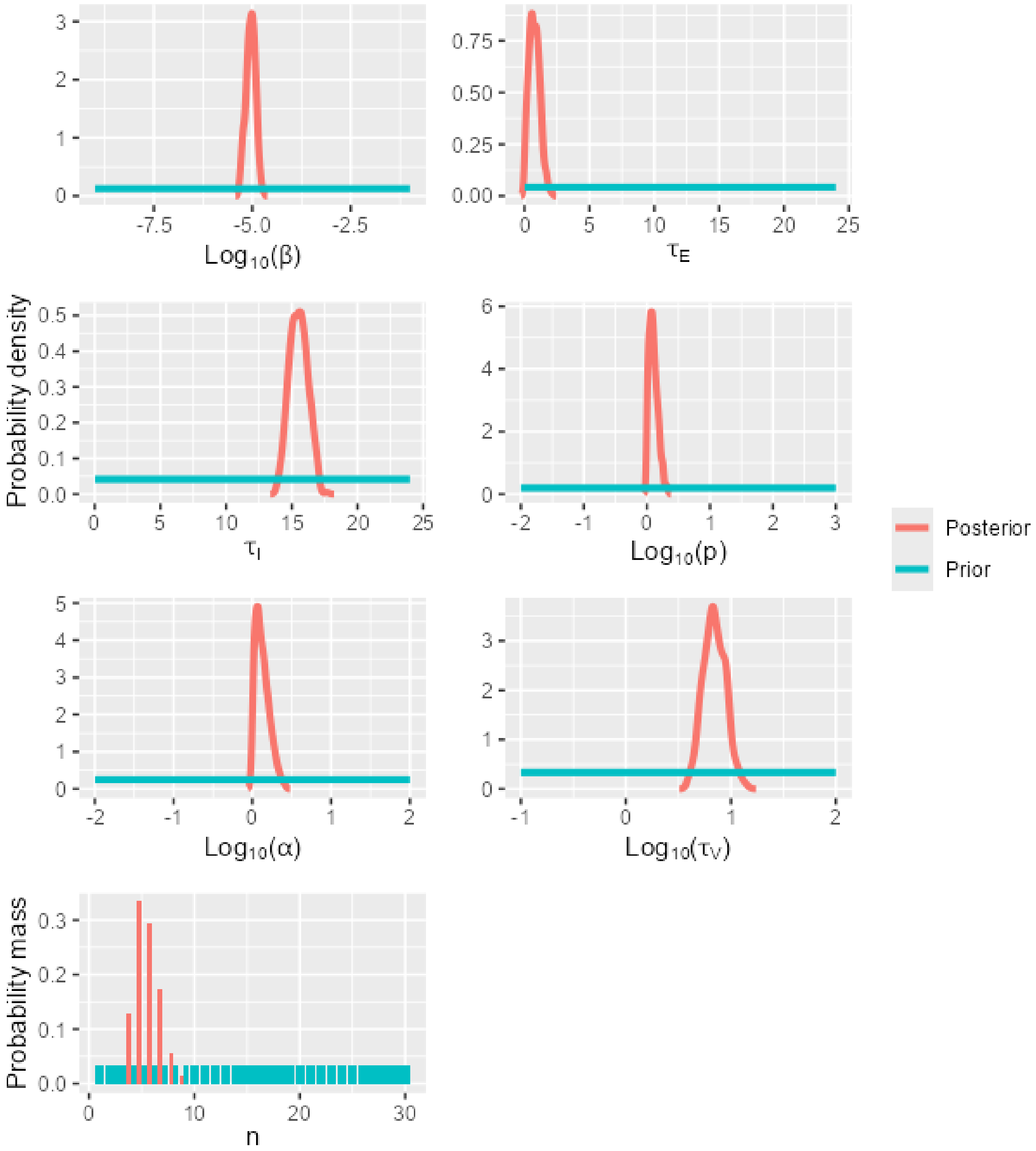

Figure S49. A matrix showing Kendall's rank correlation coefficient and the significance of the correlation (0.05 level, indicated by *) for pairs of model parameters calibrated using data for strain H5N8-2020 on CEF cells.

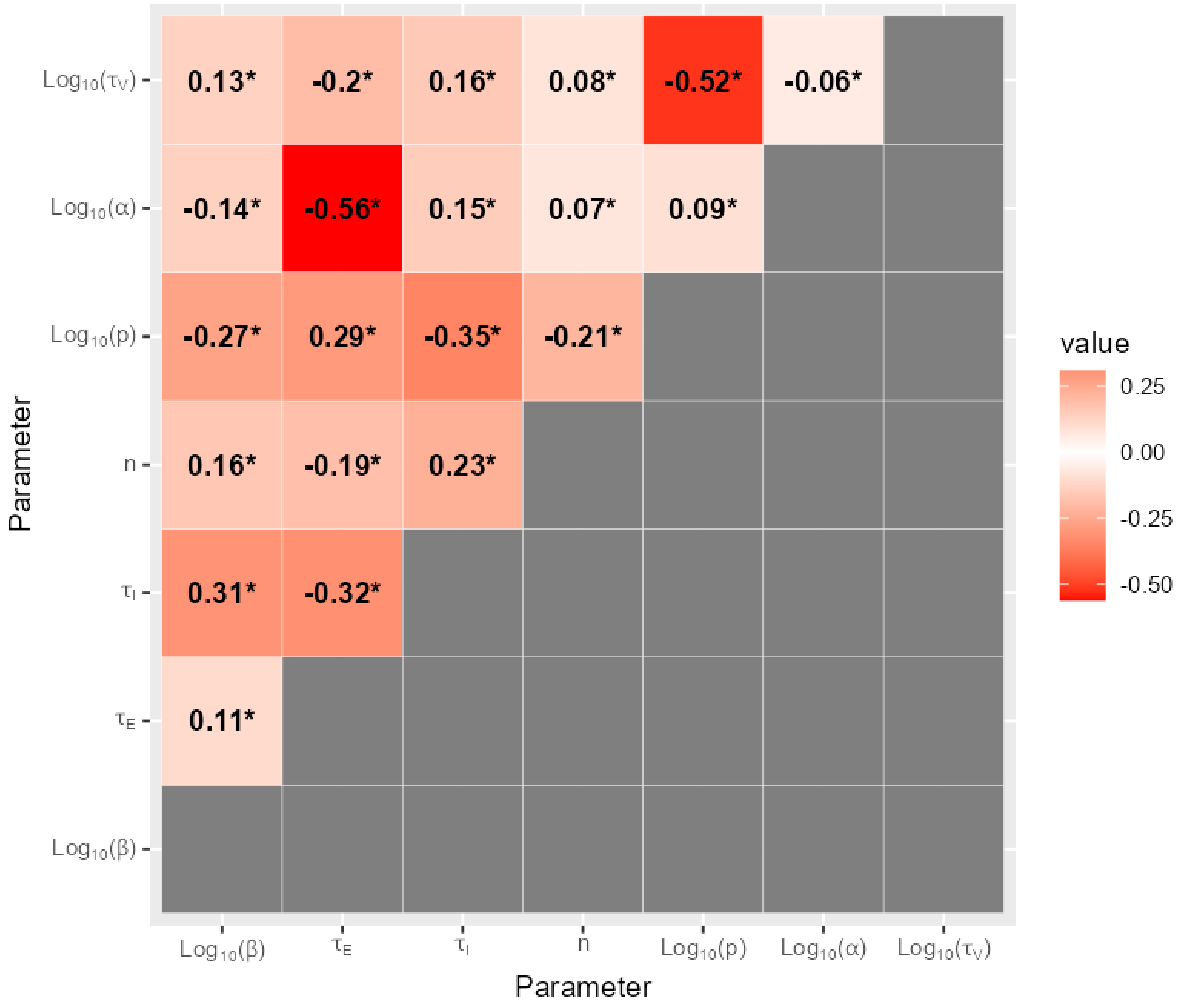

6.16 H5N8-2020 on DEF cells

Figure S50. The shifts in the distributions of model parameters during the ABC-SMC analysis of data from strain H5N8-2020 on DEF cells, when moving from the first particle population (containing accepted particles sampled from the prior distribution) to the posterior distribution via intermediate particle populations.

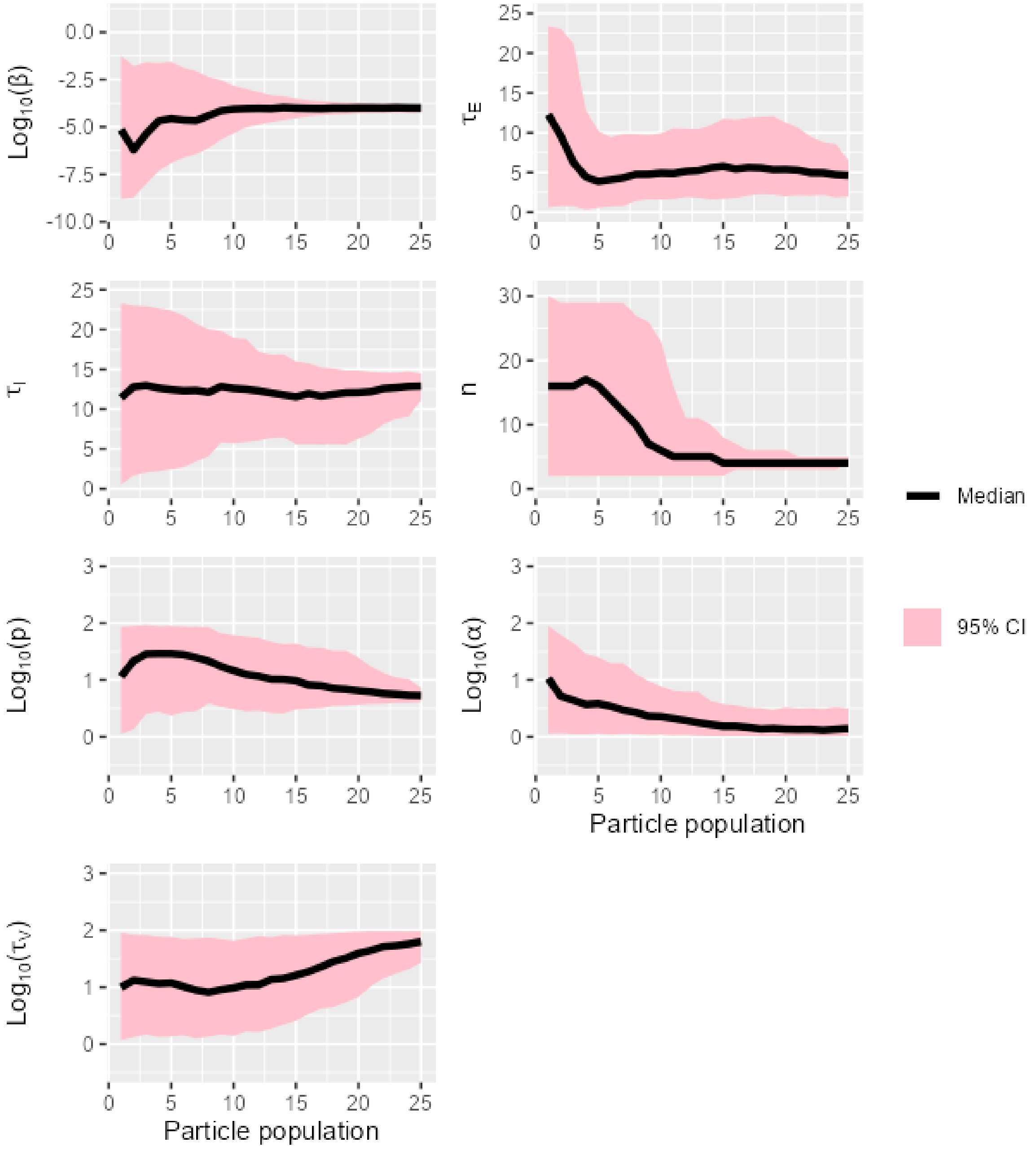

Figure S51. The overlap between prior and posterior distributions of model parameters for strain H5N8-2020 on DEF cells.

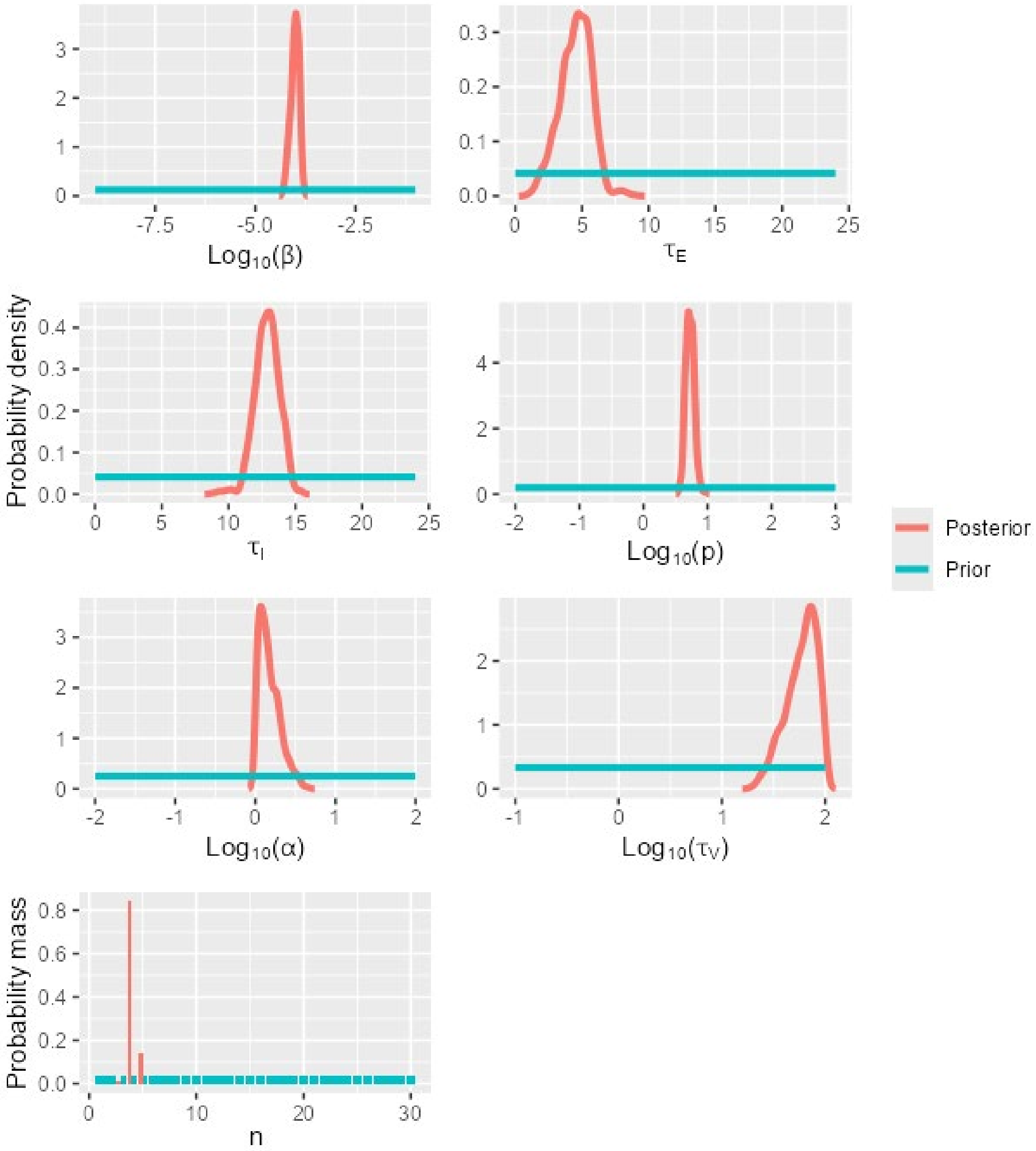

Figure S52. A matrix showing Kendall's rank correlation coefficient and the significance of the correlation (0.05 level, indicated by *) for pairs of model parameters calibrated using data for strain H5N8-2020 on DEF cells.

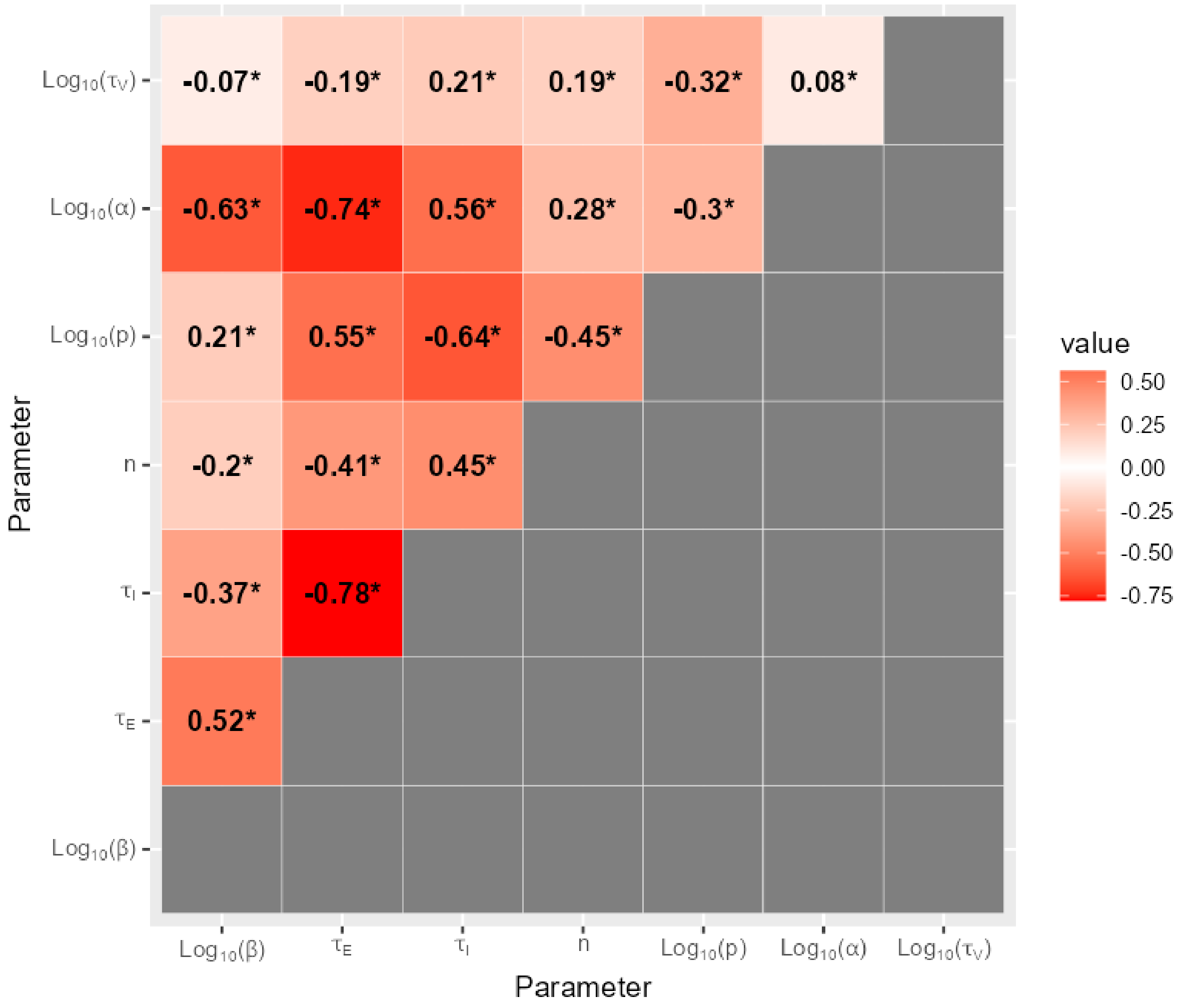
